# Supergene control of chiral development in mirror-image flowers

**DOI:** 10.1101/2025.08.29.672924

**Authors:** Haoran Xue, Marco Saltini, Nicola Illing, Kelly Shepherd, Olivia Page-Macdonald, Oliver Marketos, Caroline Robertson, Anand Shankar, Sarah Süß, Christian Kappel, Saleh Alseekh, Eva E. Deinum, Robert A. Ingle, Michael Lenhard

## Abstract

How genes determine the left-versus right-handed development of chiral structures is a fascinating question. The reciprocal placement of female and male organs on opposite sides of the midline in mirror-image flowers limits selfing and promotes efficient cross-pollination. Here we identify the molecular and developmental basis of floral handedness in butterfly lilies. Female and male organs deflect by a combination of genetically controlled chirality and gravitropism, orienting left and right with respect to an external rather than internal reference axis. Coordinated organ placement is controlled by a hemizygous supergene containing two causal loci, *MIR156-R* and *YUCCA-R*, responsible for opposite female and male organ orientation, respectively. This genomic architecture results in differential placement of the supergene alleles on the pollinators and maintenance of the reproductive polymorphism.

---

Left-right (LR) asymmetry is a fascinating example of biological pattern formation (*1*). Prominent examples include the LR asymmetric placement of internal organs in vertebrates and shell coiling in snails. In the latter, handedness is determined as early as the eight-cell stage, based on an intrinsic chirality of the actin cytoskeleton (*2–4*). By contrast, in vertebrates LR asymmetry appears to be determined only after the anterior-posterior and dorso-ventral axes have been established (*5–7*). In mice, the best understood case, the inherent chirality of microtubules in motile cilia that are aligned with the dorso-ventral and anterior-posterior axes generates a consistent left-ward flow of extracellular fluid across the node (*8, 9*). The mechanisms controlling handedness have been elucidated from rare mutants with aberrant organ placement. Except for snails, no genetically determined LR polymorphisms with coexisting left and right morphs have been found in natural populations. Thus, the effect of LR asymmetry genes on ecology and evolution remains poorly understood in animals.

The mirror-image flowers of enantiostylous plant species represent a naturally occurring LR polymorphism with demonstrated adaptive value (*10–12*). In mirror-image flowers, the style is either deflected to the left or right of the flower midline running along the dorsoventral axis, and often one or more stamens are deflected to the opposite side. In most enantiostylous species, floral handedness is not genetically determined and both left- and right-styled flowers can occur on the same individual, a condition known as monomorphic enantiostyly (*11, 13–15*). In some cases, however, floral handedness is genetically controlled, with plants forming consistently left-(L-morph) or right-styled (R-morph) flowers, a genetic polymorphism knowns as dimorphic enantiostyly (*12*). In *Heteranthera multiflora* flower handedness is determined by a single bi-allelic locus, with the right-styled allele dominant over the left-styled allele (*16*).

Mirror-image flowers represent an intriguing evolutionary solution to the challenge of avoiding self-pollination in hermaphroditic flowers while promoting efficient pollen transfer to conspecific individuals (*17*). While herkogamy, the spatial separation of male and female organs, can limit self-pollination, the reciprocal placement of sexual organs between different flowers enhances pollen transfer between them (*12, 18, 19*). As such, each flower form deposits pollen on a separate site on the pollinator’s body that matches the stigma position on the opposite flower form. Such reciprocal herkogamy is seen in heterostylous primroses and flax, where sexual organs are separated along the proximo-distal flower axis (*20*). The two morphs are determined by hemizygous *S*-locus supergenes, i.e. chromosomal segments harbouring separate causal genes for the different co-adapted traits that are inherited together because of suppressed recombination (*21–25*). Short-styled plants are heterozygous for the presence of the supergene, while long-styled ones are homozygous for its absence. Rare mutations in individual genes in the *S* locus result in homostylous individuals, with both anthers and stigma at the same position (*26, 27*). Whether mirror-image flowers have a similar genomic basis remains unclear.

The South African endemic genera *Wachendorfia* and *Barberetta* are perennial, insect-pollinated geophytes that flower once a year in spring and early summer. Their populations consist of L- and R-morphs, with the style and one of the three stamens deflected in opposite directions within a flower (*28–31*) (Figure 1A, B). In *W. paniculata* and *B. aurea*, most populations contain a roughly 1:1 ratio of L- and R-morphs (*28, 31*), and pollen-tracing experiments in *W. paniculata* have demonstrated predominantly disassortative pollen transfer between the morphs, with most pollen on a stigma originating from the anther of the opposing stamen in the other morph (*32*). This suggests that frequency-dependent selection based on disassortative pollination maintains a simple, single-locus polymorphism determining L- and R-morphs. The styles in *W. paniculata* deflect already in closed buds, possibly involving some rotation (*13*), yet the origin of handedness and the reference axes based on which it is defined are unknown. As such, *Wachendorfia* and *Barberetta* offer the opportunity to elucidate the genetic and molecular control of handedness and its effect on plant-pollinator interactions. Here we address the following questions: How does LR asymmetry develop in *Wachendorfia* and how is it oriented relative to the flower’s dorsoventral axis? How is handedness determined genetically and coordinated between styles and opposing stamens to ensure an integrated adaptive phenotype? Can the system break down to homostyly, with the style and otherwise opposing stamen on the same side of the flower?

**Fig. 1.**
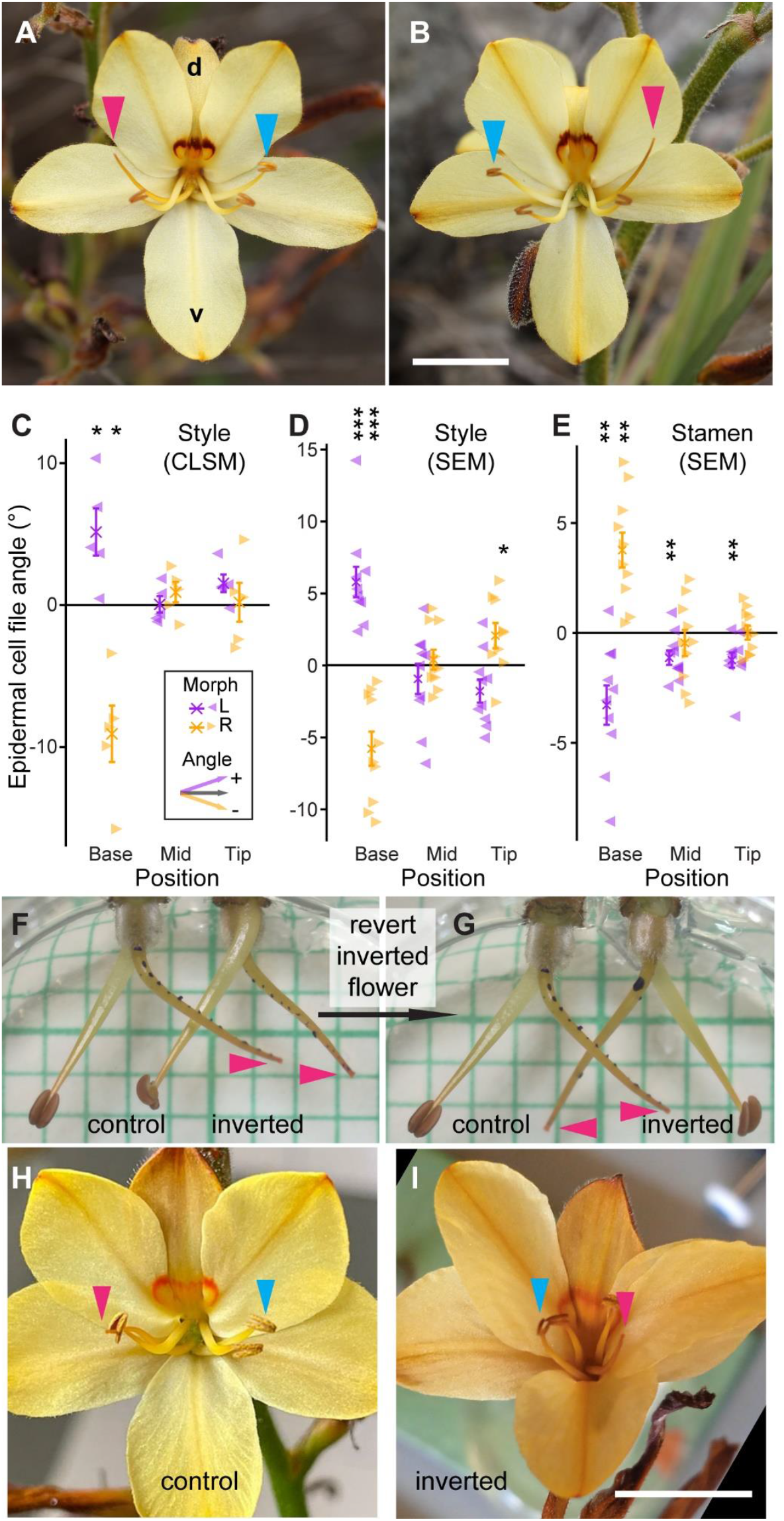
Styles of *Wachendorfia paniculata* rotate with respect to gravity. (**A, B**) Photographs of L-(A) and R-morph (B) flowers. Morph assignment is from the viewer’s perspective. Pink arrowheads indicate the stigma, blue arrowheads the anther of the opposing stamen. Dorsal (d) and ventral (v) sides are indicated. Scale bar is 10 mm. (**C**-**E**) Measurements of cell-file angles relative to long axis of the organs from styles (C, D) and stamen filaments (E) at the indicated positions based on from confocal (C) or environmental SEM images (D, E). Means and standard error of the mean are shown, as well as individual data points (arrowheads). Asterisks indicate significant differences from 0 at *p*<0.05 (*), *p*<0.01 (^**^), *p*<0.001 (^***^) as determined by one sample *t*-test. *n* = 5 per morph in (C) and *n* = 10 per morph in (D,E), except for R-morph tip in (D; *n* = 9). (**F**) Dissected R-morph buds after growth *in vitro* for 19 hours. Bud on the left was oriented as on an intact plant, with style above the opposing stamen, while the adjacent bud was inverted by 180° during culture. (**G**) As in (F), but with bud on the right reverted back to the natural orientation after the 19 hours in growth medium. Gridlines are 2 mm apart. (**H, I**) Open flowers on inflorescences cultured in the normal orientation (H) and inverted by 180° (I). Scale bar is 10 mm. Pink and blue arrowheads indicate styles and opposing anthers, respectively.

## Development of style deflection in *Wachendorfia*

Style orientation in *W. paniculata* is very developmentally robust. From 2448 inflorescences examined from 16 sites between 2022 and 2024, only a single inflorescence was found with one flower whose style orientation did not match that of the other flowers on the plant. This high stability supports genetic control of flower handedness.

To characterize the development of style deflection, we dissected unopened *W. paniculata* flower buds of different lengths and measured the alignment of the style and the opposing stamen with the flower midline (Figure S1A,B). Styles were straight in buds below 12 mm in length and became deflected from the midline in 12 to 15 mm long buds. Deflection became more pronounced in 16 to 18 mm long buds, reaching its maximum level in open flowers. These size classes correspond to their time to opening and are used below to stage buds as ‘early’ (below 12 mm long, three days to open), ‘mid’ (12 to 15 mm long, two days to open) and ‘late’ (16 to 18 mm long, one day to open). Stamen deflection broadly reflected the behavior of the style, but became pronounced only in late buds. Style deflection can also be visualized by time-lapse videos of dissected buds in culture medium (Movie S1). In some of these videos, the tip of the style does not merely follow an arc, but rather a spiral, suggesting that the style does not deflect simply by bending, but also in part by rotating.

To determine whether style deflection indeed involves rotation or twisting, we imaged the epidermis of the style at three positions spaced evenly from base to tip. Measuring the angles of epidermal cell files relative to the local long axis of the style indicated that cell files at the base were twisted in a right-handed helix in R-morph styles and in a left-handed helix in L-morph styles. By contrast, no consistent twist of cell files was visible in the middle or at the tip of the styles (Figure 1C,D; Figure S2A). Conversely, cell files in the stamen filaments had a twist in the opposite direction to that seen in styles (Figure 1E).

The cell-file twist suggests that there is an inherent left- or right-handed chirality in *W. paniculata* styles. However, to translate this chirality into macroscopic style deflection to the left or the right, it needs to be combined with polarity along two more axes in the flower, the proximo-distal axis of the style from the ovary to the stigma and the dorso-ventral axis of the flower. As *W. paniculata* flowers are arranged such that their intrinsic dorso-ventral axis and the gravity vector coincide, the style could be oriented with respect to the intrinsic dorso-ventral asymmetry or to gravity. To distinguish between these possibilities, we cultured dissected flower buds either in their natural orientation or rotated by 180°. When R-morph buds were used, styles deflected to the right, irrespective of the bud’s orientation (Figure 2F; Movie S2). As a result, the rotated flower was L-morph-like when returned it to its natural orientation (Figure 2G). The analogous effect was observed for L-morph buds, and overall 4/4 inverted L- and 7/7 inverted R-morph buds showed reorientation. To determine whether intact flowers also show this effect, we turned inflorescence stalks by 180° and cultured them until flowers opened. Of eight L- and R-morph inflorescences cultured in this upside-down position for one to three days, flowers opened on five and four inflorescences, respectively, and the majority of these flowers (13/14 for L-, 8/11 for R-morph) had styles deflected to the opposite side from their natural orientation (Figure 2H, I). These data indicate that *W. paniculata* styles and opposing stamens are oriented with respect to gravity.

**Fig. 2.**
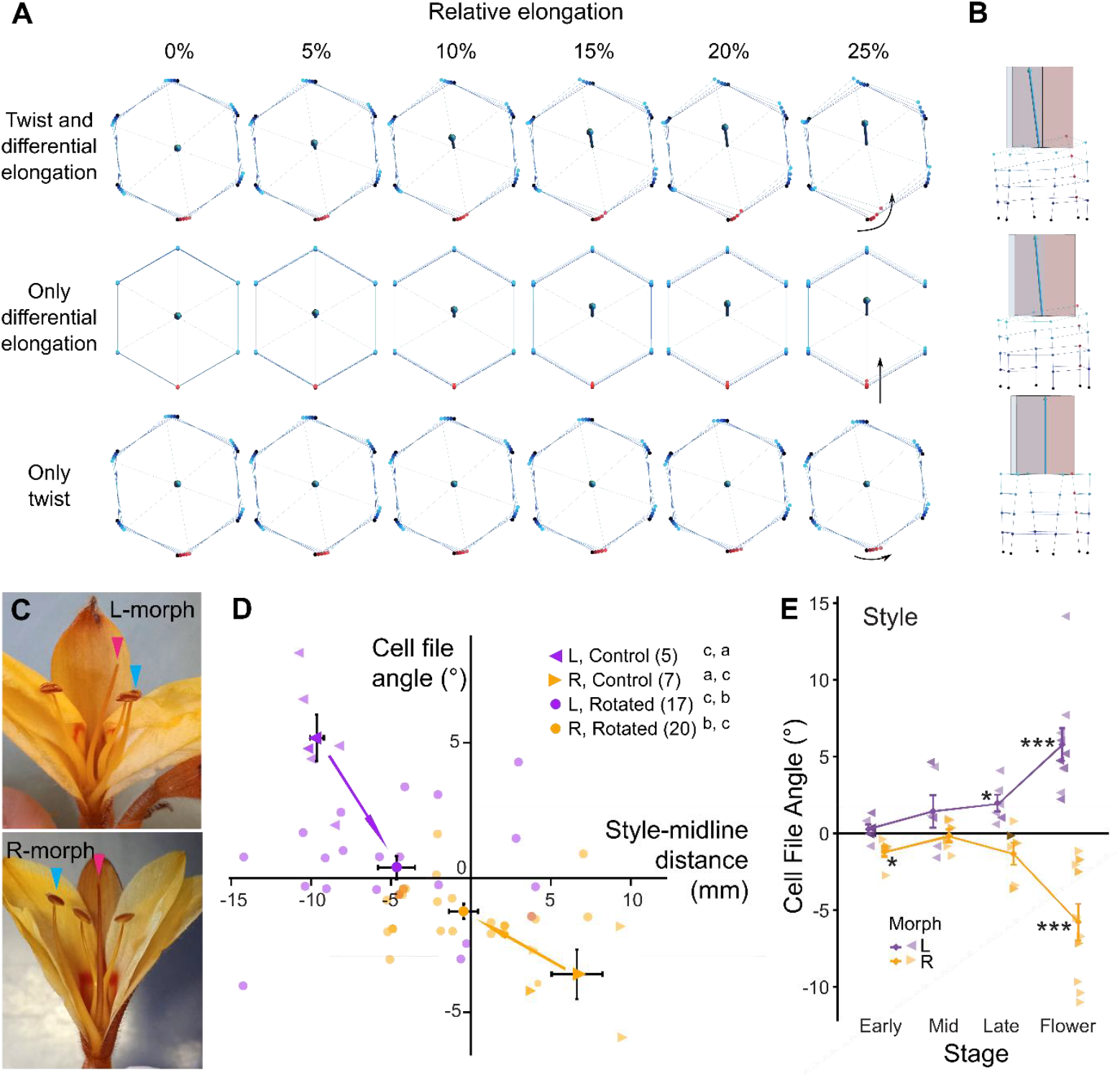
*Wachendorfia* style deflection results from twist and gravitropism-induced differential cell elongation. (**A, B**) Top view (A) and side view (B) of modelled styles under three scenarios from top to bottom: full model with differential elongation and twist, model with differential elongation only, and model with twist only. Vertices and edges are coloured by height, with lighter colours representing higher layers. (A) Sequential top views at increasing relative elongations. Black arrows show the motion of the (red) vertices that elongate the most in the differential elongation cases. (B) Side view. Red vertices connect the edges at the bottom of the style that undergo maximum elongation. (**C**) Straight styles resulting from bud rotation. Abaxial tepals have been removed for clarity. Pink and blue arrowheads indicate styles and opposing anthers, respectively. (**D**) Quantification of style-midline distance and cell file angle in styles from flowers without (control, arrowheads) and with rotation (rotated, circles) for 15 hours before opening. Transparent symbols show individual styles, filled symbols are means ± SEM. Sample sizes are given in brackets. Superscript letters indicate significantly different groups based on Tukey’s HSD after two-way ANOVA for style-midline distance (first letter) and cell-file angle (second letter). (**E**) Quantification of cell-file angles at the bases of styles from different stage buds as defined in Figure S1. Asterisks indicate significant differences from 0 at *p*<0.05 (*), *p*<0.01 (^**^), *p*<0.001 (^***^) as determined by one-sample *t*-test. *n* = 6 per morph for early, mid and late-stage buds and 10 per morph for flowers.

Gravitropic responses in plants are generally based on differential growth by cell expansion on the upper versus the lower side of an organ. Indeed, cells on the abaxial side were more than 20% longer and more than 30% wider than those on the adaxial side (Figure S2C,D). Thus, macroscopic style deflection appears to result from the combination of an intrinsic chirality in the style with a gravitropic response that causes stronger cell elongation on the basal side of the style.

## Style deflection by a gravitropic response combining with an intrinsic chirality

To test whether both twisting and differential growth are necessary and sufficient ingredients to produce the characteristic deflection of the *Wachendorfia* style, we built a simple biomechanical model. The framework consists of a bead-spring system, which lets us explore growth dynamics without the realism and computational burden of realistic organ-scale geometries (*15, 33*).

The style is represented as four identical hexagonal parallelepipeds stacked cap-to-cap. The choice of using four parallelepipeds is pragmatic: they are few enough for rapid computation, yet sufficient to show bending. In its starting state each segment has the same resting length, so the column is perfectly straight, mimicking an early developmental stage. We then imposed a right-handed twist and a gravitropically driven differential elongation, in which lower edges, i.e., those below the horizontal cross-section midline, lengthen more than those above.

We record both deflection (the angle between the final growth direction and the plane tangent to the axis of greatest elongation) and spin (indicating the accumulated angular deformation along the axis) after growth with different relative elongation (Figure 2A,B; Figure S3A,B). Deflection is maximal when twist and differential elongation act together. Unsurprisingly, spin appears only when twist is present. Differential elongation by itself simply bends the style upwards, while twist alone produces no deflection, instead yielding a straight style. Thus, the style only deflects away from the midline of the flower, i.e. moves out of the plane perpendicular to the basal hexagon running through its abaxial and adaxial vertices (bottom- and top-most vertices in Figure 2A), when both twist and differential elongation are combined (Movie S3); otherwise it stays within this plane (Movie S4, S5).

Because we model a right-handed twist, the style initially points left. Increasing the number of segments would likely reverse this, ultimately matching the rightward orientation observed empirically. How the plant terminates twisting at the appropriate length remains an open question worth investigating in future studies.

To test the prediction from the model that equal growth of the upper and lower sides of the styles should result in straight styles, we cultured intact flower buds on a clinostat with constant rotation, removing a consistent gravity stimulus. As predicted, these constantly rotated buds developed into flowers with styles and opposing anthers much closer to the midline (Figure S4), with many flowers forming essentially straight styles with minimal deflection (Figure 2C,D). Thus, the gravitropic response is a key contributor to macroscopic style deflection in *W. paniculata*.

We next asked whether the intrinsic chirality of the style results from a handedness of the epidermal cells driven by helical microtubules, as shown for mutants in *Arabidopsis thaliana* and other species with twisting growth (*34–37*). If so, twisting of the epidermal cell files should already be visible before the style begins to deflect, although it may be exaggerated during development; also, the straight styles produced by buds on the clinostat should still have some degree of twisted epidermal cell files. Neither prediction was met. Cell-file angles were only significantly different from 0 in late-stage buds (for L-morphs) and in open flowers (for both morphs), but not in younger stages (Figure 2E). In addition, we did not detect any significant cell-file twisting in the experimentally generated straight styles, suggesting that the epidermal cell-file twist follows, but does not drive style deflection (Figure 2D).

As a more direct test, we imaged cortical microtubules in epidermal and subepidermal cells of the style by anti-tubulin immunolabelling (Figure S5A-D). While helical microtubule arrangements were clearly visible in many cells, there was no consistent handedness to these within the styles of one morph and no difference between the morphs, neither in epidermal nor subepidermal cells (Figure S5E). Thus, these findings provide no evidence for a causal role of helical microtubules in epidermal or subepidermal cells for style orientation.

Our anecdotal observations of *Wachendorfia* plants in natural populations had suggested that style twisting is exaggerated when styles partially dry out, in a manner reminiscent of the increased helical coiling of awns in some Geraniaceae upon drying (*38, 39*). To investigate this in more detail, we dried L- and R-morph *W. thyrsiflora* styles from open flowers and buds of different stages. Indeed, dried styles showed increased twisting at the base, often resulting in coiling of the style (Figure S6A). In virtually all cases, the increased twisting (and coiling) had the same handedness as that of the epidermal cell files, i.e. left-handed in L-morph and right-handed in R-morph styles (Figure S6B,C). The drying-induced enhanced twisting and coiling was observed from mid-stage buds onwards and became more frequent in late-stage buds and flowers. This parallels the style movement away from the midline (Figure S1C), suggesting that the tissue-level chirality revealed by drying underlies style deflection.

## A hemizygous *R* locus supergene

The L:R morph ratio close to 1:1 in most natural *W. paniculata* populations suggests that flower handedness has a simple genetic basis with a single Mendelian locus. To identify morph-associated sequences, we Illumina-sequenced pooled DNA of 110 L- and 110 R-morph individuals from a single *W. paniculata* population and searched for morph-specific *k*-mers. Assembling reads containing such *k*-mers identified a sequence with similarity to *Arabidopsis thaliana YUCCA* (*YUC*) genes as an R-morph-specific gene candidate. To facilitate further analysis we established high-quality *W. paniculata, W. thyrsiflora* and *Barberetta aurea* genome assemblies using PacBio-sequencing of an R-morph individual from each species (Table S1). We also sequenced pooled DNA from L- and R-morph individuals from *W. brachyandra, W. multiflora, W. thyrsiflora* and *B. aurea*. Mapping the reads from the sequenced pools to the appropriate reference genome identified regions of between 100 and 200 kb that had a coverage ratio of L-versus R-morph reads close to 0 (Figure 3A; Figure S7A-F; Table S2). Inspecting the alignments in the corresponding regions indicated that the L-morph reads mapping to these regions were derived from repetitive sequences. The putative single-copy sequences were essentially not covered in the L-morph pools and had roughly half the average genome-wide coverage in the R-morph pools (Figure S8). To confirm that this genomic region is indeed absent from L-morphs, but present in all R-morphs, we PCR-genotyped the 220 *W. paniculata* individuals from the above pools. The morph-restricted *YUC* gene was present in all 110 R-morph plants and absent from all 110 L-morph plants (Figure 3B; Table S3). This was also confirmed in the other species using more limited genotyping (Table S3). Thus, flower handedness appears to be determined by a hemizygous region in *Wachendorfia* and *Barberetta*, with R-morph individuals carrying a chromosome with the extra region and L-morph individuals homozygous for the chromosome lacking it. We term this region the *R* locus.

**Fig. 3.**
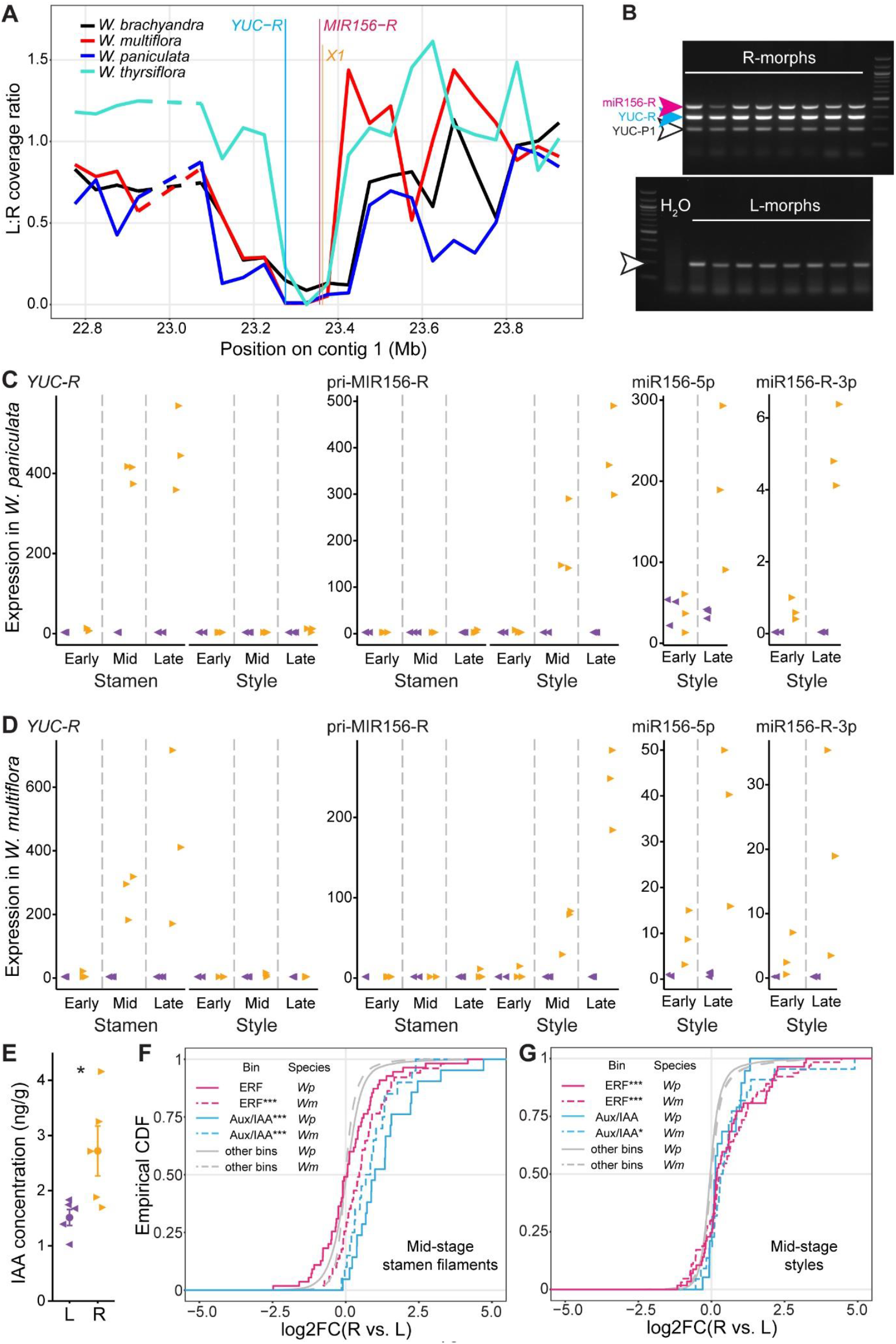
The *R* locus is a hemizygous supergene with three expressed genes in *Wachendorfia*. (**A**) Coverage ratio of whole-genome sequencing reads from L-versus R-morph pools of the four *Wachendorfia* species mapped against the *W. paniculata* reference assembly. (B) PCR genotyping of *W. paniculata* L- and R-morph individuals for the indicated targets. (**C, D**) Expression of the indicated genes and processed microRNAs in styles and opposing-stamen filaments of different-stage buds from *Wachendorfia paniculata* (C) and *W. multiflora* (D) based on RNA-seq and sRNA-seq, respectively. Expression is in normalized read count for mRNA/precursor and count per million (CPM) for miRNA. (**E**) Indole-3-acetic acid (IAA) concentration in filaments of the opposing stamens from L- and R-morph flowers of *W. thyrsiflora*. * indicates significant difference at *p*<0.05 (Wilcoxon Rank Sum test) (**F, G**) Empirical cumulative distribution function (Empirical CDF) plots of log2-fold changes from R-versus L-morph mid-stage stamen filaments (F) and styles (G) comparing the indicated MapMan categories to the genomic background (‘other bins’). Asterisks indicate significant differences from other bins at *p*<0.05 (*) and p<0.001 (^***^) based on Wilcoxon-Rank Sum test with Benjamini-Hochberg correction. *Wp* is *W. paniculata, Wm* is *W. multiflora*.

We annotated the reference genomes, taking into account transcriptome data from RNA-sequencing of styles and anther filaments (see below). This indicated that the hemizygous region contains two transcribed genes present across all three sequenced species, namely the *YUC* homologue, which we termed *YUC-R*, and a *MIR156* gene, termed *MIR156-R*, which gives rise to the microRNA *miR156-5p* (Figure S9A). In addition, the *Wachendorfia R* loci also contain a non-coding RNA gene with no homology to known genes (*Wp002WmWpStrST*.*819*, termed *X1* for simplicity). No related sequence was detected in the *B. aurea R* locus. The *R* loci contained a high fraction of repetitive elements (80.6% - 86.8%; Table S2), yet this was not significantly higher than in the surrounding chromosomal region (Figure S7G). The scaffolds carrying the *R* loci were largely syntenic between *W. paniculata, W. thyrsiflora* and *B. aurea* (Figure S10A), supporting their shared evolutionary origin. Thus, the R-morph phenotype is associated with a hemizygous region containing two conserved genes.

## Gene expression in deflecting styles and stamens

We next asked whether the genes at the *R* locus are expressed in the deflecting organs by performing RNA-seq on dissected styles and stamen filaments from early, mid and late-stage L- and R-morph buds of *W. paniculata* and *W. multiflora*. This analysis indicated that *YUC-R* was expressed specifically in stamen filaments of mid and late-stage R-morph buds, while pri-MIR156-R expression was restricted to the styles of mid and late-stage R-morph buds (Figure 3C,D). Expression of *X1* was seen in styles from early buds, but declined in mid- and late-stage buds (Figure S9B,C). Processing of the pri-MIR156-R stem-loop precursor (Figure S9A) generates the mature miR156-5p (derived from 5’ arm of precursor) and the miR156-R-3p molecules. Small-RNA sequencing from mid and late-stage styles showed a significant increase in miR156-5p levels in R-versus L-morph styles (Figure 3C,D). However, the *W. paniculata* L-morph styles also contained fully processed miR156-5p, likely derived from the other eleven *MIR156* genes outside of the *R* locus which give rise to identical miR156-5p sequences. In contrast, miR156-R-3p, which can be distinguished from the miR156-3p sequences derived from the other non-*R*-locus *MIR156* genes, was detected only in R-morph styles in both species, indicating that the pri-MIR156-R transcript is being processed into mature miR156-5p microRNA. By contrast, no sRNA derived from *X1* was detected. We confirmed the ability of the pri-MIR156-R transcripts from *W. paniculata* and *B. aurea* to give rise to mature miR156-5p by overexpressing the corresponding *MIR156-R* genes in *A. thaliana*. The transgenic lines for both constructs showed the previously described miR156-5p overexpression phenotypes with an extended juvenile phase, a higher total leaf number and smaller and more rounded leaves (Figure S11).

YUC enzymes catalyze the rate-limiting step in auxin biosynthesis from tryptophan. To test whether *YUC-R* expression leads to increased auxin levels, we measured IAA concentrations in stamen filaments dissected from mid and late-stage *W. thyrsiflora* buds. R-morph stamen filaments contained significantly higher auxin levels than those from L-morph buds, indicating that *YUC-R* encodes a functional YUC enzyme (Figure 3E). Additional evidence for increased auxin levels in stamen filaments of R-morph buds was provided by the specific enrichment of the auxin-related MapMan category Aux/IAA in the differentially expressed genes in mid-stage stamen filaments from both *W. paniculata* and *W. multiflora* (Figure 3F; Data S1). Analysis of the style samples indicated consistent enrichment of the ethylene-response factor (ERF) MapMan category in mid-stage styles of both species (Figure 3G; Data S1). We did not detect any downregulation of miR156-targeted *SQUAMOSA Promoter-Binding Protein-Like* (*SPL*) transcripts or other predicted target mRNAs in R-versus L-morph styles, nor any slicing of *SPL* transcripts by miR156-5p (Figure S12; Data S2), suggesting that miR156-5p mainly acts by translational repression in *Wachendorfia* styles (*40, 41*). Thus, together this suggests that *YUC-R* expression in mid-stage anther filaments causes their deflection to the left by increasing auxin levels. For styles, we consider *MIR156-R* as the more likely causal gene than *X1*, given their expression dynamics, the lack of any homology for *X1* and its absence from the *B. aurea R* locus despite the likely shared evolutionary origin of enantiostyly in *Wachendorfia* and *Barberetta* (*31, 42*).

## Homostyly resulting from deletions in the *R* locus supergene

Based on the above results, *MIR156-R* and *YUC-R* are strong candidates for determining the handedness of style and stamen deflection. As transformation of *Wachendorfia* is not possible at present, we searched for naturally occurring mutations in these genes. By analogy to derived homostyly in heterostylous taxa such as *Primula*, where mutations of individual genes within the *S-*locus supergene cause the stigma and anthers to occupy the same position in the flower (*22, 23, 27*), we predicted that null mutations in *MIR156R* should result in left-homostylous flowers, with both styles and opposing stamens deflected to the left. In contrast, *YUC-R* mutations should lead to right-homostylous flowers, with both these organs on the right of the midline. From surveying many populations of *Wachendorfia* in the Western Cape, we identified a single population containing right-homostylous plants of *W. multiflora*, two such populations for *W. paniculata*, and two populations of *W. brachyandra* with left-homostylous individuals (Figure 4A-F; Figure S13; Table S3). We performed whole-genome sequencing on the mutant individuals and L- and R-morph plants from the same populations (when present) and mapped the reads to the *W. paniculata* reference genome (Figure 4G). For the right-homostylous plants, we found four mutant haplotypes carrying full or partial deletions of *YUC-R*, but intact *MIR156* sequences. Two of these mutant haplotypes were from the same *W. multiflora* population at Redelinghuys, and the others were from two different *W. paniculata* populations. Two of the deletions affected the entire *YUC-R* transcribed sequence, while two only affected the C-terminal end, resulting in the predicted loss of a highly conserved region of the YUC-R protein (Figure 4H). Conversely, left-homostylous plants from the two *W. brachyandra* populations had intact *YUC-R* sequences, but lacked the *MIR156* and *X1* genes because of large deletions. These mutations were confirmed by Sanger sequencing of PCR products spanning the predicted deletion boundaries (Figure S14; Data S3), and co-segregation of the mutations with the homostylous phenotypes was shown by PCR genotyping (Table S3). Given the geographical origins of the different mutant haplotypes (Figure S13), we conclude that there were at least two independent primary mutations in both the *YUC-R* and *MIR156-R/X1* loci leading to homostylous phenotypes. However, due to the proximity of the populations and the extent of the deletions, we cannot exclude the possibility that the larger *YUC-R* deletions were secondarily derived from the initial shorter C-terminal deletions. Coverage analysis indicates that except for the single left-homostylous individual found in Rondebosch Common, all other homostylous samples from sites with many homostylous individuals were homozygous for the mutant *R* locus, suggesting their efficient selfing (Table S4). These results provide strong support that the *R*-locus supergene contains separate genes for stamen and style orientation. In R-morph plants, *YUC-R* causes stamen deflection to the left, and *MIR156-R* (or *X1*) cause style deflection to the right.

**Fig. 4.**
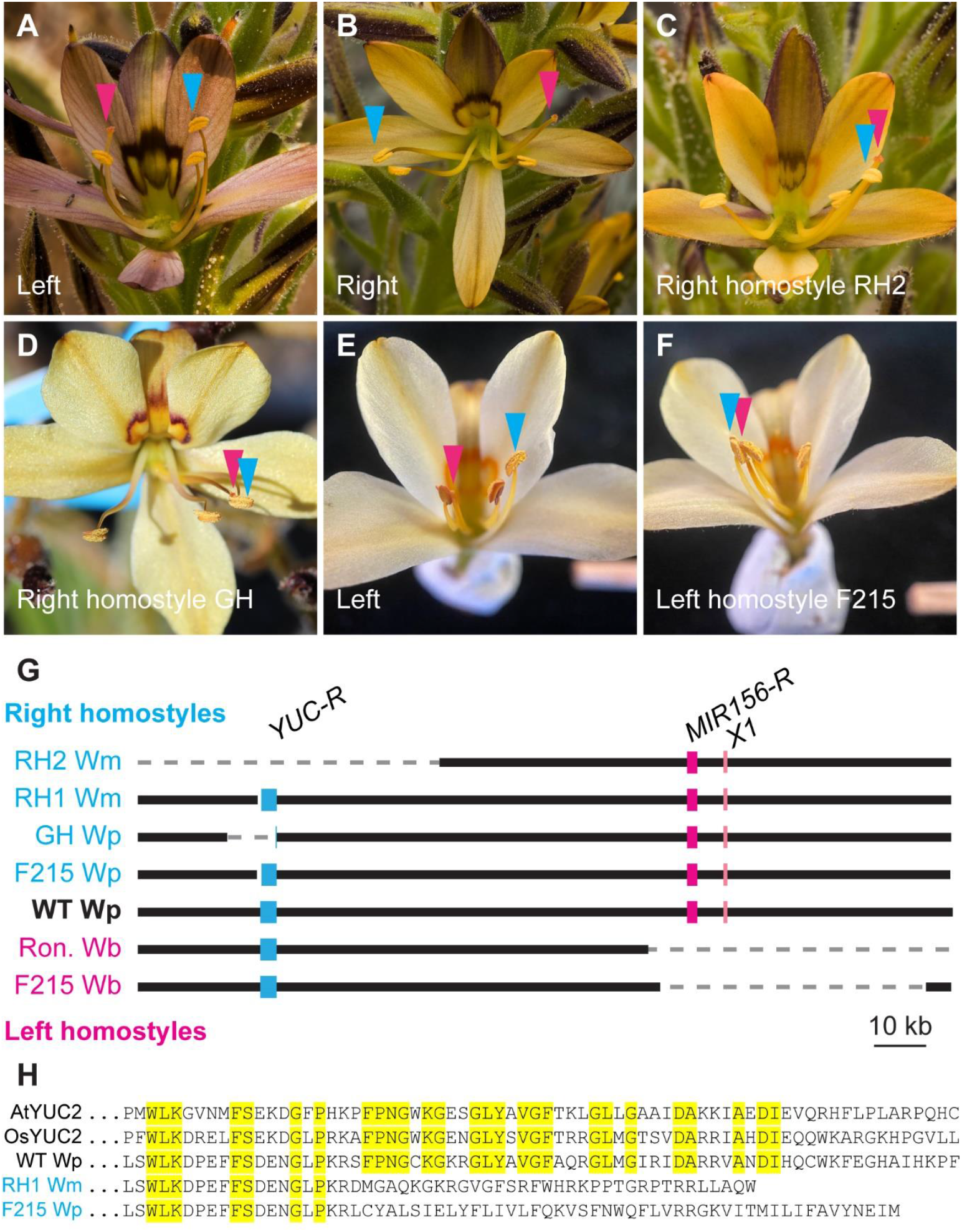
Mutations of *YUC-R* or *MIR156-R/X1* result in right or left homostyles, respectively. (**A**-**D**) Flowers of L- and R-morph of *W. multiflora* (Redelinghuys) compared to right-homostylous *W. multiflora* (Redelinghuys, RH) and *W. paniculata* (Groot Hagelkraal, GH). (E, F) Flowers of L-morph versus left-homostylous *W. brachyandra* (Farm 215). (**G**) Schematic representation of *R* loci from right- and left-homostylous mutants compared to wild type. Positions are with respect to the *W. paniculata R* locus. *Wm, W. multiflora*; *Wp, W. paniculata*; *Wb, W. brachyandra*; RH, Redelinghuys; GH, Groot Hagelkraal; F215, Farm 215; Ron., Rondebosch Common. (**H**) Alignments of predicted C-terminal YUC-R protein sequences from wild-type and the RH1 *W. multiflora* and F215 *W. paniculata* right homostyle mutants with YUC2 protein from *Arabidopsis thaliana* (AtYUC2) and rice (OsYUC2). Wild-type YUC-R sequence is from *W. paniculata*. Identical residues between the three wild-type proteins are shown in yellow.

## Evolution of the *R* locus

Dimorphic enantiostyly with genetically determined L- and R-morphs is thought to have evolved from monomorphic enantiostyly, where left- and right-styled flowers occur on the same plant (*11, 43*). *Dilatris ixioides*, a South African species from the Haemodoraceae that is closely related to the *Wachendorfia*/*Barberetta* clade displays monomorphic enantiostyly (*42*). To determine the possible origin of the *R*-locus supergene in *Wachendorfia*/*Barberetta*, we generated a reference genome assembly for *D. ixioides*, identified close homologues of *W. paniculata YUC-R* (see Methods for the criterion used) and performed phylogenetic analysis. The *D. ixioides* genome contains only one close homologue of *W. paniculata YUC-R*. The topology of the phylogenetic tree indicates that the gene duplication that gave rise to *YUC-R* only occurred after the divergence of *Dilatris* from the *Wachendorfia*/*Barberetta* clade, estimated at between 28 and 52 million years ago (Figure S15A,B). Genome-wide K_s_ analysis detected a recent whole-genome duplication in the *Wachendorfia*/*Barberetta* clade (Figure S15C), which seems to have given rise to the *YUC-P1* and *YUC-P2* paralogues, with *YUC-P2* subsequently lost in *B. aurea*. However, the *YUC-R*/*YUC-P* duplication is significantly older than this, but appears younger than the monocot τ whole-genome duplication (Figure S15C). Synteny analyses indicated large-scale homology between scaffolds carrying most closely related *YUC* paralogue across the four sequenced genomes (Figure S10B). These also carried the only non-*R*-locus *MIR156* gene with an extended sequence match to *MIR156-R* (55 bp including miR156-5p in *W. paniculata*), located 2 Mb from the *YUC* paralogue. Comparing the *R*-locus containing scaffolds with those carrying the *YUC-P* paralogues and related *MIR156* genes suggested that the *R*-locus scaffold resulted from a large-scale segmental duplication after divergence from *Dilatris* (Figures S10C,D), followed by structural rearrangements to bring *YUC-R* and *MIR156-R* close together and likely their neofunctionalization for controlling enantiostyly.

Hemizygous genes are expected to be under less efficient selection than diploid genes because of their reduced effective population size (*44, 45*). As predicted, the *YUC-R* clade showed a significantly higher dN/dS ratio (0.55 [95% confidence interval 0.40-0.73]) than the *YUC-P* clade (0.16 [95% confidence interval 0.08-0.28]), supporting our interpretation of the *R* locus as a hemizygous region under long-term negative-frequency dependent selection.

## Discussion

Although the mirror-image flowers of *Wachendorfia* were first described almost 140 years ago, it has remained unclear how floral handedness develops and how it is genetically controlled (*46*). The functional importance of these mirror-image flowers in promoting efficient cross-pollination results from the reciprocal pollen placement on, and pick-up from, the left or right wings of pollinators by the two morphs (*10, 32, 47*). Here we have identified the developmental and molecular basis of this ecologically important, genetically controlled left-right asymmetry. The development of the mirror-image flowers in *Wachendorfia* requires three processes: growth of the style and opposing stamen along the proximo-distal axis of the flower, a gravitropic response leading to unequal growth on the lower versus upper side of the organs, and a genetically encoded intrinsic chirality. Thus, in contrast to organ placement of vertebrates (*5–7*), LR asymmetry in *Wachendorfia* is defined relative to an asymmetric external input, rather than to internally determined axes. The intrinsic chirality of the style does not appear to result from a helical arrangement of cortical microtubules in epidermal and subepidermal cells, as observed in many mutants with organ twisting (*34–37*). Rather, based on their drying-induced enhanced twisting and coiling, we suggest that it is due to a helical structure or tissue interaction inside the styles, similar to the coiling awns of Geraniaeceae (*38, 39*).

The intrinsic chirality of the organs is flipped by the presence of the *R-*locus supergene (Figure S16). Expression of *MIR156-R* in developing styles leads to a right-handed twist of the style, while *YUC-R* expression in stamen filaments causes a left-handed twist. The coordinated change in style and stamen deflection between the morphs - essential for promoting cross-pollination - results from a hemizygous supergene architecture as seen also for other floral polymorphisms (*22, 23, 48*), leading to co-inheritance of two independent handedness-reversing genes. Similar to genes on Y-chromosomes, *YUC-R* appears to be under less efficient purifying selection than its paralogues. Although it is unclear when *YUC-R* became part of the hemizygous supergene after its origin by duplication, the considerable difference in dN/dS ratios suggests that this hemizygous polymorphism has been maintained for a long time, supporting a model of negative frequency-dependent selection mediated by pollinators. This is in line with the roughly 1:1 ratio of the morphs observed in most *W. paniculata* populations (*28*). At the same time, enantiostyly can also break down to homostyly, with both style and the normally opposing stamen on the same side of the midline, by mutations in either of the two handedness genes. This is expected to lead to more efficient self-pollination and may provide reproductive assurance in pollinator-limited regions.

Dimorphic enantiostyly with genetic control of handedness in *Wachendorfia* and *Barberetta* likely evolved from monomorphic enantiostyly as seen in *Dilatris* (*11*). Similar to the evolution of separate male and female sexes (*49*), this process of genetically fixing and canalizing a variable phenotype may have occurred via two evolutionary steps (*43, 50*). First, floral handedness was fixed to that of an L-morph, before second, the evolution of the *R* locus established the R-morph and the extant polymorphism. Identifying the molecular changes underlying the first step remains an open challenge.

What is the ultimate source of chirality in *Wachendorfia* and *Barberetta* flowers? The intrinsic chirality of cellulose microfibrils is a potential candidate. Individual microfibrils form right-handed helices in isolation and right-handed microfibril bundles are seen in mutants with reduced pectin levels in their cell walls (*51–53*). Right-handed cellulose microfibril helices result in left-handed cell-file and tissue twist, similar to what is seen in L-morph styles (*37, 54*). This seems consistent with left style deflection being the homozygous recessive trait that develops in the absence of the *R*-locus supergene and that may be overwritten by *MIR156-R* activity. We note that ERF activity has been directly linked to pectin modification (*55, 56*). Thus, the increased expression of *ERF* transcription factors in R-morph styles may modify pectin levels or structure and thus modulate the properties of cellulose microfibrils to switch the intrinsic chirality of the organ.

In summary, our study into the molecular basis for a naturally occurring LR polymorphism underlines the importance of hemizygous supergenes in controlling separate morphs in plants. We identify genes that can reverse an intrinsic chirality, and a novel mechanism for determining left and right with reference to an external input. Combined with the results from pollen tracing (*32*), our work suggests that the activity of the *R* locus results in spatially separate placement of the dominant and recessive haplotypes on pollinators’ bodies, leading to their preferential transfer within populations and long-term maintenance of an iconic floral polymorphism by negative-frequency dependent selection.

## Acknowledgments

We thank Spencer Barrett for critical discussion and advice on enantiostyly and plant reproductive systems, Alice Fairnie, Sam McCarren and Bruce Anderson for help with field work and critical discussion, Otto Baumann, René Schneider, Anh Minh Do, Dongbo Shi, Janica Mej Theron, Dirk Lang, Emma Bender, Ralph Gräf and Marianne Grafe for help with microscopy and image analysis, Mathias Scharmann for advice on genomic analyses, Silvia Vignolini for discussion, the Botanical Garden at University of Potsdam for providing material, and all landowners and institutions for providing permissions to sample.

## Funding

Human Frontier Science Program grant number RGP0036/2021 (NI, EED, ML) Harry Crossley Research Fellowship (CR)

## Author contributions

Conceptualization: HX, MS, NI, EED, RAI, ML

Methodology: MS

Investigation: HX, MS, NI, KS, OPM, OM, CR, AS, SS, CK, SA, EED, RAI, ML

Formal analysis: HX, MS, NI, CK, ML

Visualization: HX, MS, NI, ML

Funding acquisition: NI, EED, ML

Supervision: NI, EED, RAI, ML

Writing – original draft: HX, MS, NI, ML

Writing – review & editing: HX, MS, NI, KS, OPM, CR, EED, RAI, ML

## Competing interests

Authors declare that they have no competing interests.

## Data and materials availability

All sequencing data and whole-genome assemblies will be made available in NCBI under BioProject PRJNA1138953. All other data are available in the main text or the supplementary materials.

## Supplementary Materials

Materials and Methods

Figs. S1 to S16

Tables S1 to S6

References(*57–93*)

Movies S1 to S5

Data S1 to S6

## Notes

### Competing Interest Statement

The authors have declared no competing interest.

