## Supplemental Information for "Supergene control of chiral development in mirror-image flowers"

### Materials and Methods

#### Plant material, populations, permits

Plant material for genetic analyses are summarized in Table S6 below. W. paniculata infloresences from sites on the Cape Peninsula were used for measurement of cell file angles during bud development, and the gravity experiments. In addition to plants from natural populations listed in Table S6, we also used W. thyrsiflora L- and R-morph plants cultivated in the Botanical Garden at the University of Potsdam.

#### Phenotyping, dissections

W. paniculata inflorescences were picked in the field and stored in water in the laboratory. Developmental stages of W. paniculata buds were defined by measuring the length of buds, and then dissecting the buds by removing the abaxial tepals. The abaxial tepals were also removed from flowers to image style and stamen deflection. The number of days it took buds to flower was measured by tagging the pedicel at the base of buds, and noting the time it took them to open.

#### Light, confocal and scanning electron microscopy, measurements of cell file angles, lengths and widths

*Light microscopy*

Photographs of dissected buds or flowers were taken with an iPhone 11 fitted with a 10X macrolens. Images were analysed in ImageJ, with style and stamen deflection measured as an angle from the midline of the bud or flower.

*Confocal microscopy*

Whole pistils were fixed for 1 hour at room temperature using 4% (w/v) paraformaldehyde prepared in 1X phosphate buffered saline (136.89 mM NaCl, 2.68 mM KCl, 5.37 mM Na2HPO4, 1.76 mM KH2PO4; pH7.4). Fixed samples were washed three times in PBS, followed by a final rinse in dH2O. Endogenous pigments were removed using ClearSee (10% (w/v) Xylitol; 5% (w/v) sodium deoxycholate and 25% (w/v) Urea in dH2O; (57)). Fixed pistils were incubated in ClearSee for 8-10 weeks with the solution replaced three times per week. Cleared pistils were rinsed in dH2O for 1 hour before overnight staining at room temperature with calcofluor white (Sigma, product no. 18909). Samples were destained overnight with dH2O. All staining and destaining steps were performed in the dark with gentle agitation. Samples were placed on their adaxial side on long coverslips (25 × 50 mm) in water for imaging using a Zeiss LSM 880 Confocal. Detailed z-stacks were taken at three, evenly spaced positions along the style: base, mid and tip. The sample was then flipped onto its abaxial side using tweezers and imaging repeated. Z-stacks of styles were converted to 3D projections using ZEN-lite software. A central line of cells was selected and the angle between the cell file relative to the edge of the style calculated using ImageJ. Two-sample t-tests were performed to test for a significant difference between cell file angles in left- versus right-handed samples.

*Environmental scanning electron microscopy*

Environmental scanning electron microscope (eSEM) images of dissected stamens and styles were captured on a ThermoFisher Apreo Serial Block Face FESEM. For measurement of cell file angles, stamens and styles were imaged at the base, at the tip and half way along the length of the style/stamen. For measurement of cell lengths and breadth, the base of the style was imaged, above the ovary, from abaxial and adaxial viewpoints. Cell file angles were measured in Image J by measuring the relative angles at three points- and subtracting the angle taken at the middle of the style or stamen, from the average of the angles of the two outer surfaces. The length and breadth of 5 adjacent cells at the midline of the abaxial and adaxial view were measured in Image J, with three measurements per sample.

*In vitro* experiments: upside down flowers, upside down inflorescences, rotating flower buds

Whole inflorescences, flowers and dissected floral organs of W. paniculata can be maintained on ½ strength Murashige & Skoog (MS) agar (2.165 g MS basal medium, 7 g agar in 1 L distilled water, autoclaved and cooled to 60°C before pouring). To assess whether W. paniculata styles are oriented with respect to gravity, pairs of mid-stage buds were selected from a single inflorescence. From each ½ MS agar plate, a 3X3 cm block of agar was removed to create a wall of agar into which dissected organs could be inserted. Buds were cut from the inflorescence using a scalpel blade, preserving a 0.5 cm pedicel. All petals and the adaxial stamens were removed and the pedicel pressed into the exposed wall of agar. The removal of a block of agar prevented the dissected organs from scraping against the agar during development. One bud was inserted in the natural orientation, the other "upside down" (rotated 180 degrees around the pedicel axis). The dissected organs were imaged using a dissecting microscope (Olympus SZ61) fitted with a Zeiss Axiocam 208 colour camera. Imaging was repeated after 12-24 hrs.

Whole inflorescences were also subjected to the inversion treatment. Inflorescences were maintained in sterile universals filled with ½ MS agar. The mouth of each universal was sealed with parafilm and the entire inflorescence hung upside down from a retort stand. Flowers were imaged as they opened over the course of 1-3 days.

To investigate the development of Wachendorfia flowers in the absence of a gravitational input, W. paniculata buds were rotated on a clinostat. ½ MS agar was poured into 1.5 mL microfuge tubes. Mid-stage W. paniculata buds were cut from the inflorescence using a scalpel blade and the pedicels inserted into the agar. Tubes were sealed with parafilm, and the buds rotated on the clinostat for 1-2 days, or until opening.

Immunolocalization

The immunolocalization largely followed Du et al. 2021 (58). Whole pistils were put into freshly prepared fixative (4% (w/v) paraformaldehyde, 0.5% (w/v) glutaraldehyde, 0.3% (v/v) Tween-20, 0.3% (v/v) Triton X-100) in microtubule stabilization buffer (MTSB) (PIPES 15.12 g/L, $MgSO_4 \cdot 7H_2O$ 1.24 g/L, EGTA 1.90 g/L, pH = 6.9) and were vacuum infiltrated at -0.075 MPa (550 mm Hg) for 10 min three times, followed by an additional 3 h fixation at room temperature. After removing the fixative, the samples were incubated in 10%, 20%, and 30% sucrose in MTSB, for 20 minutes each. The fixed pistils were stored in MTSB at 4°C until embedding in agarose gel.

The pistils were embedded in 7% low-melting agarose gel and cut into trapezoidal prisms to retain information about their orientation. If the style was not flatly embedded in the agarose gel, it was divided into a maximum of four parts. The bottom of each agarose block was cut parallel to the embedded part of the style, enabling longitudinal sectioning of each part. Sectioning was performed using a vibratome (VT1000 S, Leica Mikrosysteme, Wetzlar, Germany) at speed of 0.65 mm/s, sectioning frequency of 40 Hz, and cutting thicknesses of 170 μm. The sections were stored in MTSB in a 24 deep-well plate at 4°C.

For cell permeabilization, MTSB wsas replaced by 200 µL enzymatic solution containing 2% driselase (w/v) and 1% (v/v) Triton X-100 in MTSB and incubated at room temperature for 25 min. The samples were washed 3 times with TBS (8.8 g/L NaCl, 20 mM Tris-Hcl, pH = 8.0) for 5 min and washed with methanol stored at −20°C. After removing the methanol, the sections were incubated with 200 µL primary antibody solution, which contained 1% bovine serum albumin (BSA) and either monoclonal mouse-α-tubulin antibody DM1A (Sigma-Aldrich, St. Louis, USA) with 1:500 dilution or monoclonal mouse anti-β-tubulin antibody E7 (Developmental Studies Hybridoma Bank, Iowa City, USA) with 1:200 dilution in TBS. The samples were incubated at 4°C with shaking for 20 h. The primary antibody solution was removed and the samples were washed 3 times with TBS for 10 min. Then the sections were incubated with 200 µL secondary antibody solution, which contained Alexa Fluor 488 conjugated donkey anti-mouse IgG (ThermoFisher Scientific, Waltham, USA) at a dilution of 1:500 and 1% BSA in TBS. Incubation was carried out for 2.5 h in the dark at 37°C.

The sectioned were positioned on microscopic slides, embedded in Fluoromount-G, then covered with high-precision coverslips of 170 µm thickness. Following sealing with nail polish, the slides were stored at 4°C until confocal microscopy was performed. A laser scanning microscope (LSM 710, Zeiss, Oberkochen, Germany) equipped with a 40x water immersion objective was used to capture z-stack images from various regions of the style. To detect signal from Alexa Fluor 488, a 488 nm laser line (power: 30 mW) is used for excitation and emission from 503 nm to 572 nm was collected. The pinhole size was 1.0 AU (34.5 µm).

The image analysis was conducted using ImageJ. Microtubule arrays each cell were assigned into four class: left-handed helix, right-handed helix, parallel, and disarray. Chi-square tests were conducted to compare proportions of different classes between left- and right-morph plants.

### Modelling

Our goal was to test the joint roles of axial twist and gravitropically driven differential growth in generating the characteristic lateral deflection of the *Wachendorfia* style. To do so we chose the simplest mechanical representation that (i) permits independent control over twisting and growth, (ii) remains computationally lightweight for parameter sweeps, and (iii) is transparent enough to highlight causal mechanisms without excessive geometric detail. A bead-spring stack of four identical, hexagonal parallelepipeds fulfilled these criteria.

The position of the bead $(j,k)$ is $\mathbf{r}_{j,k} = (x_{j,k}, y_{j,k}, z_{j,k})$ with layer index $j$=1,...,4 and vertex index $k$=0,...,5. In the initial configuration the stack is aligned with the $z$-axis and its centroid lies at $(0,0,z)$. Axial springs of rest length $l_L$ connect corresponding beads in adjacent layers, while transverse springs of rest length $l_T$ connect neighbouring beads within each hexagon. A constant twist α is imposed between successive layers.

Growth is introduced as a perturbation by modifying the rest lengths of specific springs, as in (*15*).

Differential elongation is implemented such that springs attached to beads lower in the gravitational field (i.e., for negative y-values) experience greater rest length increase. Specifically,

$$\Delta L = \Delta_{\mathsf{max}} \frac{-y_{j,k}}{\sqrt{x_{j,k}^2 + y_{j,k}^2}} \Theta\left(-y_{j,k}\right), \qquad (1)$$

where Θ is the Heaviside step function. Growth proceeds layer-by-layer: first $j$=1, then $j$=1,2, and so on.

To retain analytical tractability we restrict bead motion to the line joining paired nodes in consecutive layers, reducing the dynamics to the *z* component. The equation of motion for bead (*j*,*k*) is therefore

$$\begin{aligned}\ddot{z}_{j,k} = &-k_L\left(1-\frac{l_L+\Delta L}{L_{j,k}}\right)(z_{j,k}-z_{j-1,k}) - k_L\left(1-\frac{l_L+\Delta L}{L_{j+1,k}}\right)(z_{j,k}-z_{j+1,k}) \\ &- k_T\left(1-\frac{l_T}{T_{j,k}}\right)(z_{j,k}-z_{j,k-1}) - k_T\left(1-\frac{l_T}{T_{j,k+1}}\right)(z_{j,k}-z_{j,k+1}) - c_{\text{damp}}\dot{z}_{j,k},\end{aligned} \quad (2)$$

where $L_{j,k} = |\mathbf{r}_{j,k} - \mathbf{r}_{j-1,k}\}|$ and $T_{j,k} = |\mathbf{r}_{j,k} - \mathbf{r}_{j,k-1}|$ are the distance between adjacent beads expressed as function of *z* only, and $c_{\text{damp}}$ is a damping constant to ensure small oscillations converge to their equilibrium positions. Periodic indexing applies (i.e., $k$=0 ≡ 6).

The system of ODEs was solved following the procedure described by (*15*). Implementation details are provided in our Mathematica notebook (Data S6). The configuration shown in Figure 2A,B and Movies S3–S5 corresponds to the final mechanical equilibrium of the system.

### DNA extraction, Illumina and PacBio sequencing

Mature leaves from 110 left and 110 right morph individuals of W. paniculata were sampled into individual tea bags and dried down in silica gel. DNA was extracted from dried leaf material using a modified CTAB-protocol (59) (https://figshare.com/s/0bc4d41c8c3cb1adb61e). DNA from all individuals of the same morph from one locality was pooled, purified again using AMPure beads, before shipment to Novogene for short-read sequencing.
Flower buds were harvested from 50 L- and 50 R-morph individuals for W. brachyandra, W. thyrsiflora and W. multiflora and frozen in liquid nitrogen on the day of harvest and stored at -80°C. Buds were also harvested from 40 L- and 40 R-morph B. aurea individuals and dried in silica gel. Two buds from each individual (60 mg) were ground to a fine powder under liquid nitrogen using a pestle and mortar. The powder was added to 450 µl DNA extraction buffer (1.4 M KCl, 0.02 M EDTA, 0.1 M Tris pH 8, CTAB (2% (w/v), β-mercaptoethanol (0.5% (v/v), PEG 20,000 4% (w/v), and incubated at 65°C for 15 min before further purification with chloroform as described (59). DNA from all individuals of the same morph in each species was pooled before shipping to Novogene Europe for short-read sequencing.

Buds were also harvested from homostylous individuals of W. brachyandra, W. multiflora, W. paniculata and left and right morph individuals from the same populations (when present), and dried in silica gel. Two buds from each individual (60 mg) were ground to a fine powder under liquid nitrogen using a pestle and mortar. The powder was added to 450 µl DNA extraction buffer (1.4 M KCl, 0.02 M EDTA, 0.1 M Tris pH 8, CTAB (2% (w/v), β-mercaptoethanol (0.5% (v/v), PEG 20,000 4% (w/v), and incubated at 65°C for 15 min before further purification with chloroform as described (59).

Genomic DNA for PacBio sequencing was isolated from 1 g of floral buds from a right morph W. paniculata, W. thyrsiflora and B. aurea individual. Floral buds were ground to a fine powder with a pestle and mortar under liquid nitrogen, whereafter the frozen powder was resuspended in 5 ml high salt buffer I (50 mM EDTA, 0.16 M NaCl), and incubated at 50°C for 10 min. The samples were centrifuged in a JA14 rotor at 10,000 g for 10 min at room temperature, whereafter the pellet was resuspended in 5 ml H1 Lysis buffer from the Macherey-Nagel™ NucleoBond™ HMW DNA kit. Genomic DNA was extracted according to the manufacturer's instructions.

### RNA extraction, RNA-seq, small RNA-seq

For RNA-seq and small RNA-seq, stamen samples consisted of the filament of the abaxial stamen, while style samples were dissected from above the ovary. Wet tissue (10-60 mg) was pooled into one biological sample, frozen in liquid nitrogen and stored at -80°C until RNA extraction. Total RNA was isolated using a modified protocol from (60). Any contaminating DNA was removed from RNA samples using the RNA clean and concentrator kit (Zymo Research). RNA samples were treated with GenTegraRNA (NBS Scientific, USA) according to manufacturer's instructions to allow for shipping at ambient temperature. For RNA-seq, stamen and style samples were collected from early, mid, and late buds of left and right morph individuals of W. paniculata and W. multiflora (2 x 3 x 2 x 2 = 24 types of sample in total). For each type of sample, three biological replicates were collected. Adapter and quality trimming were conducted using Trim Galore version 0.6.10.

Genome assembly and annotation, identification of R locus, analysis (RepeatMasker, synteny plots)

We generated genome assemblies of W. paniculata, W. thyrsiflora, B. aurea, and D. ixioides using hifiasm version 0.19.0-r534 (61) and we used default settings with only “--hg-size” (estimated haploid genome size) specified. We estimated contiguity and completeness of the assemblies using QUAST version 5.2.0 (62) and BUSCO version 5.4.4 (63). The database chosen for the BUSCO analysis was liliopsida_odb10. The “-m genome” and “--augustus” options were specified. We used RepeatModeler (64) and RepeatMasker (65) to annotate the repeats in the assemblies. Then we conducted the structural annotation of the repeat masked genomes using BRAKER3 (66), with the partition Viridiplantae of the protein database OrthoDB v11 (67) as protein evidence. For W. paniculata, we aligned the trimmed style and stamen RNA-seq reads to the genome assemblies with HISAT2 version 2.2.1 (68), and used the aligned reads as RNA evidence.

We used BWA-MEM version 0.5.5 (69) to align Illumina DNA-seq reads to the genome assemblies of W. paniculata, W. thyrsiflora and B. aurea, then used SAMtools version 1.3.1 (70) to sort and index alignment files. We performed coverage analyses with the pool-seq data of W. brachyandra, W. multiflora, W. paniculata, W. thyrsiflora, and B. aurea. We removed reads mapped to the annotated repeats with SAMtools version 1.3.1. Then we calculated read coverage of each pool over 50-kb windows using mosdepth version 0.3.3 (71) and coverage of each window was normalized by dividing by the genome-wide average read coverage of each. The ratio of normalized coverage for left versus right pools was calculated. Windows with this ratio < 0.1 and normalized right-pool coverage > 0.05 was considered as hemizygous in right morph. For the aligned reads of mutant individuals and R-morph plants from the same populations (when present), we removed reads with mapping quality < 10 using SAMtools version 1.3.1. We used IGV version 2.14.1 67 to investigate regions of interest.

We used the SynMap2 function (*72*) with CODEML function on CoGe (https://genomevolution.org/) to calculate the rate of synonymous substitution ($K_S$) between paralogous genes in syntenic blocks in each genome. Then we calculated the $K_S$ between *YUC-R* and its paralogues with the CODEML program (*73*, *74*) in the PAML package (*75*). We used R package GENESPACE version 1.3.1 with BRAKER3 annotations (*76*, *77*) to examine the synteny among contigs.

Gene expression analysis

We assembled the style and stamen RNA-seq reads aligned to the primary genome assembly of *W. paniculata* (see the preceding section) into a trascriptome with StringTie version 2.2.1 (*78*). We conducted functional annotation for the style+stamen trascriptome using Mercator4 v6.0 (*79*) with Prot-scriber and Swissprot annotations included. Since SMALL AUXIN UP-REGULATED (SAUR) genes were not included in the Mercator bins, we manually created a Mercator bin: 11.2.2.6, "Phytohormone action.auxin.perception and signal transduction.auxin responsive genes *(SAUR)", for genes that were annotated as SAUR genes. We performed transcript quantification with Salmon version 1.9.0 (*80*) with the "quant" command and the "--no_bowtie", "--gcBias" and "--validateMappings" settings. We conducted gene differential expression analysis with the count data using the R package DESeq2 (*81*). We compared the expression level of each gene between left- and right-morphs for each organ (style or stamen), developmental stage (early, mid, or late bud), and species (*W. paniculata* or *W. multiflora*). In each comparison, only genes with averaged normalized expression level > 100 were included in the following analyses. Then we conducted the MapMan enrichment analysis using MapMan version 3.5 (*82*), with the results of the gene differential expression analysis and the Mercator4 annotation as the input.

For small RNA-seq data, we quantified the expression levels of miR156-5p and 3p with the grep function.

#### *MIR156-R* identification and miR156-5p/3p target site prediction

Two transcripts, Wp002WmWpStrST.817 and Wp002WmWpStrST.819, which mapped to the hemizygous region, were identified in the style and stamen transcriptomes. CPC (*83*) predicted that these transcripts were unlikely to encode proteins (coding probabilities of 0.0191525 and 0.019659, respectively).

These transcripts were expressed in the styles of R-morph *W. paniculata* (Figure 3; Figure SX). Sequences corresponding to these transcripts were identified in pooled R-morph sequences of *W. brachyandra*, *W. multiflora* and *W. thyrsiflora*. Only the Wp002WmWpStrST.817 transcript was found in the *B. aurea* pooled R-morph sequences and the assembled genome. A ClustalW alignment of these sequences was submitted to the RNAz server (*84*), which predicted that the 1800 bp long Wp002WmWpStrST.817 transcript was likely to fold into nine stable RNA structures. Blastn identified that nucleotides 250 to 336 had significant similarity to the *Camellia sinensis* KT004847 csn-miR156 gene, and Arabidopsis thaliana NR_143299 ath-MIR156 precursor. This region corresponds exactly to one of the stable RNA structures predicted by RNAz (Figure SX). RNAz did not predict any stem-loop structures suggestive of primary-microRNAs for the Wp002WmWpStrST.819 transcript.

psRNAtarget (*85*) was used to predict miR156-5p/3p target sites in the stamen and style transcriptome generated in the section "Gene expression analysis".

#### PCR genotyping

Sequence reconstruction and primer design: Three of the deletions (RH2 Wm, RH1 Wm, and F215 Wp) are accompanied by insertions. To reconstruct the borders of insertions, we extracted sequencing reads that aligned to the minus strand of the 3' flanking regions of the deletions and the mates of these sequencing reads. Then we conducted multiple sequence alignment of with MUSCLE (*86*). Primers were designed using Primer3 (*87*).

Reagents, kits and machinery used for PCR genotyping: if products were to be sequenced or were particularly difficult to amplify, Q5® High-Fidelity 2X Master Mix (M0492) from New Englad Biolabs (NEB) was used. This is a hot-start polymerase, and reactions with 98°C melting temperature used this polymerase. All other genotyping PCRs used NEB One*Taq*® DNA Polymerase (M0480), NEB Deoxynucleotide (dNTP) Solution Mix (N0447), and the primers were synthesised by Inqaba Biotechnical industries (pty) Ltd, South Africa. The SimpliAmp™ Thermal

Cycler (A24811) from ThermoFisher Scientific was used for PCR cycling. All PCR products were loaded on 2% agarose gels made with SeaKem® LE Agarose (50004), 1x TAE buffer and Ethidium bromide (0.025ul/ml gel). Electrophoresis ran for 45 minutes to 80 minutes. Gels were loaded with 5 ul of either NEB Quick-Load® Purple 100 bp DNA Ladder (N0551) or NEB Quick-Load® Purple 50 bp DNA Ladder (N0556). Cycling conditions and the list of primers used can be seen in Table S5.

Populations with homostyle plants were genotyped as follows: PCRs on WT and homostyle individuals were performed using primers for miR156, YUC-R and YUC-P1 (as an internal non-E-locus gene control) to determine if there were any complete deletions of these genes. Following sequencing, all WT and homostyle individuals were PCR-genotyped again using primers to the predicted deletions/insertions. For Redelinghuys where there are two predicted YUC-R deletions/insertions, the WT and homostyle plants were screened with each set of 3 primers separately.

PCR products from one individual each were excised from agarose gels or purified directly using the QIAquick Gel Extraction Kit (28704) from Qiagen. These purified products were then Sanger sequenced at the DNA Sequencing Unit in the Central Analytical Facilities at Stellenbosch University, South Africa.

*Arabidopsis thaliana* transformation

The *W. paniculata* and *B. aurea MIR156-R* genes was amplified from gDNA using Phusion high-fidelity DNA polymerase and the primers GGGGACAAGTTTGTACAAAAAAGCAGGCTAACTCACATCTCCGCAACCCAA and GGGGACCACTTTGTACAAGAAAGCTGGGTATGGGCAAMTACCAACTAATCATTGACC (for amplification of the *W. paniculata MIR156-R*) and GGGGACAAGTTTGTACAAAAAAGCAGGCTATTATTTATGGAACTAGAAGCTGCC and GGGGACCACTTTGTACAAGAAAGCTGGGTTTATTCCATCACATACATCAACCATACACC (for *B. aurea MIR156-R*). The PCR product was recombined first into pDONR221 and then into pB2GW7. The pB2GW7-WpMIR156-R, pB2GW7-BaMIR156-R and pB2GW7 empty vectors were introduced into *Agrobacterium tumefaciens* GV3101, and Arabidopsis Col-0 plants were transformed by floral-dipping (*88*). The resulting seed was sown on a 1:1 mix of peat (Jiffy Products, Norway) and vermiculite and sprayed with 0.03% (v/v) BASTA (phosphinothricin), 0.05% (v/v) silwet L-77 at 7 and 10 days after germination to select for transgenic plants. Putative transgenic T1 progeny were confirmed by PCR genotyping and then self-fertilised to obtain T2 seed which was treated as described above. T2 plants were harvested when the first open flower was observed and the following phenotypes recorded: number of primary rosette leaves and the percentage of these leaves displaying abaxial trichomes. These data were gathered from 7 to 47 plants per independent transgenic line.

IAA measurements

IAA was extracted and quantified as described (*89*). Briefly, homogenized frozen plant materials (50 mg) were extracted by incubation with 1 ml of pre-cooled (−20°C) MTBE:MeOH (3:1, v:v) mixture. The extracts were incubated for 30 min on an orbital shaker at 4°C before sonicating them for 15 min on an ice-cooled sonication bath. The samples were centrifuged for 10 min at 10 000 *g* at 4°C. The supernatant was transferred to new microcentrifuge 2-ml tubes, a volume of 0.5 ml of acidified water (0.01% HCl) was added and the samples were again thoroughly vortexed for 1 min. After that, the samples were kept on an orbital shaker for an additional 30 min at 4°C. The samples were centrifuged at a speed of 10 000 ***g*** for 10 min at 4°C. The upper supernatant was collected and dried down using a SpeedVac concentrator. The dried pellets were resuspended in 100 µl water:methanol (50:50) solution and the resuspended samples were immediately subjected to UPLC-ESI-MS/MS hormonal analysis.

MS/MS analysis was achieved using QTRAP 6500 (AB Sciex Germany GmbH) with a multiple-reaction monitoring (MRM) scan type equipped with an electrospray ionization (ESI) source and attached to the UPLC system (Waters Acquity UPLC system; Waters). Analytical UPLC separation was achieved on a reversed-phase (RP) C18-column. (ACQUITY UPLC HSS T3 VanGuard Pre-column, 100 Å, 1.8 µm; Waters). A UPLC separation method performed using a binary solvent system consists of water containing 0.1% (v/v) formic acid (solvent A) and methanol containing 0.1% (v/v) formic acid (solvent B). The gradient parameters for RP-UPLC separation were as follows: 62% eluent A for 6.5 min; 45% eluent B from 6.5 to 7.0 min; 10% eluent A from 7.0 to 7.1 min, held at 0% eluent A from 7.1 to 8.1 min and returned to initial conditions by 8.1 min. From 8.1 to 10.0 min the column was re-equilibrated and conditioned to 62% eluent A. The autosampler temperature was set at 10°C. The injection volume was 5 µl.

Phylogenetic analysis of YUC genes

The criterion used for searching homologs of YUC-R is "at least one BLASTN match with percentage of identity higher than 90% and Bit Score higher than 40".

We used the software package BEAST v2.7.6 (*90*) to estimate the time of divergence between YUC-R genes and YUC-P genes. First, we did model selection of jModelTest (*91*) using Akaike information criterion (AIC) as the criterion. The GTR + I + Gamma model with nCat = 4 was selected. Then we set the time to the most recent ancestors of Commelinales and Zingiberales (79.8 ± 9.87 MYA), Commelinales and Poales (106.7 ± 8.3 MYA), and Monocots and Eudicots (135.76 ± 0.51 MYA) (*92*), as calibration points. Ten independent Markov Chain Monte Carlo runs, each with 1 x $10^8$ generations (first 10% was burn-in) were conducted. We combined the output of the eight runs using LogCombiner v2.7.6 (*90*) with a sample frequency of 5,000. Then we used TreeAnnotator v2.7.6 (*90*) to generate the maximum clade credibility tree, which was visualized in FigTree v1.4.4 (http://tree.bio.ed.ac.uk/software/figtree/) (Figure S15A).

We used the HyPhy platform (https://www.hyphy.org/) together with MUSCLE (*86*) and IQ-TREE v1.6.12 (*93*) to estimate the dN/dS values of *YUC-R* and *YUC-P* lineages. When using IQ-TREE, bootstrapping was conducted 1,000 times.

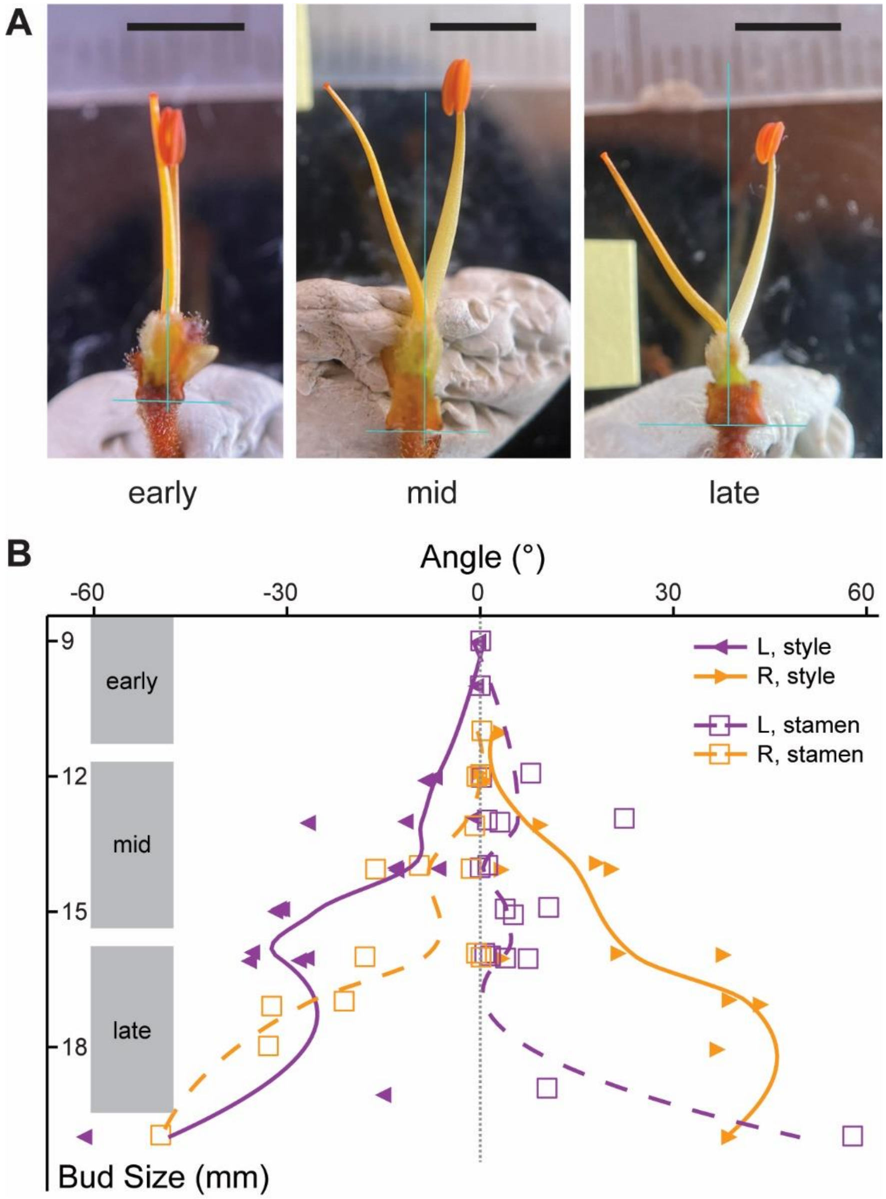


**Fig. S1. Development of style and stamen deflection**

(**A**) Images of dissected *Wachendorfia paniculata* flower buds from the indicated stages, retaining the gynoecium and the opposing stamen. The blue reference line along the flower midline was used to measure the angle of style and stamen deflection. Scale bar is 5 mm.

(**B**) Quantification of style (triangles) and stamen (squares) deflection across early, mid and late-stage buds from top to bottom. Lines are Loess trendlines.

Type or paste caption here. Create a page break and paste in the figure above the caption.

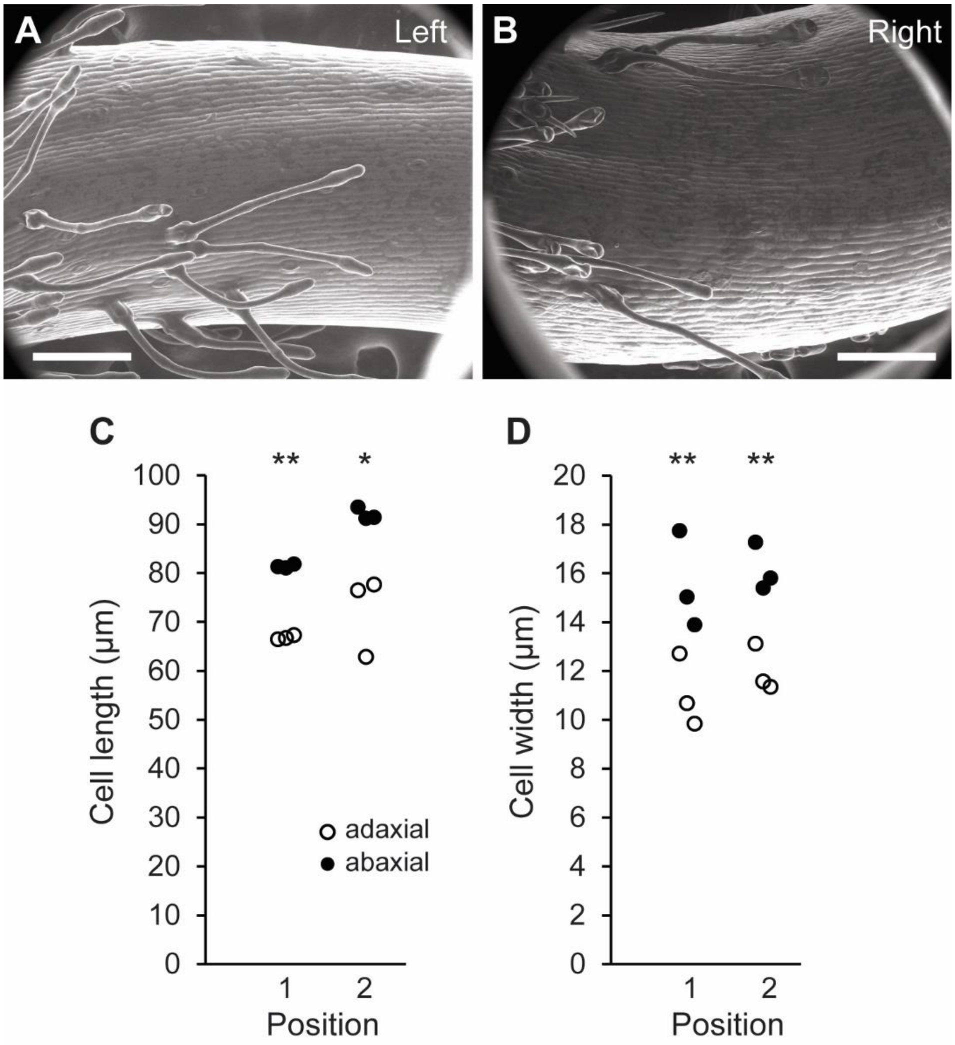


**Fig. S2. Twisting cell files and differences in cell size between adaxial and abaxial sides of *Wachendorfia paniculata* styles**

(**A**, **B**) SEM images of style bases of L- (A) and R-morph (B) flowers showing twisted cell files. Scale bars are 200 μm.

(**C**, **D**) Measurements of cell length (C) and cell width (D) from two paired positions on the adaxial and abaxial sides of ESEM images of *Wachendorfia paniculata* R-morph styles from open flowers. Each dot is the average from five cells, $n$ = 3 styles. Asterisks indicate significant differences between adaxial and abaxial sides based on paired $t$-test a $p<0.05$ (*) and $p<0.01$ (**).

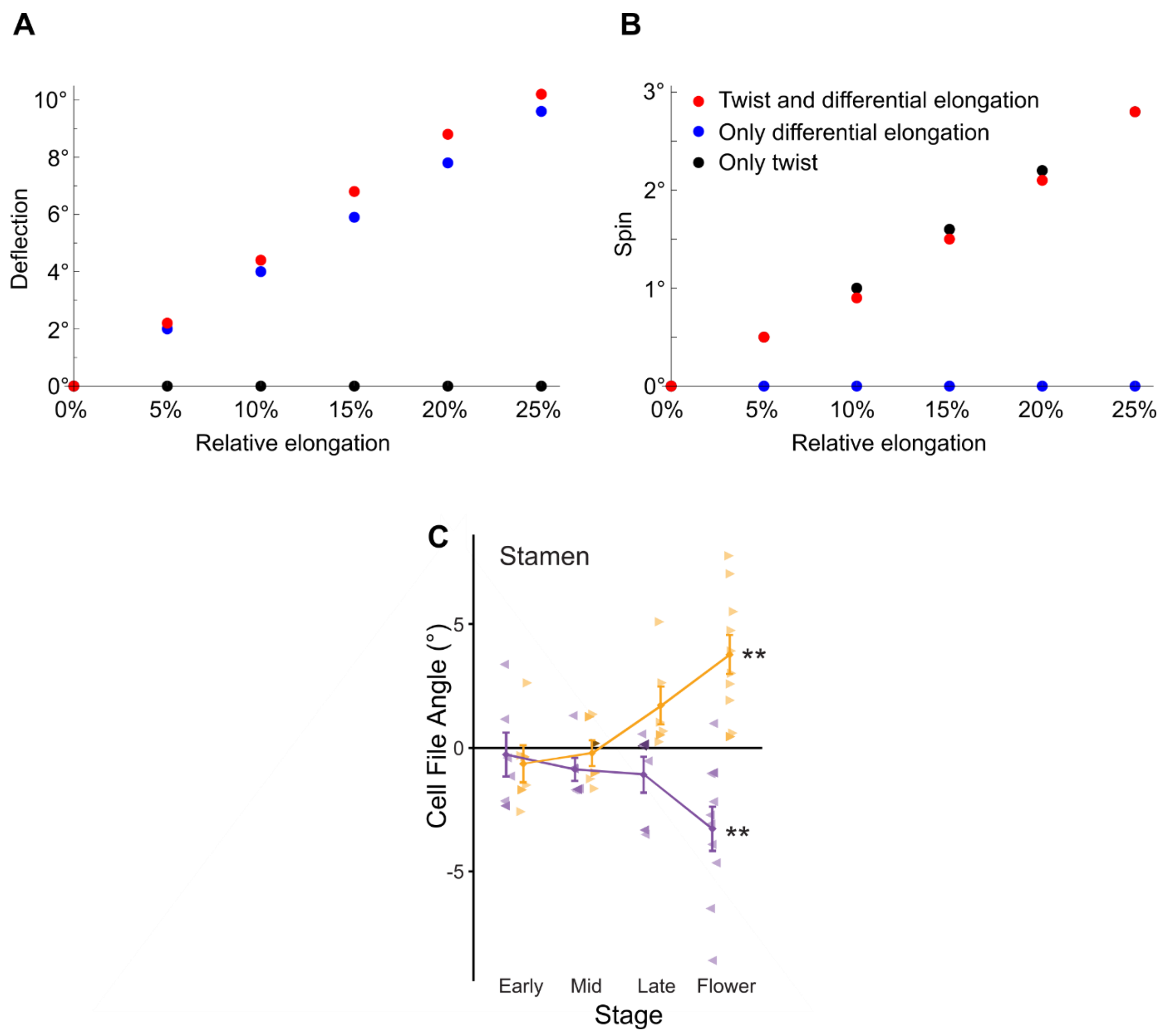


**Fig. S3. Model output under different scenarios and cell-file twisting in stamens**

(**A**, **B**) Relationship between the relative longitudinal elongation and the resulting (A) deflection angle and (B) spin of the top layer of the structure. All angles and elongation values are referenced to the initial spring length and orientation of the top layer. Data points represent: full model with both differential elongation and twist (red), model with differential elongation only (blue), and model with twist only (black).

(**C**) Quantification of cell-file angles at the bases of stamen filaments from different stage buds as defined in Figure S1. Asterisks indicate significant differences from 0 at $p<0.05$ (*), $p<0.01$ (**), $p<0.001$ (***) as determined by one-sample $t$-test.

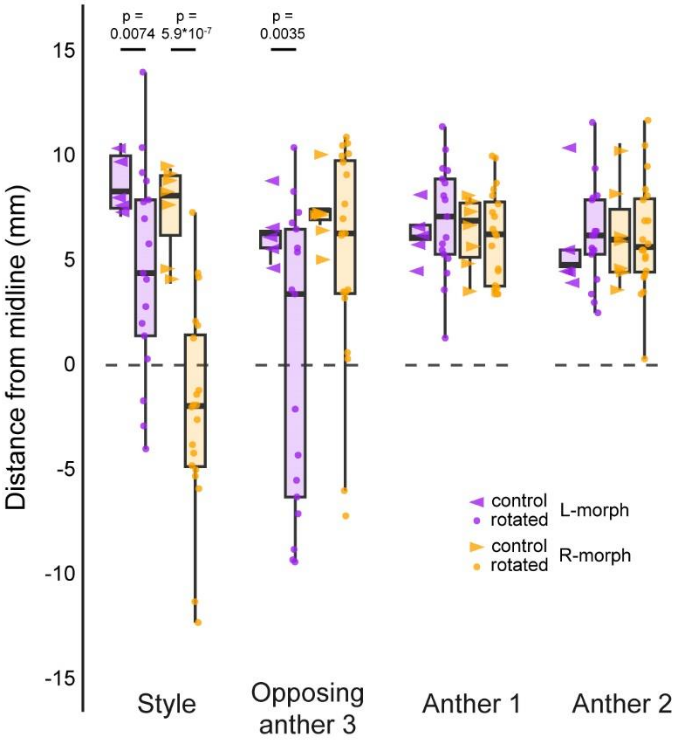


**Fig. S4. Effect of bud rotation on organ deflection**

Distances from the midline are shown for the indicated organs from flowers without (control, arrowheads) and with rotation for 15 hours before opening (rotated, circles) from L- and R-morph plants. Anther 1 is the adaxial anther on the same side as the style, Anther 2 is the adaxial anther on the other side from the style, i.e. on the same side as anther 3. *p*-values are from two-sample *t*-tests comparing control and rotated flowers. Comparisons without indicated *p*-values were not significant ($p>0.05$).

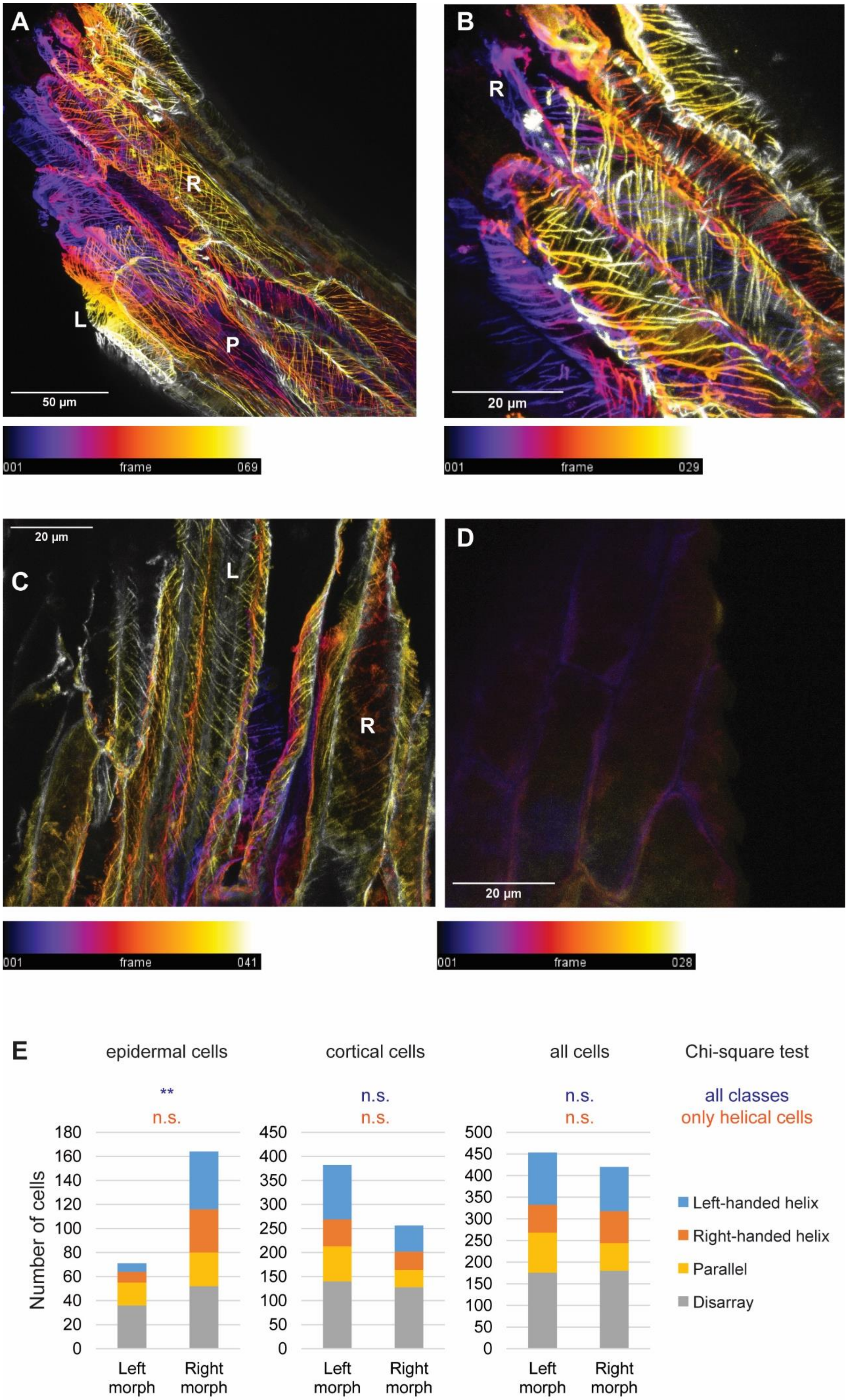

A
R
L
P
50 µm
001
frame
069
B
R
20 µm
001
frame
029
C
20 µm
L
R
001
frame
041
D
20 µm
001
frame
028
E
epidermal cells
cortical cells
all cells
Chi-square test
**
n.s.
n.s.
n.s.
n.s.
n.s.
all classes
only helical cells
Number of cells
Left morph
Right morph
Left-handed helix
Right-handed helix
Parallel
Disarray

**Fig. S5. Immunolocalization of cortical microtubules in *Wachendorfia thyrsiflora* styles**

(**A**-**D**) Confocal microscopy z-stacks of longitudinal style sections from flowers of *Wachendorfia thyrsiflora* stained with mouse anti-tubulin antibody E7 (A,B) or DM1A (C) or no primary antibody (D) and secondary anti-mouse antibody labelled with Alexa FluorTM488. The false colors indicate depth along the z-axis as indicated by the color scales below, with 0 most distant from the viewer. L, R, P indicate cells with left-handed microtubule helix, right-handed microtubule helix or parallel microtubules, respectively. Scale bars are defined in the figure.

(**E**) Quantification of microtubule orientation in longitudinal style sections from flowers of left- and right-morph *W. thyrsiflora* plants. Left to right panels show epidermal cells, cortical cells and all cells. Results from Chi-square test for independence are shown across all four classes (top row) and only comparing cells with left- and right-handed helices (bottom row). **, significantly different from expected at $p<0.01$ after Bonferroni correction.

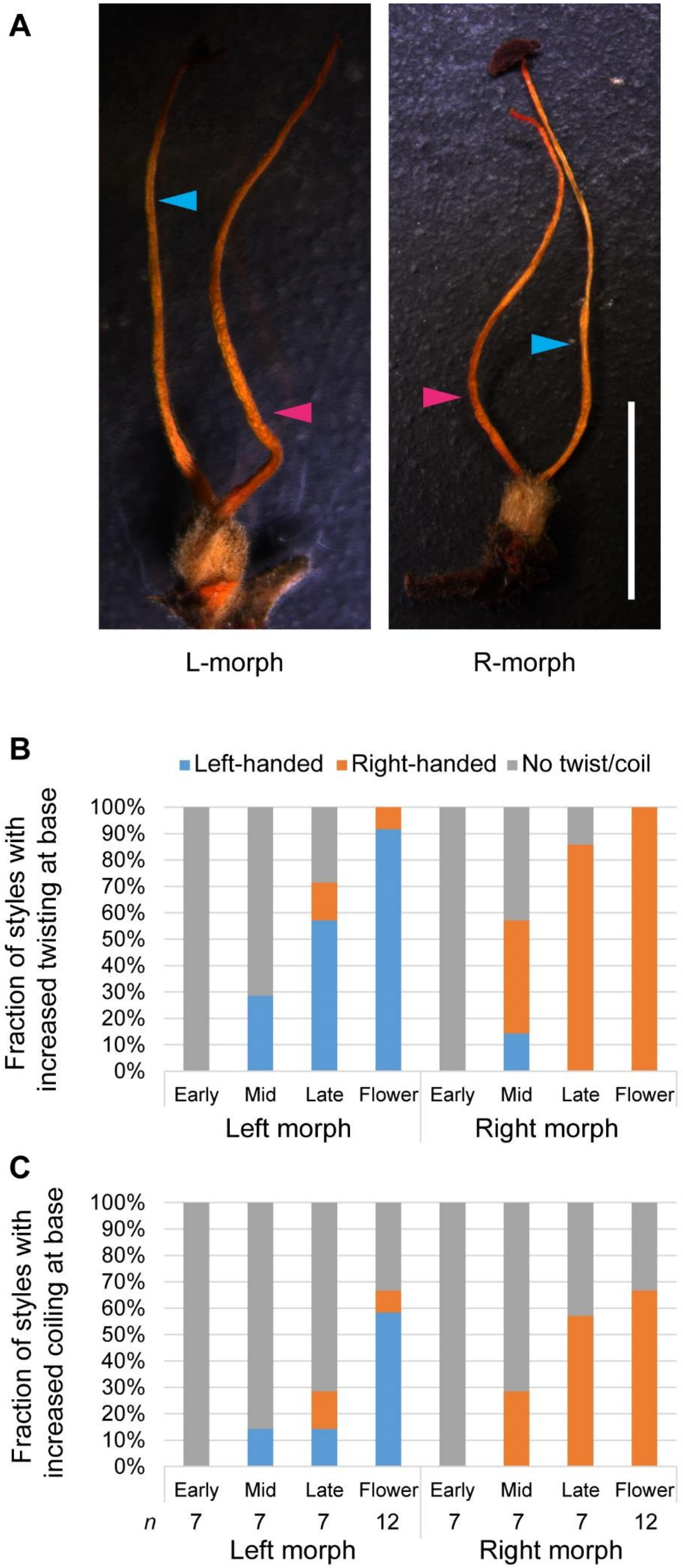
A
L-morph
R-morph
B
Left-handed
Right-handed
No twist/coil
Fraction of styles with increased twisting at base
100%
90%
80%
70%
60%
50%
40%
30%
20%
10%
0%
Early
Mid
Late
Flower
Early
Mid
Late
Flower
Left morph
Right morph
C
Fraction of styles with increased coiling at base
100%
90%
80%
70%
60%
50%
40%
30%
20%
10%
0%
Early
Mid
Late
Flower
Early
Mid
Late
Flower
n
7
7
7
12
7
7
7
12
Left morph
Right morph

**Fig. S6. *Wachendorfia* styles have an intrinsic chirality**

(**A**) Dried styles (pink arrowhead) and opposing stamens (blue arrowhead) from mature L-morph (left) and R-morph (right) flowers. Styles show coiling. Remaining floral organs have been removed prior to drying. Scale bar is 5 mm.

(**B**, **C**) Fraction of styles showing increased twisting (B) or coiling (C) with the indicated handedness or no increased twisting or coiling (grey) from buds of the indicated stages and open flowers. *n* is indicated below the bars in (C) and refers to both graphs.

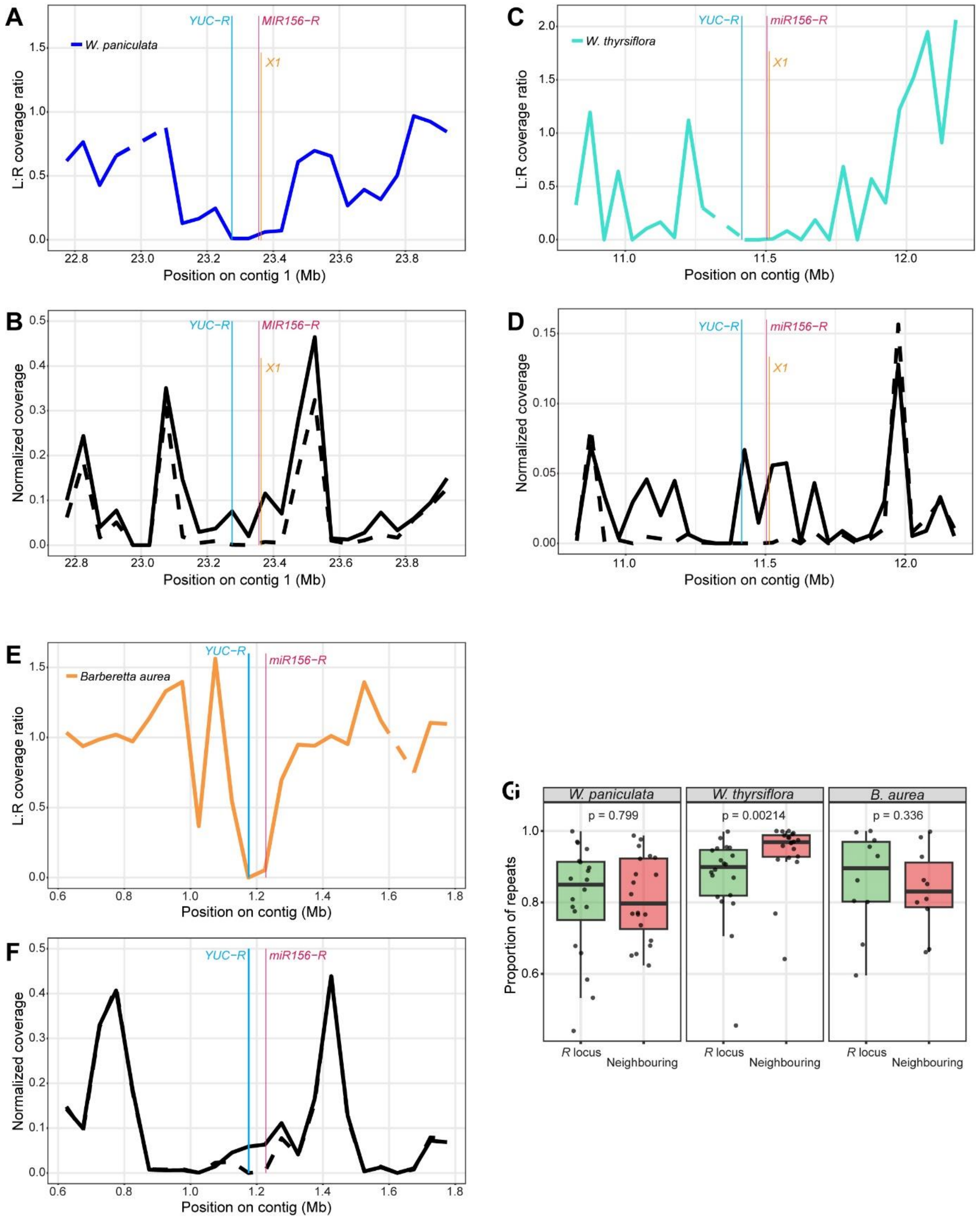
A
W. paniculata
YUC−R
MIR156−R
X1
L:R coverage ratio
Position on contig 1 (Mb)
B
Normalized coverage
C
W. thyrsiflora
miR156−R
Position on contig (Mb)
D
E
Barberetta aurea
F
G
W. paniculata
W. thyrsiflora
B. aurea
p = 0.799
p = 0.00214
p = 0.336
Proportion of repeats
R locus
Neighbouring

**Fig. S7. Identification of the *R*-locus supergenes in *Wachendorfia paniculata, Wachendorfia thyrsiflora* and *Barberetta aurea***

(**A**-**F**) Coverage ratios of whole-genome sequencing reads from L- versus R-morph pools (A,C,E) and normalized coverage values from R-pools (continuous lines) and L-pools (dashed lines) (B,D,F) from *W. paniculata* (A,B), *W. thyrsiflora* (C,D) and *B aurea* (E,F) mapped against their respective reference assemblies. Location of *R*-locus genes is indicated by coloured vertical lines. Note the absence of the *X1* gene from the *R* locus in *B. aurea*.

(**G**) Proportion of repeats in 20 (for *W. paniculata* and *W. thyrsiflora*) or 10 (for *B. aurea*) 10-kb windows in the *R* loci and in 20 (for *W. paniculata* and *W. thyrsiflora*) or 10 (for *B. aurea*) 10-kb windows in neighbouring regions of the indicated species. *p*-values are from Wilcoxon rank sum test.

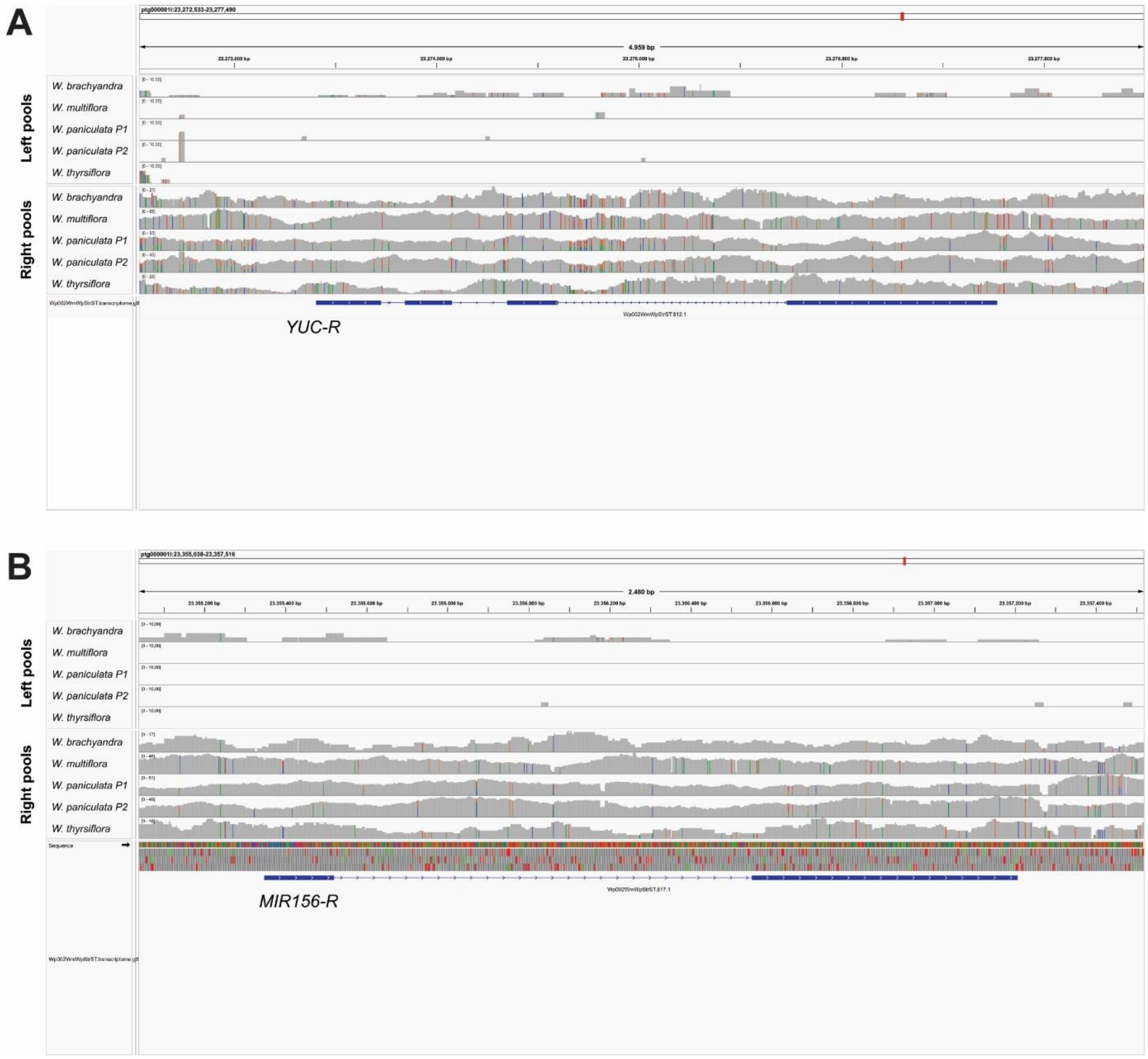


**Fig. S8. Identification of hemizygous *R*-locus genes in four *Wachendorfia* species and in *Barberetta aurea***

(**A**, **B**) Integrated Genome Viewer (IGV) screenshots showing the mapping of the L- and R-pools from the indicated species against the *R*-locus genes *YUC-R* (A) and *MIR156-R* (B). Reads from the pools were mapped against the *W. paniculata* reference genome. Gene models are shown in blue below.

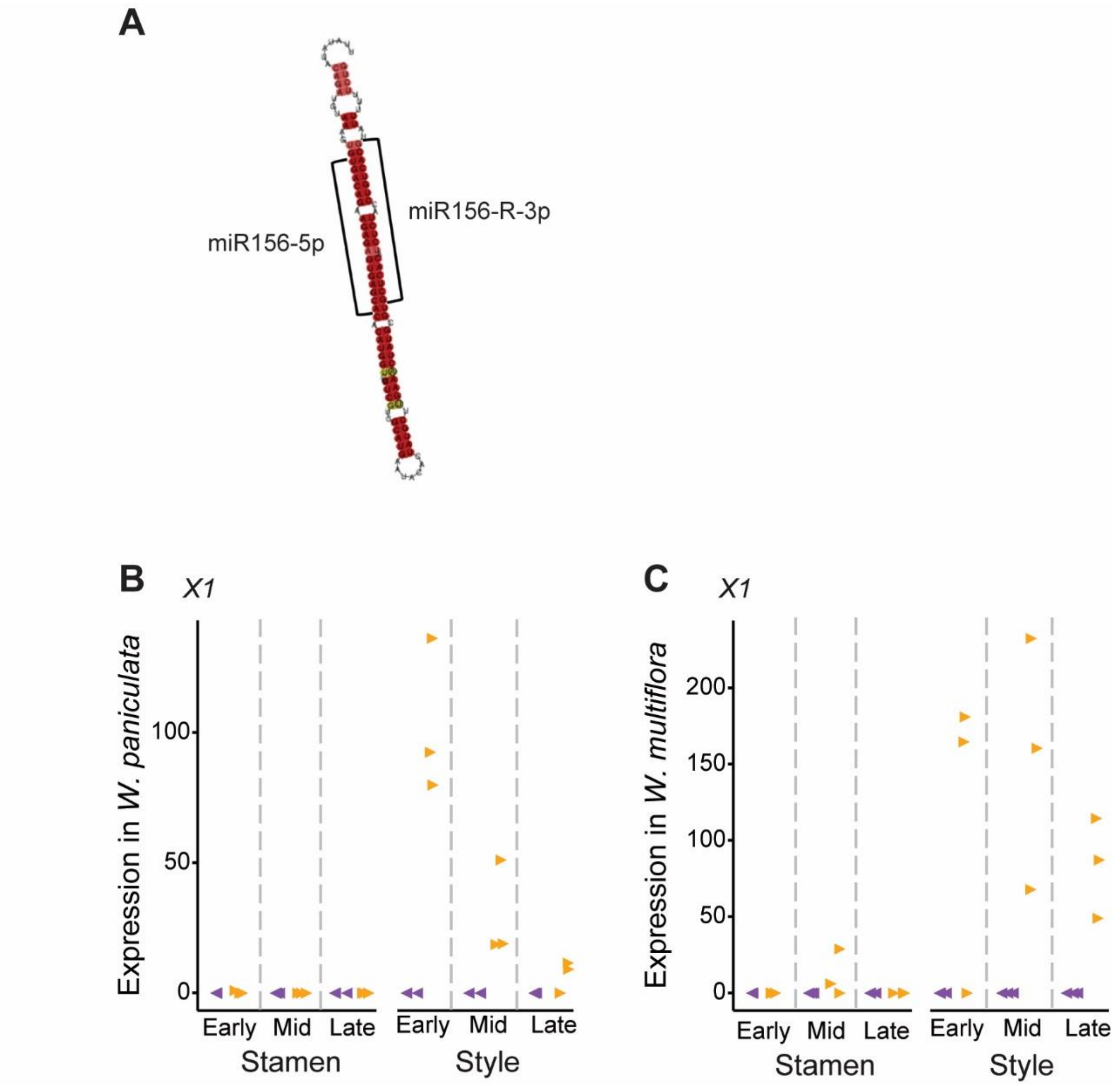


**Fig. S9. Non-coding RNA-genes at the *R* locus**

(**A**) RNAz output for pri-miR156-R.

(**B**, **C**) Expression of the *X1* transcript in styles and opposing-stamen filaments of different-stage buds from *Wachendorfia paniculata* (B) and *W. multiflora* (C) based on RNA-seq. Expression is in normalized read counts.

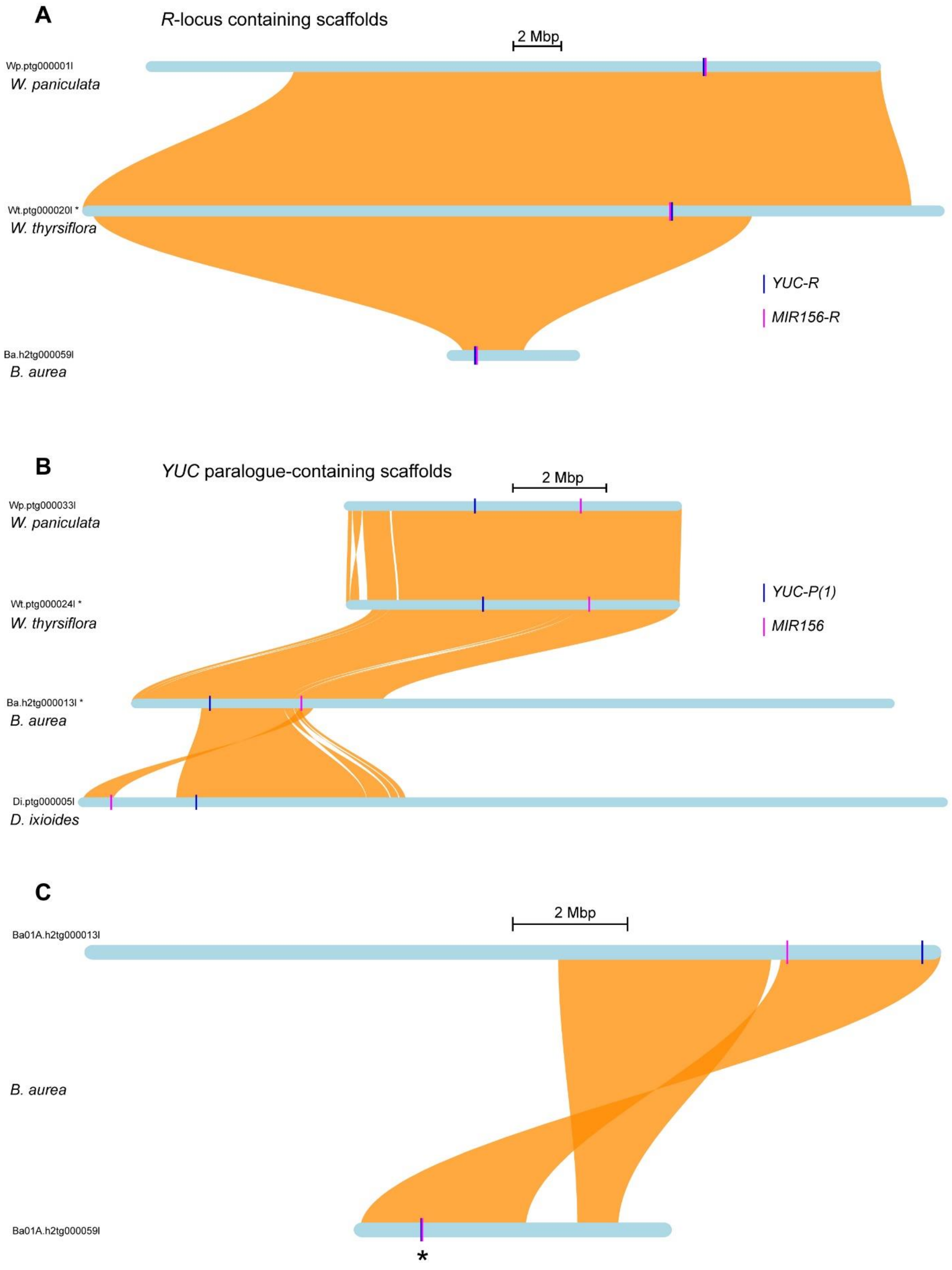


(continued on next page)

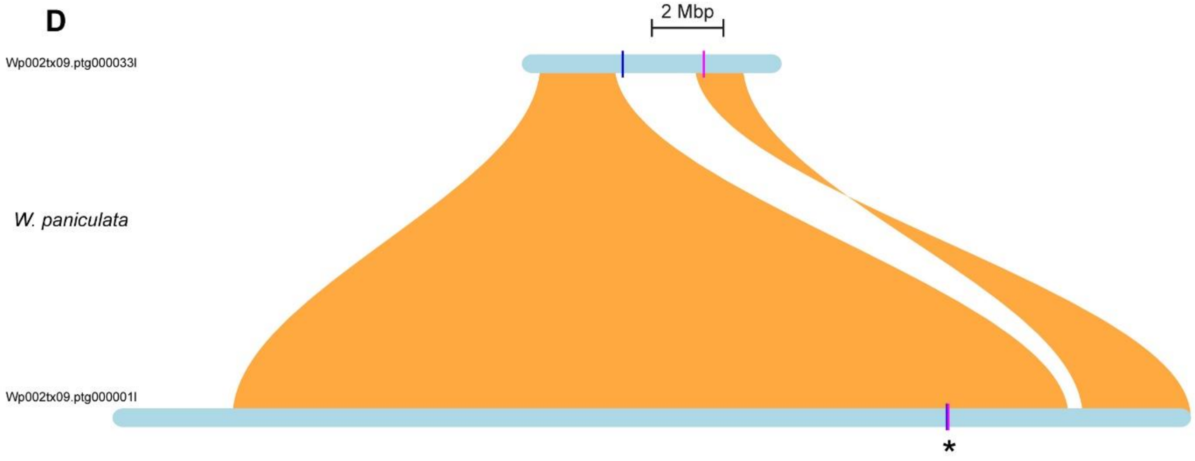


**Fig. S10. Synteny analyses**

(**A**, **B**) Synteny plots generated by GENESPACE comparing (A) the *R*-locus containing scaffolds of *W. paniculata*, *W. thyrsiflora* and *B. aurea*, (B) the *YUC-P(1)* paralogue containing scaffolds of *W. paniculata*, *W. thyrsiflora*, *B. aurea* and *D. ixioides*.

(**C**, **D**) Synteny plots generated by GENESPACE comparing the R-locus containing and the paralogue-containing scaffolds in *B. aurea* (C) and *W. paniculata* (D). The *R*-locus is marked by an asterisk in (C, D).

The GENESPACE output underlying these plots is given in Data S2.

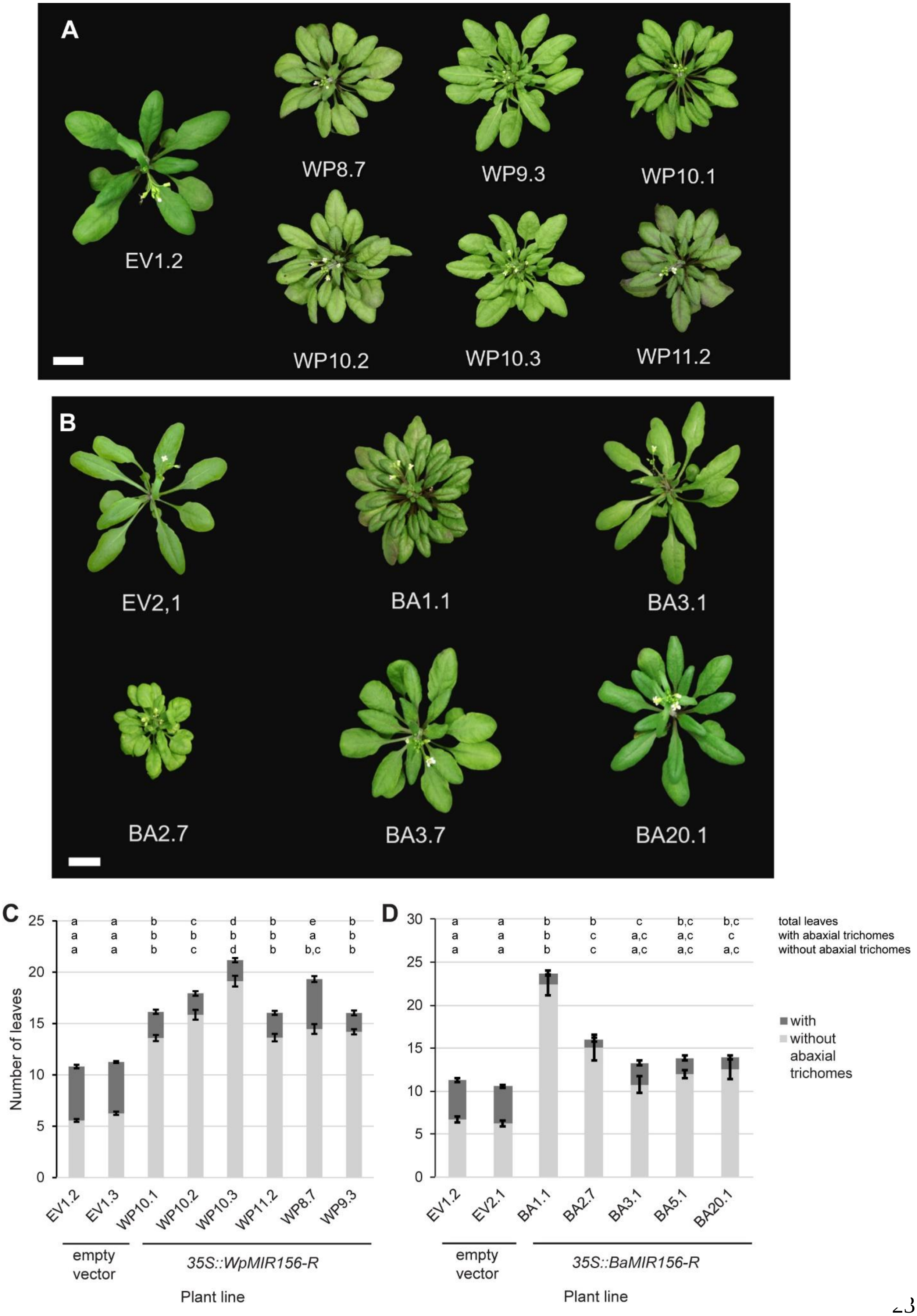
A
WP8.7
WP9.3
WP10.1
EV1.2
WP10.2
WP10.3
WP11.2
B
EV2,1
BA1.1
BA3.1
BA2.7
BA3.7
BA20.1
C
Number of leaves
EV1.2
EV1.3
WP10.1
WP10.2
WP10.3
WP11.2
WP8.7
WP9.3
empty vector
35S::WpMIR156-R
Plant line
D
EV1.2
EV2.1
BA1.1
BA2.7
BA3.1
BA5.1
BA20.1
empty vector
35S::BaMIR156-R
Plant line
total leaves
with abaxial trichomes
without abaxial trichomes
with
without
abaxial
trichomes

**Fig. S11. Overexpression of *Wachendorfia paniculata* and *Barberetta aurea MIR156-R* genes in *Arabidopsis thaliana***

(**A**) Transgenic *A. thaliana* plants carrying an empty vector (EV) or a *35S::WpMIR156-R* (WP) construct.

(**B**) Transgenic *A. thaliana* plants carrying an empty vector (EV) or a *35S::BaMIR156-R* (BA) construct.

(**C**, **D**) Number of juvenile (without abaxial trichomes) and adult leaves (with abaxial tricomes) formed by transgenic *A. thaliana* plants carrying an empty vector (EV) or a *35S::WpMIR156-R* (WP) construct (C) or an empty vector (EV) or a *35S::BaMIR156-R* (BA) construct (D) at flowering. Each bar represents transgene-carrying T2 plants derived from an independent transformation event. *n* is between 7 and 47 T2 plants per line. Values are means ± SEM. Letters above indicate significant differences for total leaf number (upper row), leaves with (middle row) and without abaxial trichomes (lower row) based on ANOVA followed by Tukey's HSD test at $p<0.05$.

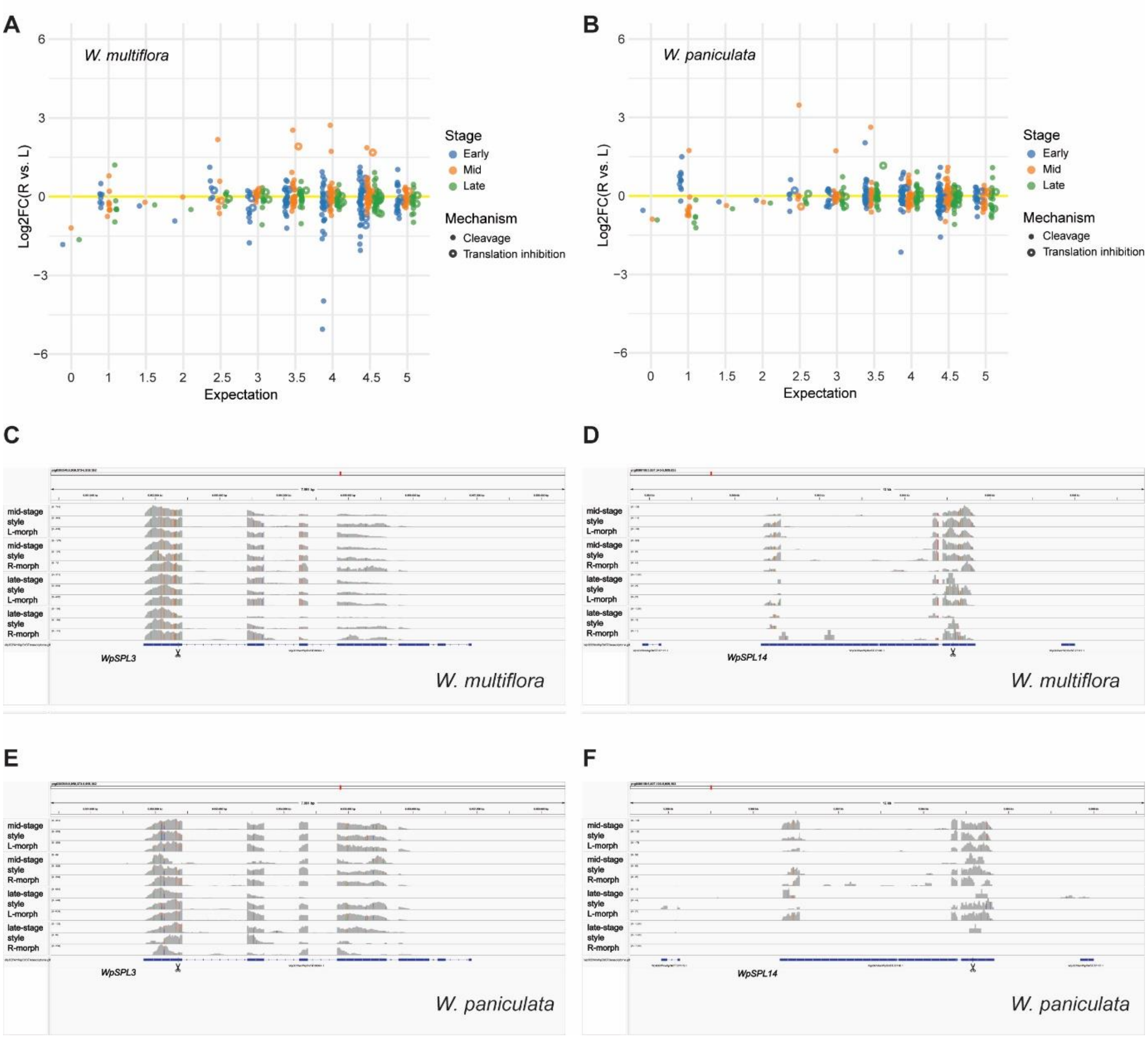


**Fig. S12. No evidence of transcript cleavage by miR156-5p in developing styles**

(**A**, **B**) $Log_2$-fold changes ($Log_2FC$) of transcript abundance between R- and L-morph styles at the indicated developmental stages from *W. multiflora* (A) and *W. paniculata* (B). Transcripts shown are ones predicted to be targets of miR156-5p/3p in *Wachendorfia*. Expectation value on x-axis is based on the output of psRNATarget, with lower values indicating higher confidence for targeting. Predicted mode of action (transcript cleavage or translation inhibition) on the targets is shown by filled or open circles, respectively. Abundance changes were not significant for any of the transcripts. The predicted targets are given in Data S3.

(**C**-**F**) IGV screenshots from mapping of RNA-seq reads from mid-stage styles of L- or R-morph plants against the *W. paniculata* genome in the region of the predicted *miR156* targets *SPL3* (C, E) and *SPL14* (D, F). Samples were from *W. multiflora* (C, D) or *W. paniculata* (E, F) as indicated. Predicted *miR156* cleavage sites are indicated by scissors.

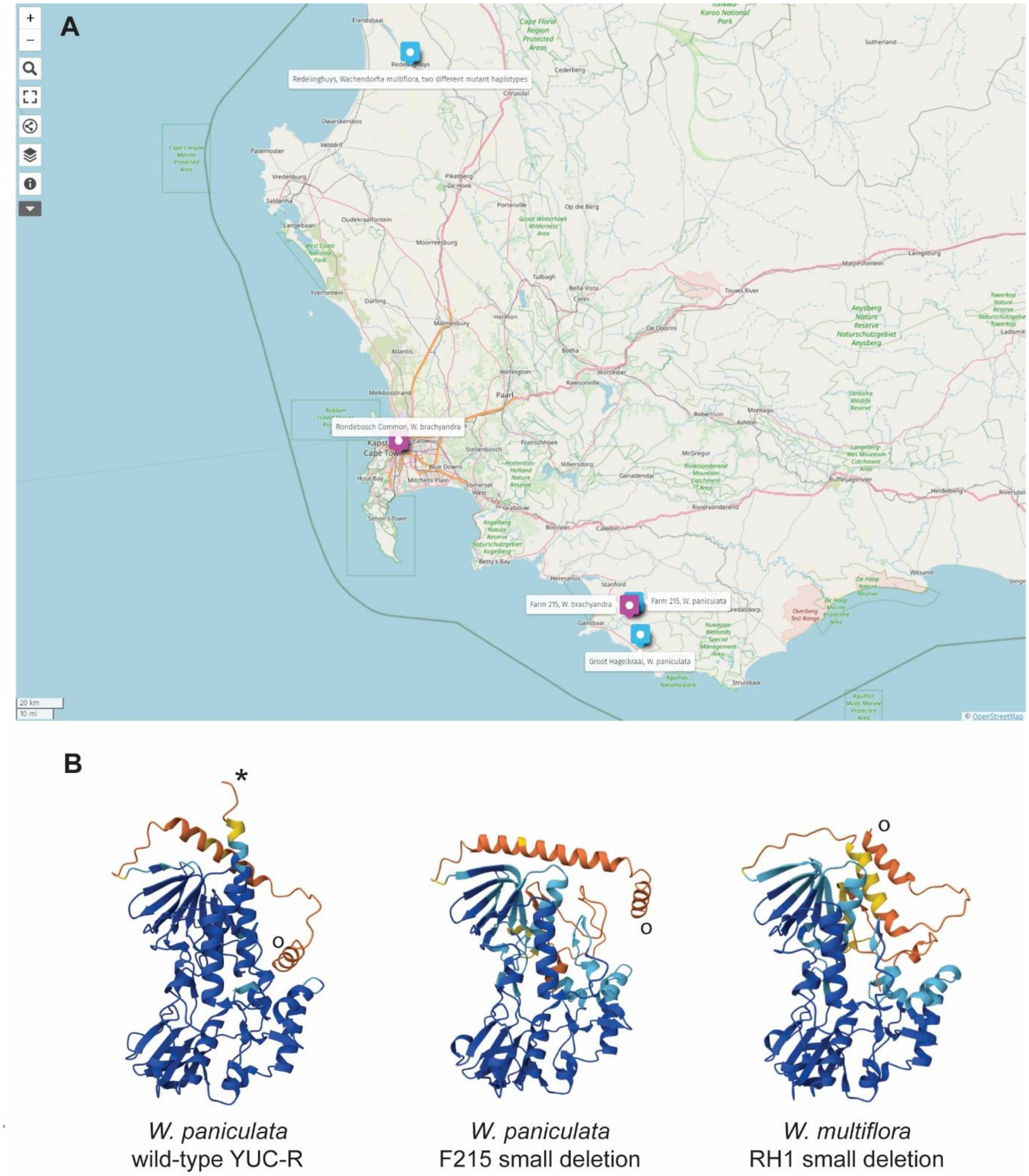


**Fig. S13. Homostylous mutants from natural *Wachendorfia* populations**

(**A**) Geographical origins of different homostylous *Wachendorfia* mutants from the Western Cape province of South Africa. Blue symbols indicate right-homostyles, purple symbols show left homostyles.

(**B**) Predicted structures of proteins encoded by the indicated *YUC-R* alleles based on AlphaFold3. The N-termini are indicated by the open circles, the C-terminus is shown by the asterisk in the wild-type structure. Structures were aligned based on the confidently predicted regions in blue.

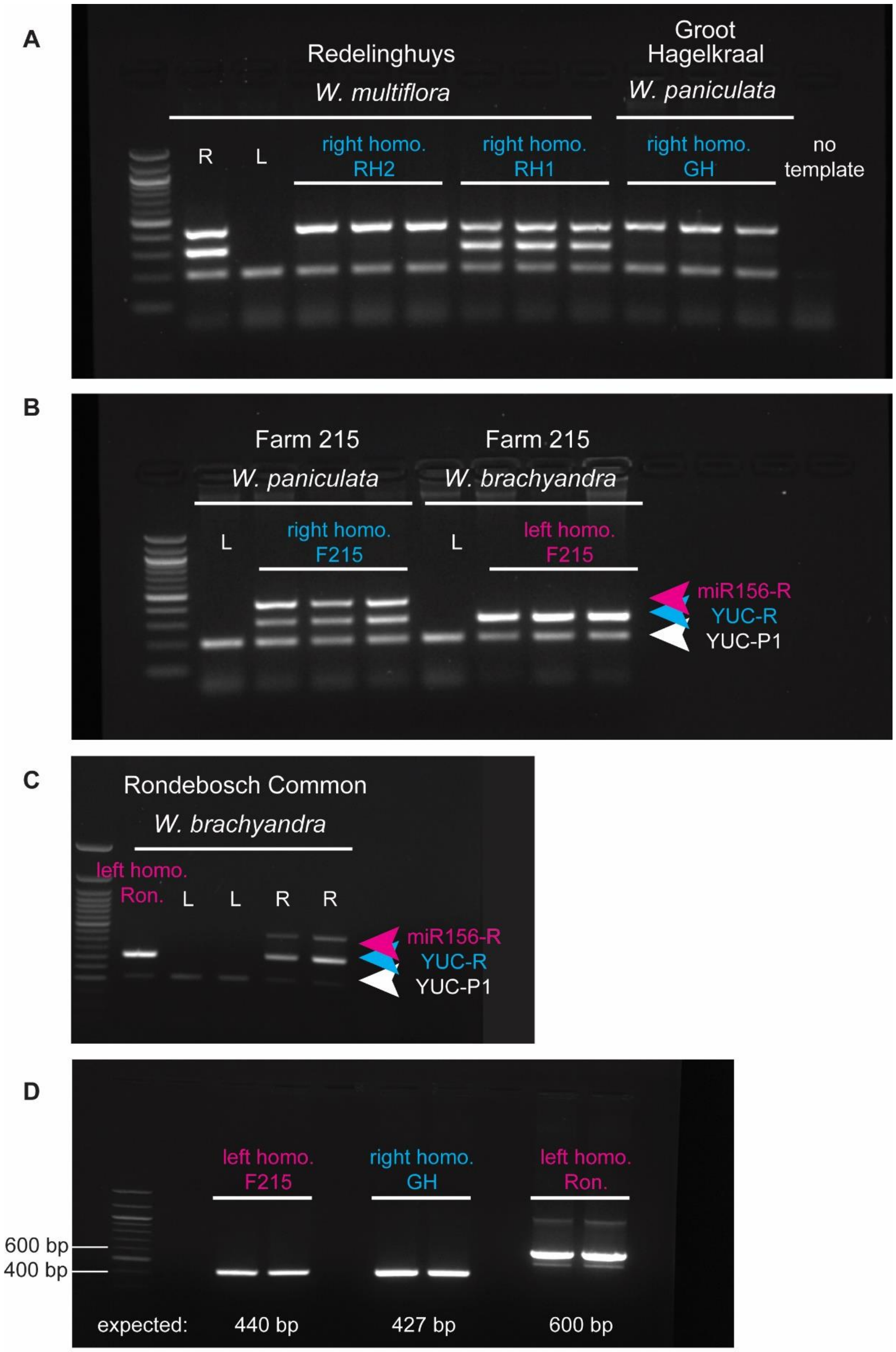


**Fig. S14. PCR genotyping of homostylous *Wachendorfia* mutants**

(**A**-**C**) PCR genotyping of plants with the indicated phenotype from the populations shown above with three primer pairs against *MIR156-R*, *YUC-R* and *YUC-P1*, as indicated by colored arrowheads. Annotation in (B) also applies to (A).

(**D**) PCR genotyping of homostylous mutants with primer pairs specific to the predicted deletion alleles. Expected sizes are indicated underneath. Sanger sequences from these PCR products are shown in DataS4.

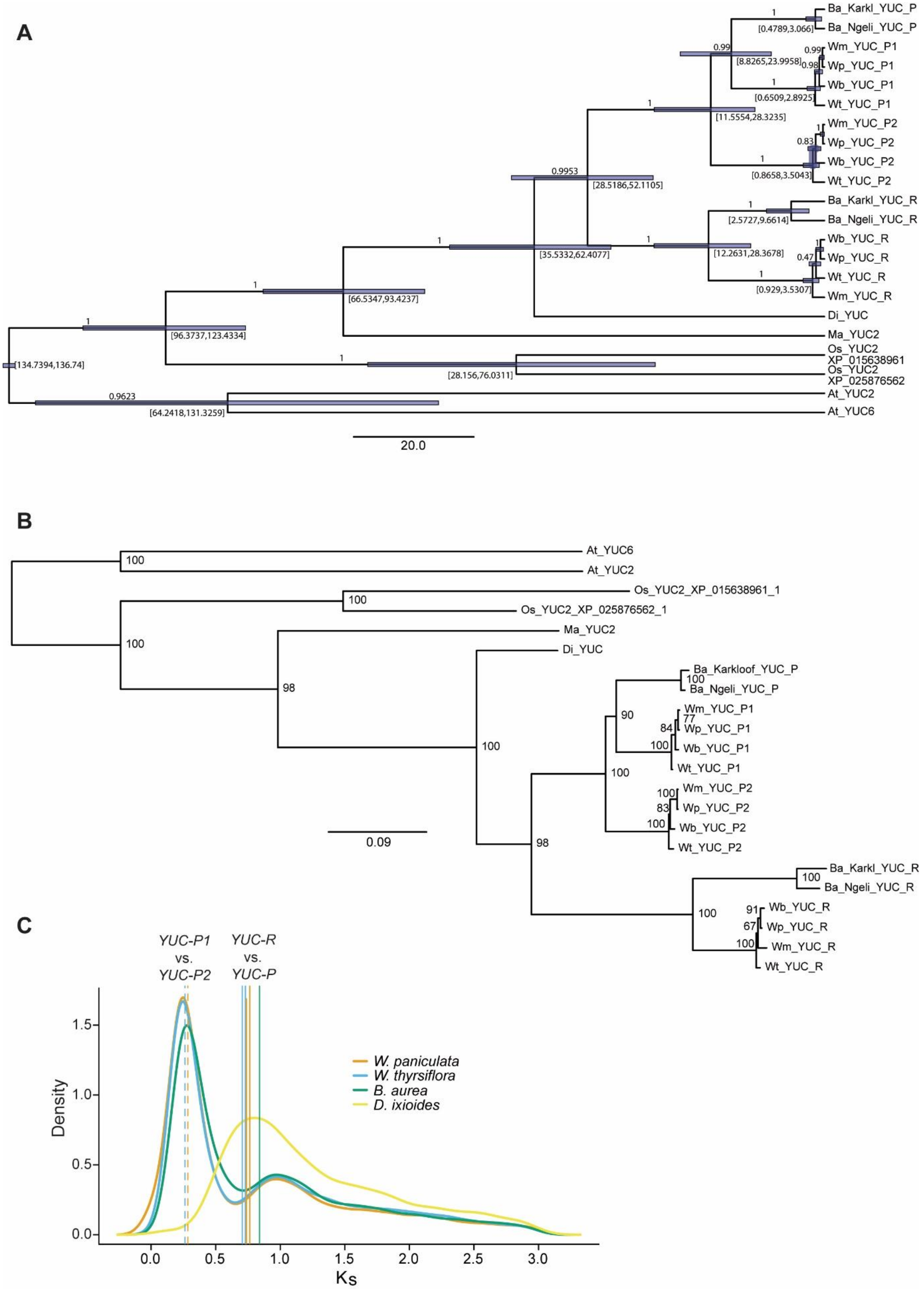
A
Ba_Karkl_YUC_P
Ba_Ngeli_YUC_P
Wm_YUC_P1
Wp_YUC_P1
Wb_YUC_P1
Wt_YUC_P1
Wm_YUC_P2
Wp_YUC_P2
Wb_YUC_P2
Wt_YUC_P2
Ba_Karkl_YUC_R
Ba_Ngeli_YUC_R
Wb_YUC_R
Wp_YUC_R
Wt_YUC_R
Wm_YUC_R
Di_YUC
Ma_YUC2
Os_YUC2
XP_015638961
Os_YUC2
XP_025876562
At_YUC2
At_YUC6
[0.4789,3.066]
[8.8265,23.9958]
[0.6509,2.8925]
[11.5554,28.3235]
[0.8658,3.5043]
[28.5186,52.1105]
[2.5727,9.6614]
[35.5332,62.4077]
[12.2631,28.3678]
[0.929,3.5307]
[66.5347,93.4237]
[96.3737,123.4334]
[134.7394,136.74]
[28.156,76.0311]
[64.2418,131.3259]
0.99
0.98
0.83
0.9953
0.47
0.9623
20.0
B
At_YUC6
At_YUC2
Os_YUC2_XP_015638961_1
Os_YUC2_XP_025876562_1
Ma_YUC2
Di_YUC
Ba_Karkloof_YUC_P
Ba_Ngeli_YUC_P
Wm_YUC_P1
Wp_YUC_P1
Wb_YUC_P1
Wt_YUC_P1
Wm_YUC_P2
Wp_YUC_P2
Wb_YUC_P2
Wt_YUC_P2
Ba_Karkl_YUC_R
Ba_Ngeli_YUC_R
Wb_YUC_R
Wp_YUC_R
Wm_YUC_R
Wt_YUC_R
0.09
C
YUC-P1 vs. YUC-P2
YUC-R vs. YUC-P
Density
Ks
W. paniculata
W. thyrsiflora
B. aurea
D. ixioides

**Fig. S15. Evolution of *YUC-R***

(**A**) Phylogenetic tree of *YUC-R* and most closely related homologues from the indicated species generated by BEAST. Branch support values based on posterior probabilities are given above the branches. Dating of divergence times with 95% confidence intervals is shown below the branches, if possible to the right of the branch point.

(**B**) Phylogenetic tree of *YUC-R* coding sequence and most closely related homologues from the indicated species generated by IQ-TREE. Branch support values based on bootstrapping 1,000 times are shown.

(**C**) Genome-wide Ks distributions between syntenologs as determined by CoGe. The peak at Ks ≈ 1.0 most likely corresponds to the τ whole-genome duplication. The peak at Ks ≈ 0.25 represents a *Wachendorfia*/*Barberetta*-specific whole-genome duplication. Solid vertical lines indicate Ks values comparing *YUC-R* to the *YUC-P* paralogue sequences from the three species indicated by line colour. Dashed vertical lines indicate Ks values comparing *YUC-P1* and *YUC-P2* in the two *Wachendorfia* species.

At: *Arabidopsis thaliana*, Ba: *Barberetta aurea*, Di: *Dilatris ixioides*, Ma: *Musa acuminata*, Os: *Oryza sativa*, Wb: *Wachendorfia brachyandra*, Wm: *Wachendorfia multiflora*, Wp: *Wachendorfia paniculata*, Wt: *Wachendorfia thyrsiflora*. The sequences used are given in Data S5.

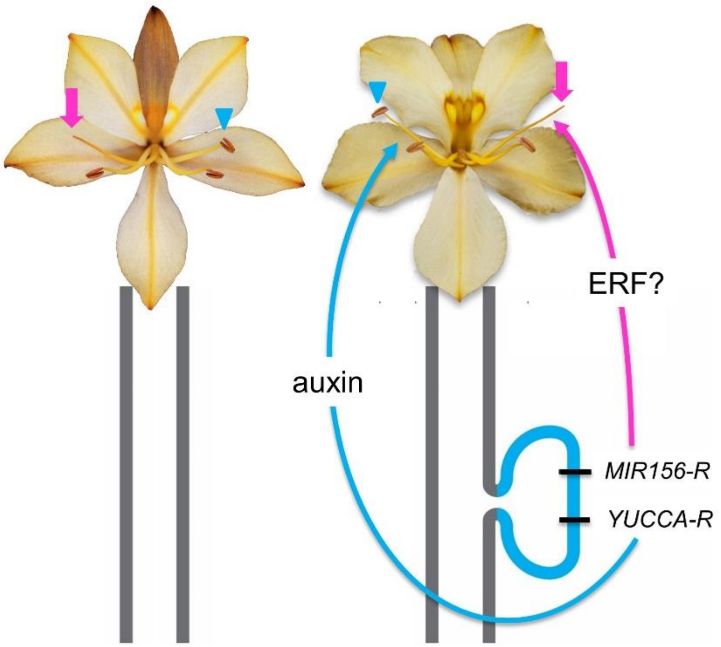


**Fig. S16. Model for the genetic control of enantiostyly in *Wachendorfia* and *Barberetta***

L-morph plants (left) are homozygous for the chromosome lacking the *R* locus. R-morph plants (right) carry the *R*-locus supergene with *YUC-R* and *MIR156-R* as the genes responsible for reversing the stamen and style orientation relative to the L-morph situation. *YUC-R* expression in the filament of the opposing stamen causes increased auxin signalling, while *MIR156-R* activity may modulate *ERF* expression.

**Table S1. Properties of genome assemblies for *Wachendorfia paniculata*, *Wachendorfia thyrsiflora* and *Barberetta aurea***

| **Species** | *Wachendorfia paniculata* | | | *Wachendorfia thyrsiflora* | | | *Barberetta aurea* | | | *Dilatris ixioides* | | |
|---|---|---|---|---|---|---|---|---|---|---|---|---|
| **Sequencing technology** | PacBio CCS | | | PacBio CCS | | | PacBio CCS | | | PacBio CCS | | |
| **Amount and depth** | 37.30 Gbp (54x) | | | 35.71 Gbp (60x) | | | 27.55 Gbp (49x) | | | 103.7 Gbp (161x) | | |
| **Assemblers** | hifiasm 0.19.0-r534 | | | hifiasm 0.19.0-r534 | | | hifiasm 0.19.0-r534 | | | hifiasm 0.19.0-r534 | | |
| | Primary assembly | Haplotype 1 | Haplotype 2 | Primary assembly | Haplotype 1 | Haplotype 2 | Primary assembly | Haplotype 1 | Haplotype 2 | Primary assembly | Haplotype 1 | Haplotype 2 |
| **Assembly length (bp)** | 688,641,917 | 656,731,322 | 572,172,692 | 598,071,203 | 400,635,364 | 403,089,839 | 563,910,667 | 538,602,047 | 500,835,959 | 645,732,908 | 637,701,696 | 514,352,464 |
| **Number of contigs** | 263 | 548 | 409 | 683 | 897 | 365 | 1,064 | 1,479 | 898 | 2,813 | 3,165 | 734 |
| **Contig N50 (bp)** | 14,422,858 | 9,034,623 | 7,829,804 | 34,200,289 | 18,055,458 | 18,306,225 | 15,046,265 | 8,233,157 | 7,065,229 | 12,068,788 | 10,162,248 | 9,549,551 |
| **BUSCO scores* (dataset: embryophyta_odb10)** | C:98.8%[S:79.3%,D:19.5%],F:0.6%,M:0.6%,n:1614 | C:98.5%[S:79.0%,D:19.5%],F:0.6%,M:0.9%,n:1614 | C:97.6%[S:79.9%,D:17.7%],F:0.7%,M:1.7%,n:1614 | C:98.8%[S:78.9%,D:19.9%],F:0.6%,M:0.6%,n:1614 | C:76.7%[S:65.8%,D:10.9%],F:1.0%,M:22.3%,n:1614 | C:75.5%[S:66.0%,D:9.5%],F:1.3%,M:23.2%,n:1614 | C:98.3%[S:84.4%,D:13.9%],F:0.8%,M:0.9%,n:1614 | C:97.6%[S:84.4%,D:13.2%],F:0.9%,M:1.5%,n:1614 | C:97.1%[S:83.5%,D:13.6%],F:0.9%,M:2.0%,n:1614 | C:98.9%[S:96.8%,D:2.1%],F:0.6%,M:0.5%,n:1614 | C:97.8%[S:95.0%,D:2.8%],F:0.6%,M:1.6%,n:1614 | C:98.8%[S:95.9%,D:2.9%],F:0.4%,M:0.8%,n:1614 |
| **Number of annotated genes** | 32,751 | 41,683 | 39,383 | 35,354 | 24,965 | 25,541 | 33,351 | 31,523 | 28,982 | 32,187 | 31,569 | 31,425 |
| **Percentage of repetitive elements** | 63.78% | 63.32% | 63.15% | 64.55% | 63.31% | 59.57% | 69.67% | 56.72% | 54.69% | 68.31% | 67.97% | 61.00% |

*Explanation of the notations: C: Complete BUSCOs (C); S: Complete and single-copy BUSCOs (S); D: Complete and duplicated BUSCOs (D); F: Fragmented BUSCOs (F); M: Missing BUSCOs (M); n: Total BUSCO groups searched

**Table S2. Properties of *R*-locus regions for *Wachendorfia paniculata*, *Wachendorfia thyrsiflora* and *Barberetta aurea***

| **Species** | ***Wachendorfia paniculata*** | ***Wachendorfia thyrsiflora*** | ***Barberetta aurea*** |
|---|---|---|---|
| Coordinates | ptg000001l:23,250,000-23,450,000 | ptg000020l:11,400,000-11,600,000 | h2tg000059l:1,150,000-1,250,000 |
| Size | 200 kb | 200 kb | 100 kb |
| Repeat content | 80.590% | 86.828% | 86.013% |

**Table S3. PCR genotyping of wild-type L- and R-morph plants and of homostylous mutants**
Results of PCR-genotyping the indicated samples for the indicated amplicons.

| Population | Species | Phenotypes found | Phenotype Counts | | Positive for *YUC-P1* | Positive for *YUC-R* (CDS) | Positive for *MIR156* | Positive for full *YUC-R* deletion | Positive for partial YUC-R1 deletion | Positive for wt *YUC-R* allele | Positive for full *MIR156* deletion |
|---|---|---|---|---|---|---|---|---|---|---|---|
| | | | 2023 | 2024 | | | | | | | |
| Campsbay | *W. paniculata* | Left | | | 110/110 | 0/110 | 0/110 | | | | |
| | | Right | | | 110/110 | 110/110 | 110/110 | | | | |
| Redelinghuys | *W. multiflora* | Left | 0 | 5 | 3/3 | 0/3 | 0/3 | 0/3 | 0/3 | 0/3 | 0/3 |
| | | Right | 0 | 10 | 8/8 | 8/8 | 8/8 | 2/8 | 1/8 | 8/8 | 0/8 |
| | | Right-Homostyle | 16 | 34 | 26/26 | 13/26 | 26/26 | 13/26 | 13/26 | 0/26 | 0/26 |
| Rondebosch Common | *W. brachyandra* | Left | >200 | >200 | 2/2 | 0/2 | 0/2 | 0/2 | 0/2 | 0/2 | 0/2 |
| | | Right | >200 | >200 | 3/3 | 3/3 | 3/3 | 0/3 | 0/3 | 3/3 | 0/3 |
| | | Left-Homostyle | 1 | 0 | 1/1 | 1/1 | 0/1 | 0/1 | 0/1 | 1/1 | 1/1 |
| Farm 215 | *W. brachyandra* | Left | n/a | 11 | 5/5 | 0/5 | 0/5 | 0/5 | 0/5 | 0/5 | 0/5 |
| | | Left-Homostyle | n/a | 30 | 5/5 | 5/5 | 0/5 | 0/5 | 0/5 | 5/5 | 5/5 |
| | *W. paniculata* | Left | n/a | 15 | 4/4 | 0/4 | 0/4 | 0/4 | 0/4 | 0/4 | 0/4 |
| | | Right-Homostyle | n/a | 15 | 5/5 | 5/5 | 5/5 | 0/5 | 5/5 | 0/5 | 0/5 |
| Groot Hagelkraal | *W. paniculata* | Right-Homostyle | n/a | >200 | 8/8 | 0/8 | 8/8 | 8/8 | 8/8 | 0/8 | 0/8 |

**Table S4: Coverage analysis of *R*-locus genes in pools and individual homostylous mutant samples**

Coverage of the two *R*-locus genes was normalized to the whole-genome background.

| Sample | Species | Location | Phenotype | *YUC-R* | *MIR156-R* | Interpretation for *R*-locus genotype |
|---|---|---|---|---|---|---|
| | *W. brachyandra* | Kenilworth Race Course | L-morph (pool) | 0.0465349 | 0.028075 | *L/L* |
| | *W. brachyandra* | Kenilworth Race Course | R-morph (pool) | 0.45676 | 0.346539 | mostly *R/L* |
| | *W. multiflora* | Langebaan | L-morph (pool) | 0.0011069 | 0.000221 | *L/L* |
| | *W. multiflora* | Langebaan | R-morph (pool) | 0.904049 | 0.567009 | mostly *R/L* |
| | *W. paniculata* | Campsbay firebreak pop. 1 | L-morph (pool) | 0.0002902 | 0 | *L/L* |
| | *W. paniculata* | Campsbay firebreak pop. 1 | R-morph (pool) | 0.58158 | 0.569928 | mostly *R/L* |
| | *W. paniculata* | Campsbay firebreak pop. 2 | L-morph (pool) | 0.0002161 | 0.000432 | *L/L* |
| | *W. paniculata* | Campsbay firebreak pop. 2 | R-morph (pool) | 0.554299 | 0.711284 | mostly *R/L* |
| | *W. thyrsiflora* | Botriver lagoon | L-morph (pool) | 0 | 0 | *L/L* |
| | *W. thyrsiflora* | Botriver lagoon | R-morph (pool) | 0.668558 | 0.376155 | mostly *R/L* |
| | *B. aurea* | Ngeli Forest | L-morph (pool) | 0.0012708 | 0 | *L/L* |
| | *B. aurea* | Ngeli Forest | R-morph (pool) | 0.657766 | 0.447124 | mostly *R/L* |
| Wb_FA_L1 | *W. brachyandra* | Farm 215 | L-morph | 0 | 0 | *L/L* |
| Wb_FA_L2 | *W. brachyandra* | Farm 215 | L-morph | 0 | 0 | *L/L* |
| Wb_FA_Lh1 | *W. brachyandra* | Farm 215 | Left-homostylous | 0.943607 | 0 | $R^m/R^m$ |
| Wb_FA_Lh2 | *W. brachyandra* | Farm 215 | Left-homostylous | 0.87011 | 0 | $R^m/R^m$ |
| Wb_RO_Mut1 | *W. brachyandra* | Rondebosch Common | Left-homostylous | 0.465967 | 0 | $R^m/L$ |
| Wb_RO_WtR1 | *W. brachyandra* | Rondebosch Common | R-morph | 0.364355 | 0.454295 | *R/L* |
| Wm_RE_L1 | *W. multiflora* | Redelinghuys | L-morph | 0 | 0.000658 | *L/L* |
| Wm_RE_L2 | *W. multiflora* | Redelinghuys | L-morph | 0 | 0 | *L/L* |
| Wm_RE_R1 | *W. multiflora* | Redelinghuys | R-morph | 1.06484 | 0.932645 | $R^m/R^m$ |
| Wm_RE_R2 | *W. multiflora* | Redelinghuys | R-morph | 1.08025 | 0.936683 | $R^m/R^m$ |
| Wm_RE_Rh1 | *W. multiflora* | Redelinghuys | Right-homostylous | 0.843579 | 0.882236 | $R^m/R^m$ |
| Wm_RE_Rh2 | *W. multiflora* | Redelinghuys | Right-homostylous | 0.801143 | 0.750708 | $R^m/R^m$ |
| Wm_RE_Rh3 | *W. multiflora* | Redelinghuys | Right-homostylous | 0 | 0.923807 | $R^m/R^m$ |
| Wm_RE_Rh4 | *W. multiflora* | Redelinghuys | Right-homostylous | 0 | 0.838316 | $R^m/R^m$ |

| | | | | | | |
|---|---|---|---|---|---|---|
| Wm_RE_Rh5 | *W. multiflora* | Redelinghuys | Right-homostylous | 0 | 0.817914 | $R^m/R^m$ |
| Wm_RE_Rh6 | *W. multiflora* | Redelinghuys | Right-homostylous | 0 | 1.02542 | $R^m/R^m$ |
| Wm_KB_R1 | *W. multiflora* | Koeberg | R-morph | 0.422001 | 0.559107 | *R/L* |
| Wm_KB_R2 | *W. multiflora* | Koeberg | R-morph | 0.908003 | 0.950927 | *R/R* |
| Wp_FA_L1 | *W. paniculata* | Farm 215 | L-morph | 0 | 0 | *L/L* |
| Wp_FA_L2 | *W. paniculata* | Farm 215 | L-morph | 0 | 0 | *L/L* |
| Wp_FA_Rh1 | *W. paniculata* | Farm 215 | Right-homostylous | 0.874659 | 0.890659 | $R^m/R^m$ |
| Wp_FA_Rh2 | *W. paniculata* | Farm 215 | Right-homostylous | 0.80767 | 0.851548 | $R^m/R^m$ |
| Wp_GH_Rh1 | *W. paniculata* | Groot Hagelkraal | Right-homostylous | 0.0320318 | 0.987267 | $R^m/R^m$ |
| Wp_GH_Rh2 | *W. paniculata* | Groot Hagelkraal | Right-homostylous | 0.0194941 | 1.10741 | $R^m/R^m$ |

**Table S5: Primers used in this study**

| Species | Population | Target | Primer | Sequence | amplicon size (bp) | Cycling conditions |
|---|---|---|---|---|---|---|
| All *Wachendorfia* species | All | miR156-R conserved sequence | W_miR156-R_F | ATCTCCGCAACCCAAACCTC | 435 | 94°C 3 min, (94°C 10 s, 56°C 20 s, 72°C 30 s) 40 cycles, 72°C 5 min |
| | | | W_miR156-R_R | GGCATGTCAATCCAATATCATCCG | | |
| | | Within conserved YUC-R sequence | W_YUC-R_F | CCTCTACAGTCCCATCCACAA | 313 | |
| | | | W_YUC-R_R | TGCAAGTGATTTCCTGTGAGA | | |
| | | Within conserved YUC-R paralogue gene, YUC-P1, sequence | W_YUC-P1_F | CGTGGTCTCATTTAGCAGCA | ~200 | |
| | | | W_YUC-P1_R | TGGGCGAACCTGAAAATAAC | | |
| *W. brachyandra* | Farm 215 | 5' Flank of miR156 deletion | Wb_F215_del_F | TTTCTTCCGTTCCTTTGGGG | 440 | 98°C 3 min, (98°C 10 s, 63°C 20 s, 72°C 30 s )40 cycles, 72°C 5 min |
| | | 3' Flank of miR156 deletion | Wb_F215_del_R | AGACGAAATCCAAGAACTCTCA | | |
| *W. brachyandra* | Rondebosch Common | 5' Flank of miR156 deletion | Wb_Ron_del_F | GTAACTTGGGACGACATGCT | 600 | |
| | | 3' Flank of miR156 deletion | Wb_Ron_del_R | AGGGTTCGCTCACTGAATCT | | |
| *W. paniculata* | Groot Hagelkraal | 5' Flank of YUC-R deletion | Wp_GH_del_F | CCCAGTAAGCCATGTCTCGA | 427 | 98°C 3 min, (98°C 10 s, 66°C 20 s, 72°C 30 s)40 cycles, 72°C 5 min |
| | | 3' Flank of YUC-R deletion | Wp_GH_del_R | GCTTCATCTCCATGGCGATC | | |
| *W. paniculata* | Farm 215 | Within the small deletion/insertion in right homostyle plants | Wp_F215_smalldel_ ins_F | TGAGGCAATATCCGAACCCG | 340 | 94°C 3 min, (94°C 10 s, 58°C 20 s, 72°C 30 s ) 40 cycles, 72°C 5 min |
| *W. paniculata*, *W. multiflora* | Farm 215 and Redelinghuys | Downstream to YUC-R coding sequence | Wp_F215_Wm_RH1_ no_smalldel_F | AGAGATCATGAGAACGGTCAGA | 789 | |
| | | within YUC-R intron3 (between exon3 and exon4) | Wp_F215_Wm_RH1_ smalldel_R | TGCGCTTTCCATTGAACTGT | | |
| *W. multiflora* | Redelinghuys | Within the small deletion/insertion in right homostyle plants | Wm_RH1_smalldel_ ins_F | AGAGGAGTACGCCATGCTAA | 484 | 94°C 3 min, (94°C 10 s, 58°C 20 s, 72°C 30 s) 30 cycles, 72°C 5 min |
| | | Within the large deletion/insertion in | Wm_RH2_largedel_ ins_F | TCAAACAGGAGGCAGATGGA | 317 | |

| | | | | | | |
|---|---|---|---|---|---|---|
| | | right homostyle plants | | | | 94°C 3 min, (94°C 10 s, 52°C 20 s, 72°C 30 s )40 cycles, 72°C 5 min |
| | | Left to the 3' breakpoint of the large deletion | Wm_RH2_no_largedel_F | TCCATATTAGTCCAGTCCAGTGA | | |
| | | Right to the 3' breakpoint of the large deletion | Wm_RH2_largedel_R | TGATTGGACACCTAGATGCAAG | 759 | |

**Table S6: Summary of populations of *Wachendorfia* species, *B. aurea* and *D. ixioides* used in this study.**

| Species | Study | Location | Permission | Permit | GPS co-ordinates |
|---|---|---|---|---|---|
| *W. paniculata* | Genome assembly | Campsbay firebreak | SANparks | CRC/2023-2024/009--2023/V1 | 33°57'36.93"S 18°23'14.43"E |
| *W. paniculata* | 100 left vs 100 right; RNA-seq | Campsbay firebreak | SANparks | CRC/2023-2024/009--2023/V1 | 33°57'25.83"S 18°23'19.06"E |
| *W. paniculata* | Right homostyle | Farm 215, Private Nature Reserve | Martin Groos, Cape Nature | CN35-87-25844 | 34°34'18.98"S 19°30'7.53"E |
| *W. paniculata* | Right homostyle | Groot Hagelkraal | Eskom, Cape Nature | CN35-87-25844 | 34°40'41.43"S 19°34'23.57"E |
| *W. multiflora* | 50 left vs 50 right | Langebaan | SANparks | CRC/2023-2024/009--2023/V1 | 33° 6'16.83"S 18° 2'48.01"E |
| *W. multiflora* | RNA-seq | West Coast National Park | SANparks | CRC/2023-2024/009--2023/V1 | 33° 6'28.00"S 18° 0'20.93"E |
| *W. multiflora* | Right homostyle | Redelinghuys | Cape Nature and Redelinghuys municipality | CN35-87-25844 | 32°29'5.99"S 18°32'13.46"E |
| *W. thyrsiflora* | Genome assembly | Botanical Garden of the University of Potsdam | | | 52°24'12.5"N 13°01'29.0"E |
| *W. thyrsiflora* | 50 left vs 50 right | Botriver lagoon | Riki Erwee, Cape Nature | CN35-87-25844 | 34°18'9.32"S 19° 8'42.84"E |
| *W. brachyandra* | 50 left vs 50 right | Kenilworth Race Course | Cape Nature | CN35-87-25844 | 33°59'55.80"S 18°29'1.49"E |
| *W. brachyandra* | Left homostyle mutant | Rondebosch Common | Friends of the Rondebosch Common, Cape Nature | CN35-87-25844 | 33°57'20.53"S 18°29'7.16"E |
| *W. brachyandra* | Left homostyle mutant | Farm 215, Private Nature Reserve | Martin Groos, Cape Nature | | 34°34'14.39"S 19°30'19.21"E |
| *B. aurea* | Genome assembly | Rockwood, Karkloof Nature Reserve | Sue Stradford, Private landowner | Private landowner permission | 29°19'5.24"S 30°15'12.49"E |
| *B. aurea* | 40 left vs 40 right | Ngeli Forest | Merensky timber | Private landowner permission | 30°34'59.10"S 29°38'54.83" |
| *D. ixioides* | Genome assembly | Die Drift farm | Stephen and Carolyn Viljoen, Cape Nature | CN35-87-25844 | 32°41'29.71"S 19°16'42.81"E |

**Movie S1. Style deflection in a dissected bud of *Wachendorfia paniculata***

Time-lapse movie of dissected buds of *W. paniculata*. The movie was recorded over 53 hours. Sepals and petals were removed. Jumps in the movie result from power-outages during recording.

**Movie S2: Style deflection in dissected bud of *Wachendorfia paniculata* with and without rotation by 180°**

Time-lapse movie of dissected buds of *W. paniculata*. The movie was recorded over 19 hours. Sepals, petals and the two adaxial stamens were removed. Both buds were from the same R-morph plant. The bud on the left was cultured in its normal orientation with the abaxial stamen below the ovary, while the bud on the left had been rotated by 180°, such that the abaxial stamen was now above the ovary.

**Movie S3: Model output with both twist and differential elongation**

Output of the biophysical model of *Wachendorfia* style deflection with both twist and differential elongation. The red nodes indicate the abaxial side with stronger elongation. The plane forming the crossbar of the T when viewed from above is the plane tangent to the axis of greatest elongation, relative to which deflection of the arrow representing the long axis of the style was measured. The plane forming the stem of the T represents the flower midline, i.e. it is perpendicular to the base of the style and runs through the basal nodes representing the adaxial and abaxial sides of the flower.

**Movie S4: Model output with only twist**

Same as in Movie S3, but here the model was run with only twist, but without differential elongation.

**Movie S5: Model output with only differential elongation**

Same as in Movie S3, but here the model was run with only differential elongation, but without twist.

**Data S1: MapMan enrichment analysis of differentially expressed genes**

**Data S2: Genespace output for synteny analyses**

**Data S3: Prediction of *miR156-5p* targets in *W. paniculata* style and stamen transcriptome**

**Data S4: Sanger sequences of deletions from homostylous mutants**

**Data S5: Sequences used in phylogenetic reconstruction**

**Data S6: Mathematica Notebook**
