## Supplementary material for "Supergene control of chiral development in mirror-image flowers": Data S5

>Wb_YUC-R

ATGGCTAGAAAGAGATTTCAAGACACATTGCTTCATCTCCATGGCGATCATCAGCAGCATCATGACGATGACGTTGATGCAGGAGTTGAGACTTACCATCAACTCTGTGATGCTTATTGTAGAGTTGCCAAAGATTCCAAAGCAGTTGCTGTTCAAAATGGCGACCAGTCCGTCATGTTCCATGGGCCAATCATTGTAGGCGCTGGACCGTCAGGACTAGCTGTTGCCGCCTGCCTCAAGGACAAGGGCATCCCGTCGATGGTCATCGAACGGTCGGACTGCATCGCTTCCCTGTGGCAGCTGAAGACCTACGACCGTTTATGCCTCCATCTGCCAAAGCATTTCTGCGAGCTTCCACTCATGCCGTTCCCAGACGACTTCCCCAGGTATCCTACAAAAAACCAGTTCATCTCCTACCTCAAGGATTACGCCCGCCGCTTTAAAATAAAGCCGGTGTTCAACCAGACAGTCAGCAGTGCAGAGTATGATGATCGGACGCAGTTATGGAGGGTGAAGACATTGGCAAAAGAGTATGTCTGCCCTTGGTTGGTAGTGGCTACCGGAGAGAATGCGGAGGAAGTGGTGCCAAAGTTGGAAGGCATGGCGGAGTTTAAGGGCCCGATCATCCACACGAGTTTGTTCAAGTCCGGCAAGGTGTTCAATGGGAAAAAAGTGCTGGTGGTGGGCTGTGGCAATTCTGGGACGGATATTAGTTTGGACCTCTACAATCACAATGCTCAACCCTACCTCGTCGTAAGGGATGGGGTGCATATCTTGCCCAAAGAAATCCTCGGTCGGTCAACCTTTTCGCTTTGCATGTGGCTCCTTAAATGGTTGCCTTTATCAGCAGTGGATCGCTTCCTGCTGCTTATCTGCAGGATCTTGGTCGGCAACACAGAGCAATGTGGCATAAGCAGACCGGAGTTAGGGCCACTTACGCTCAAGTCTCTCTCTGGCAAGACTCCAGTCCTTGATGTTGGAACCCTTGCCAGGATCCAGTCAGGTGACATCAAGGTACGGCCAGGAATAAAGAGATTAACAAGGGATGGAGCTGAATTTGTGGATGGGACTGTAGAGGATTTTGATGCAATCATATTGGCAACTGGATACAAAAGCAATGTACTATCTTGGTTAAAGGATCCAGAGTTCTTCTCAGATGAAAATGGATTGCCAAAGAGGTCATTTCCCAACGGGTGCAAAGGCAAGCGAGGCCTATACGCTGTCGGTTTTGCGCAAAGAGGCCTGATGGGGATTAGGATCGACGCAAGAAGGGTTGCAAATGACATCCATCAATGCTGGGAGTTTGAAGGACATGCTATCCACAAACCTTTCTAG

>Wm_YUC-R

ATGGCTAGAAAGAGATTTCAAGACACATTGCTTCATCTCCATGGCGATAATCAGCAGCATCATGACGATGACGTTGATGCAGGAGTTGAGACTTACCATCAACTCTGTGATGCTTATTGTAGAGTTGCCAAAGATTCCAAAGCAGTTGCTGTTCAAAATGGCGACCAGTCCGTCATGTTCCATGGACCAATCATTGTCGGCGCTGGACCGTCAGGACTAGCTGTTGCCGCCTGCCTCAAGGACAAGGGCATCCCGTCGATGGTCATCGAACGGTCGGACTGCATCGCTTCCCTGTGGCAGCTGAAGACCTACGACCGTTTATGCCTCCATCTGCCAAAGCATTTCTGCGAGCTTCCACTCATGCCGTTCCCAGACGACTTCCCCAGGTATCCTACAAAAAACCAGTTCATCTCCTACCTCAAGGATTACGCCCGCCGCTTTGAAATAAAGCCGGTGTTCAACCAGACAGTCAGCAGTGCAGAGTATGATGATCGGACGCAGTTGTGGAGGGTTAAGACATTGGCAAAAGAGTATGTCTGCCCTTGGTTGGTAGTGGCTACCGGAGAGAATGCGGAGGAAGTGGTGCCAAAGTTGGAAGGCATGCCGGAGTTTAATGGCCCGATCATCCACACGAGTTTGTTCAAGTCCGGCAAGGTGTTCAATGGGAAAAAAGTGCTGGTGGTGGGCTGTGGCAATTCTGGGACGGATATTAGCTTGGACCTCTACAATCACAATGCTCAAACCTACCTCGTCGTAAGGGATGGGGTGCATATCTTGCCCAAAGAAATCCTCGGTCGGTCGACCTTTTCGCTTTGCATGTGGCTCCTTAAATGGTTGCCTTTATCAACAGTGGATCGCTTCCTGCTGCTTATCTGCAGGATCTTGGTCGGCAACACAGAGCAATGTGGCATAAGCAGACCGGAGTTAGGGCCACTCACGCTCAAGTCTCTCTCTGGCAAGACTCCAGTCCTTGATGTTGGAACCCTTGCCAGGATCAAGTCAGGTGACATCAAGGTTCGCCCAGGAATAAAGAGATTAACAAGGGATGGAGCTGAATTTGTGGATGGGACTGTAGAGGATTTTGATGCAATCATATTGGCAACTGGATACAAAAGCAATGTACTATCTTGGTTAAAGGATCCAGAGTTCTTCTCAGATGAAAATGGATTGCCAAAGAGGTCATTTCCCAACGGGTGCAAAGGCAAGCGAGGCCTATACGCTGTCGGTTTTGCGCAAAGAGGCCTGATGGGGATTAGGATCGACGCAAGAAGGGTTGCAAATGACATCCATCAATGCTGGAAGTTTGAAGGACATGCTATCCACAAACCTTTCTAG

>Wp_YUC-R

ATGGCTAGAAAGAGATTTCAAGACACATTGCTTCATCTCCATGGCGATCATCAGCAGCATCATGACGATGACGTTGATGCAGGAGTTGAGACTTACCATCAACTCTGTGATGCTTATTGTAGAGTTGCCAAAGATTCCAAAGCAGTTGCTGTTCAAAATGGCGACCAGTCCGTCATGTTCCATGGGCCAATCATTGTCGGCGCTGGACCGTCAGGACTAGCTGTTGCCGCCTGCCTCAAGGACAAGGGCATCCCGTCGATGGTCATCGAACGGTCGGACTGCATCGCTTCCCTGTGGCAGCTGAAGACCTACGACCGTTTATGCCTCCATCTGCCAAAGCATTTCTGCGAGCTTCCACTCATGCCGTTCCCAGACGACTTCCCCAGGTATCCTACAAAAAACCAGTTCATCTCCTACCTCAAGGATTACGCCCGCCGCTTTGAAATAAAGCCGGTGTTCAACCAGACAGTCAGCAGTGCAGAGTATGATGATCGGACGCAGTTATGGAGGGTGAAGACATTGGCAAAAGAGTATGTCTGCCCTTGGTTGGTAGTGGCTACCGGAGAGAATGCGGAGGAAGTGGTGCCAAAGTTGGAAGGCATGGCGGAGTTTAAGGGCCCGATCATCCACACGAGTTTGTTCAAGTCCGGCAAGGTGTTCAATGGGAAAAAAGTGCTGGTGGTGGGCTGTGGCAATTCTGGGACGGATATTAGTTTGGACCTCTACAATCACAATGCTCAACCCTACCTCGTCGTAAGGGATGGGGTGCATATCTTGCCCAAAGAAATCCTCGGTCGGTCAACCTTTTCGCTTTGCATGTGGCTCCTTAAATGGTTGCCTTTATCAGCAGTGGATCGCTTCCTGCTGCTTATCTGCAGGATCTTGGTCGGCAACACAGAGCAATGTGGCATAAGTAGACCGGAGTTAGGGCCACTCACGCTCAAGTCTCTCTCTGGCAAGACTCCAGTCCTTGATGTTGGAACCCTTGCCAGGATCCAGTCAGGTGACATCAAGGTACGGCCAGGAATAAAGAGATTAACAAGGGATGGAGCTGAATTTGTGGATGGGACTGTAGAGGATTTTGATGCAATCATATTGGCAACTGGATACAAAAGCAATGTACTATCTTGGTTAAAGGATCCAGAGTTCTTCTCAGATGAAAATGGATCGCCAAAGAGGTCATTTCCCAACGGGTGCAAAGGCAAGCGAGGCCTATACGCTGTCGGTTTTGCGCAAAGAGGCCTGATGGGGATTAGGATCGACGCAAGAAGGGTTGCAAATGACATCCATCAATGCTGGAAGTTTGAAGGACATGCTATCCACAAACCTTTCTAG

>Wt_YUC-R

ATGGCTAGAAAGAGATTTCAAGACACATTGCTTCATCTCCATGGCGATCATCAGCAGCATCATGACGATGACGTTGATGCAGGAGTTGAGACTTACCATCAACTCTGTGATGCTTATTGTAGAGTTGCCAAAGATTCCAAAGCAGTTGCTGTTCAAAATGGCGACCAGTCCGTCATGTTCCATGGGCCAATCATTGTCGGCGCTGGACCGTCAGGACTAGCTGTTGCCGCCTGCCTCAAGGACAAGGGCATCCCGTCGATGGTCATCGAACGGTCGGACTGCATCGCTTCCCTGTGGCAGCTGAAGACCTACGACCGTTTATGCCTCCATCTGCCAAAGCATTTCTGCGAGCTTCCACTCATGCCGTTCCCAGACGACTTCCCCAGGTATCCTACAAAAAACCAGTTCATCTCCTACCTCAAGGATTACGCCCGCCGCTTTGAAATAAAGCCGGTGTTCAACCAGACAGTCAGCAGTGCAGAGTATGATGATCGGACGCAGTTGTGGAGGGTGAAGACATTGGCAAAAGAGTATGTCTGCCCTTGGTTGGTAGTGGCTACCGGAGAGAATGCGGAGGAAGTGGTGCCAAAGTTGGAAGGCATGGCGGAGTTTAAGGGCCCGATCATCCACACGAGTTTGTTCAAGTCCGGCAAGGTGTTCAACGGGAAAAAAGTGCTGGTGGTGGGCTGTGGCAATTCTGGGACGGATATTAGTTTGGACCTCTACAATCACAATGCTCAAACCTACCTCGTCGTAAGGGATGGGGTGCATATCTTGCCCAAAGAAATCCTCGGTCGATCAACCTTTTCGCTTTGCATGTGGCTCCTTAAATGGTTGCCTTTATCAGCAGTGGATCGCTTCCTGCTGCTTATCTGCAGGATCTTGGTCGGCAACACAGAGCAATGTGGCATAAGTAGACCGGAGTTAGGGCCACTCACGCTCAAGTCTCTCTCTGGCAAGACTCCAGTCCTTGATGTTGGAACCCTTGCCAGGATCCAGTCAGGTGACATAAAGGTTCGGCCAGGAATAAAGAGATTAACAAGGGATGGAGCTGAATTTGTGGATGGAACTGTAGAGGATTTTGATGCAATCATATTGGCAACTGGATACAAAAGCAATGTACTATCTTGGTTAAAGGATCCAGAGTTCTTCTCAGATGAAAATGGATTGCCAAAGAGGTCATTTCCCAACGGGTGCAAAGGCAAGCGAGGCCTATACGCTGTCGGTTTTGCGCAAAGAGGCCTGATGGGGATTAGGATCGACGCAAGAAGGGTTGCAAATGACATCCATCAATGCTGGAAGTCTGAAGGACATGCTATCCAAAAACCTTTCTAG

>Ba_Karkloof_YUC-R

ATGAGCCCTATCTTCCAACAAAATGCCTTCAGAGGTTTTGCTAAATGTATAAACGAAGATGTTGAAGCTTATTGTAGAGTTGCCAAAGATTCCATAGCAGTTTCTGTTCATAAATCCCCTCAGTCTGTCATGATCCATGGGCCGATCATCGTTGGCGCTGGACCATCAGGGCTCGCCGTCGCCGCTTGCCTCAAGGACAAGGGGATCCCGTCGACGGTCATTGAACGGTCCGACTGCGTCGCTTCAATGTGGCAGCTGAAAACTTACGACCGTTTATGCCTCCATCTGCCAAAGCATTTCTGCGAGCTTCCCCTCATGGAGTTTCCAGCCTACTTCCCCAGGTATCCTACAAAAGAACAGTTTGTATCTTACCTGAAGGATTATGCCTGCCGGTTCGAAATAAAGCCAGTGTTCAATCAGACAGTCTGCAGTGCAGAGTATAATGAACTGATGCAGCTGTGGATGGTGAAGACATCCGCGATGCAGTATCTCAGCCGTTGGTTGGTAGTGGCTACCGGAGAGAATGCGGAGGAAGTGGTGCCAGAGTTGGATGGCATGGTGGACTTTAAGGGCTTGATCATCCACACGAGTATGTTCAAGTCCGGTAAGGTGTTCAATGGAAAAAAAGTTCTGGTAGTGGGCTGTGGCAATTCTGGGATGGATATTAGCTTGGACCTCTATAATCACAATGCTCAAGCTCACATTGTCGTTAGGGATGGGGTGCATATCTTGCCGAAAGAAATCCTCGGTCAATCCACCTTTGCCCTTTGCACGTGGCTCCTTAAATGGTTTCCTTTATCAGCAGTGGATCGCTTCCTGCTGCTCATCTGTAGGATCTTGGTCGGCAACACTGAGCAATGTGGCATAAGCAGACCGGAGTTAGGGCCACTCACGCTTAAGTCTCTCTCCGGCAAGACTCCAGTCATTGATGTTGGAGCCCTAGCCAGGATCCAGTCCGGTGACATCAAGGTTCGGCCAGGAATAAAGAGATTAACAAGAGATGGAGCTGAATTTGTGGATGGGAAGATAGAGGATTTTGATGCAATCATATTGGCAACTGGTTACAAAAGCAATGTACTATCTTGGCTAAAGGAGCCAGAGCTCTCTGATACAAATGGATTGCCAAAGAGGCCATTTCATGACGGTTGGAAAGGCAAGAGAGGCCTATATGCTGTTGGTTTTGCGCAAAGAGGCCTAATGGGGATTAAGATTGACGCAAGAAGGGTTGTCAATGACATCCATAAATGCTGGAAGGCTGAAGGAGGCTCTATCATCGAACCCTTCTAG

>Ba_Ngeli_YUC-R

ATGAGCCCTATCTTCAAACAAAATGCCTTCAGAGGTTTTGCTAAATGTATAAACGAAGATGTTGAAGCTTATTGTAGAGTTGCCAAAGATTCCATAGCAGTTGCTGTTCATAAATCCCCTCAGTCTGTCATGATCCATGGGCCGATCATCGTTGGTGCTGGACCATCAGGACTCGCCGTCGCCGCTTGCCTCAAGGACAAGGGGATCCCGTCGATGGTCATTGAGCGGTCCGACTGCGTCGCTTCAATGTGGCAGCTGAAAACTTACGACCGTTTATGCCTCCATCTGCCAAAGCATTTCTGCGAGCTTCCCCTCATGGAGTTTCCAGCCTACTACCCCAGGTATCCTACAAAAGAACAGTTTGTATCTTACCTGAAGGATTATGCCTGCCGGTTCGAAATAAAGCCAGTGTTCAATCAGACAGTCTGCAGTGCTGAGTATGATGAACTGATGCAGCTGTGGATGGTGAAGACATCCGCGATGCAGTATCTCAGCCGTTGGTTGGTAGTGGCTACCGGAGAGAATGCGGAGGAAGTGGTGCCGGAGTTGGATGGCATGGTGGACTTTAAGGGCTTGATCATCCACACGAGTATGTTCAAGTCCGGTAAGGTGTTCAATGGAAAAAAAGTTCTGGTAGTGGGCTGTGGCAATTCTGGGATGGATATTAGCTTGGACCTCTATAATCACAATGCTCAAACTCACATTGTCGTTAGGGATGGGGTGCATATCTTGCCGAAAGAAATCCTCGGTCAATCCACCTTTGCCCTTTGCACGTGGCTACTTAAATGGTTTCCTTTATCAGCAGTGGATCGCTTCCTGCTGCTCATCTGTAGGATCTTGGTCGGCAACACTGAGCAATGTGGCATAAGCAGACCGGAGTTAGGGCCACTCACGCTTAAGTCTCTCTCCGGCAAGACTCCAGTCATTGATGTTGGAGCCCTAGCCAGGATCCAGTCCGGTGACATCAAGGTTCGGCCAGGAATAAAGAGATTAACAAGAGATGGAGCTGAATTTGTGGATGGGAAGATAGAGGATTTTGATGCAATCATATTGGCAACTGGTTACAAAAGCAATGTACTATCTTGGTTAAAGGAGCCAGAGTTCTCTGATACAAATGGATCGCCAAAGAGGCCATTTCATGACGGTTGGAAAGGCAAGAGAGGCCTATATGCTGTTGGTTTTGCGCAAAGAGGCCTAATGGGGATTAAGATTGACGCAAGAAGGGTTGCCAATGACATACATAAATGCTGGAAGGCTGAAGGAGGCTCTATCATCGAACCTTTCTAG

>Wb_YUC-P1

ATGGCCGGAAAGAGATTCCACGATCCTTCGCTCCAACTCCACGGCCACCGCCAATATCAACACCCCGCCGCCTCTGAGGCCGCCACTGACGATAAAGCTGAGCAGTTTATCTGGTTCCCGGGGCCTATCATTGTTGGCGCGGGGCCGTCGGGGCTAGCCGTCGCCGCCTGCCTGAAGGAGAAAGGTATCCCGTCGATGGTCATCGAGCGGTCCGACTGCATCGCCTCCCTCTGGCAGGTGAAGACCTACGATCGTCTCTCCCTCCATCTGCCGAAGCACTTCTGCGAGCTTCCGCTCATGCCATTCCCGGCCTCCTTCCCCCAATATCCCACGAAGCAGCAGTTCGTGGCCTATCTCGAGACCTATGCACGGCACTTCGACATACGACCGGTGTTCAATCAGACCGTCGTCAATGCTGAGTACGATGATCGGATGCGATCTTGGATGGTGCAGACAACCAAGACGGCGCCCACGGAGGGGAGGACGGCGACTAAAGGGTACATCAGCCGGTGGTTGGTGGTTGCTACCGGGGAGAATTCGGAGGAGGTGGTGCCAGAGATGGAAGGGATGACGGAGTTCAAAGGACCGATCATCCATACAAGTATGTTCAAGTCAGGCGAGGTGTTCGAGGGGAAGCCAGTGCTGGTGGTGGGCTGCGGCAATTCAGGGATGGAGATCAGCTTGGACCTTCACAACCACAATGCTCGTCCTCACCTCGTAGTAAGAGATACGGTGCACATATTGCCGAGAGAAATCATGGGCCGATCGACTTTTGGGATGAGCATGTGGCTGCTCAAATGGCTGCCTCTGTCTGTGGTGGACCGCGTCCTTCTGCTCGTCTCCAGGATCGTGCTCGGCGACACGGAGCAATGCGGCATAAGCAGGCCTCGGCTGGGGCCACTCGAGCTTAAGTCTCTCTCCGGCAAGACTCCTGTCCTCGATGTCGGAACCCTAGCCAAGATTCAGTCCGGTGACATTAAGGTTCGCCCAGGAATAAGTCGATTGACTAGAGATGGGGCTGAGTTTGTCGATGGAAGGATCGAAGATTTTGATGCAATCATATTGGCAACTGGTTACAAAAGCAATGTACTCTCTTGGTTAAAGGTAGGTGAATTTTTCTCTGATAAGAGCGGATTGCCAAGGAGACCATTTCCCAACAGTTGGAAGGGCGAGCGAGGCCTCTACGCAGTTGGTTTCACGCAACGAGGCCTAATGGGGACTAAGATCGACTCGAGGAGGATTGTGCATGACATAGAGCAATGTTGGAAGGCCTCAAGGACTATGTAA

>Wm_YUC-P1

ATGGCCGGAAAGAGATTCCACGATCCTTCGCTCCAACTCCACGGCCACCGCCAATATCAACACCCCGCCGCCTCTGCGGCCGCCACTGACGATAAAGCTGAGCAGTTTATCTGGTTCCCGGGGCCTATTATTGTTGGCGCGGGGCCGTCGGGGCTAGCCGTCGCCGCTTGCCTGAAGGAGAAAGGTATCCCGTCGATGGTCATCGAGCGGTCCGACTGCATCGCCTCCCTCTGGCAGGTGAAGACCTACGATCGTCTCTCCCTCCATCTGCCGAAGCACTTCTGCGAGCTTCCGCTCATGCCATTCCCGGCCTCCTTCCCCCAATATCCCACGAAGCAGCAGTTCGTGGCCTATCTCGAGACCTATGCACGGCACTTCGACATACGACCGGTGTTCAATCAGACCGTTGTCAATGCTGAGTACGATGATCGGATGCGATCTTGGATGGTGCAGACAACCAAGACGGCGCCCACGGAGGGGAGGACGGCGACTAAGGGGTACATCAGCCGGTGGTTGGTGGTTGCTACCGGGGAGAATTCGGAGGAGGTGGTGCCAGAGATGGAAGGGATGACGGAGTTCAAAGGACCGATCATCCATACGAGTATGTTCAAGTCAGGCGAGGTGTTCGAGGGGAAGCCAGTGCTGGTGGTGGGCTGCGGCAATTCAGGGATGGAGATCAGCTTGGACCTTCACAACCACAATGCTCGTCCTCACCTCGTAGTAAGAGATACGGTGCACATATTGCCGAGAGAAATCATGGGCCGATCGACTTTTGGGATGAGCATGTGGCTGCTCAAATGGCTGCCTCTGTCTGTGGTGGACCGCGTCCTTCTGCTCGTCTCCAGGATCGTGCTCGGCGACACGGAGCAATGCGGCATAAGCAGGCCTCGGCTGGGGCCACTCGAGCTTAAGTCTCTCTCCGGCAAGACTCCTGTCCTCGATGTCGGAACCCTAGCCAAGATTCAGTCCGGTGACATTAAGGTTCGCCCAGGAATAAGTCGATTGACTAGAGATGGGGCTGAGTTTGTCGATGGAAGGATCGAAGATTTTGATGCAATCATATTGGCAACTGGTTACAAAAGCAATGTACTCTCTTGGTTAAAGGAACGTGAATTTTTCTCTGATAAGAGCGGATTGCCAAGGAGACCATTTCCCAACAGTTGGAAGGGCGAGCGAGGCCTCTACGCAGTTGGTTTCACGCAACGAGGCCTAATGGGGACTAAGATCGACTCGAGGAGGATTGTGCATGACATAGAGCAATGTTGGAAGGCCTCAAGGACTATGTAA

>Wp_YUC-P1

ATGGCCGGAAAGAGATTCCACGATCCTTCGCTCCAACTCCACGGCCACCGCCAATATCAACACCCCGCCGCCTCTGCGGCCGCCACTGACGATAAAGCTGAGCAGTTTATCTGGTTCCCGGGGCCTATTATTGTTGGCGCGGGGCCGTCGGGGCTAGCCGTCGCCGCTTGCCTGAAGGAGAAAGGTATCCCGTCGATGGTCATCGAGCGGTCCGACTGCATCGCCTCCCTCTGGCAGGTGAAGACCTACGATCGTCTCTCCCTCCATCTGCCGAAGCACTTCTGCGAGCTTCCGCTCATGCCATTCCCGGCCTCCTTCCCCCAATATCCCACGAAGCAGCAGTTCGTGGCCTATCTCGAGACCTATGCACGGCACTTCGACATACGACCGGTGTTCAATCAGACCGTCGTCAATGCTGAGTACGATGATCGGATGCGATCTTGGATGGTGCAGACAACCAAGACGGCGCCCACGGAGGGGAGGACGGCGACTAAGGGGTACATCAGCCGGTGGTTGGTGGTTGCTACCGGGGAGAATTCGGAGGAGGTGGTGCCAGAGATGGAAGGGATGACGGAGTTCAAAGGACCGATCATCCATACGAGTATGTTCAAGTCAGGCGAGGTGTTCGAGGGGAAGCCAGTGCTGGTGGTGGGCTGCGGCAATTCAGGGATGGAGATCAGCTTGGACCTTCACAACCACAATGCTCGTCCTCACCTCGTAGTAAGAGATACGGTGCACATATTGCCGAGAGAAATCATGGGCCGATCGACTTTTGGGATGAGCATGTGGCTGCTCAAATGGCTGCCTCTGTCTGTGGTGGACCGCGTCCTTCTGCTCGTCTCCAGGATCGTGCTCGGCGACACGGAGCAATGCGGCATAAGCAGGCCTCGGCTGGGGCCACTCGAGCTTAAGTCTCTCTCCGGCAAGACTCCTGTCCTCGATGTCGGAACCCTAGCCAAGATTCAGTCCGGTGACATTAAGGTTCGCCCAGGAATAAGTCGATTGACTAGAGATGGGGCTGAGTTTGTCGATGGAAGGATCGAAGATTTTGATGCAATCATATTGGCAACTGGTTACAAAAGCAATGTACTCTCTTGGTTAAAGGAAGGTGAATTTTTCTCTGACAAGAGCGGATTGCCAAGGAGACCATTTCCCAACAGTTGGAAGGGCGAGCGAGGCCTCTACGCAGTTGGTTTCACGCAACGAGGCCTAATGGGGACTAAGATCGACTCGAGGAGGATTGTGCATGACATAGAGCAATGTTGGAAGGCCTCAAGGACTATGTAA

>Wt_YUC-P1

ATGGCCGGAAAGAGATTCCACGATCCTTCGCTCCAACTCCACGGCCACCGCCAATATCAACACCCCGCCGCCTCTGCGGCCGCCGCTGACGATAAAGCTGAGCAGTTTATCTGGTTCCCGGGGCCTATCATTGTTGGCGCGGGGCCGTCGGGGCTAGCCGTCGCCGCCTGCCTGAAGGAGAAAGGTATCCCGTCGATGGTCATCGAGCGGTCCGACTGCATCGCCTCCCTCTGGCAGGTGAAGACCTACGATCGTCTCTCACTCCATCTGCCGAAGCACTTCTGCGAGCTTCCGCTCATGCCATTCCCGGCCTCCTTCCCCCAATATCCCACGAAGCAGCAGTTCGTGGCCTATCTCGAGACCTATGCACGGCACTTCGACATACGACCGGTGTTCAATCAGACCGTCGTCAATGCCGAGTACGATGATCGGATGCGATCTTGGATGGTGCAGACAACCAAGACGGCGCCCACGGAGGGGAGGACGGCGACTAAGGGGTACATCAGCCGGTGGTTGGTGGTTGCTACCGGGGAGAATTCGGAGGAGGTGGTGCCAGAGATGGAAGGGATGACGGAGTTCAAAGGACCGATCATCCATACGAGTATGTTCAAGTCAGGCGAGGTGTTCGAGGGGAAGCCAGTGCTGGTGGTGGGCTGCGGCAATTCAGGGATGGAGATCAGCTTGGACCTTCACAACCACAATGCTCGTCCTCACCTCGTAGTAAGAGATACGGTGCACATATTGCCGAGAGAAATCATGGGCCGATCGACTTTTGGGCTGAGCATGTGGCTGCTCAAATGGCTGCCTCTGTCTGTGGTGGACCGCATCCTTCTGCTCGTCTCCAGGATCGTGCTCGGCGACACGGAGCAATGCGGCATAAGCAGGCCTCGGCTGGGGCCACTCGAGCTTAAGTCTCTCTCCGGCAAGACTCCTGTCCTCGATGTCGGAACCCTAGCCAAGATTCAGTCCGGTGACATTAAGGTTCGCCCAGGAATAAGTCGATTGACTAGAGATGGGGCTGAGTTTGTCGATGGAAGGATCGAAGATTTTGATGCAATCATATTGGCAACTGGTTACAAAAGCAATGTACTCTCTTGGTTAAAGGTACGTGAATTTTTCTCTGATAAGAGCGGATTGCCAAGGAGACCATTTCCCAACAGTTGGAAGGGCGAGCGAGGCCTCTACGCAGTTGGTTTCACGCAACGAGGCCTAATGGGGACTAAGATCGACTCGAGGAGGATTGTGCATGACATAGAGCAATGTTGGAAGGCCTCAAGGACTATGTAA

>Wb_YUC-P2

ATGGCCAGAAAGAGATTCCGTGATCCTTCGCTCCAACTCAGCCACCGCCAATATAATCACCCCGCCACCGCCACTGCGGCCGCAGCTGTCGATAAAGCTGAGCAGTTCATCTGGTTTCCAGGGCCTATCATTGTAGGCGCGGGCCCTTCGGGGCTAGCCGTCGCCGCCTGCCTGAAGGAGAAGGGAATCCCGTCCATGGTCATCGAGCGCTCCGACTGCATCGCCTCCCTCTGGAAGCTGAAGACCTACGACCGTCTCTGCCTCCATCTGCCGAAGCACTTCTGCGAACTTCCACTCATGCCATTCCCCTCCTCTTTCCCCAGATATCCGACGAAGCAGCAGTTCGTGGCCTACCTCGAGACCTATGCACGGCACTTCGACATACAACCGATGTTCAATCAGACCGTCGTCAATGCCGAGTACGATGATTGGATGCAGTCGTGGAGGGTGAAGACGACGATCGAGGCGACGGAGGAGAAGGCGGCAACTAGGGAGTTCATTAGCCGATGGTTGGTGGTTGCTACCGGAGAGAATGCGGAGGAGGTGGTGCCGGAGATGGAAGGGATGACGGAGTTCAAAGGACGGATCATCCATACGAGCATGTTCAAGTCGGGCGAGGTGTTCGAGGGGAAGCGAGTGCTGGTGGTGGGCTGCGGCAATTCAGGGATGGAGATCAGCTTGGATCTTCACAACCACAATGCTCATCCTCACATCGTCGTAAGAGATACGGTGCACGTATTGCCCAGAGAAATCATGGGCCGATCGACTTTTGGGCTTTGTATGTGGCTACTCAAATGGCTGCCTTTGTCCGCTGTGGATCGCTTCCTTCTGCTTGTCTCCAGGATCGTACTCGGCGACACTGAAAAATGCAGAATAAGCAGGCCTCAGTTAGGGCCACTCGAGCTGAAGTCTCTCTCTGGCAAAACTCCGGTCCTCGACGTCGGAACCCTAGCCAAGATTCAGTCCGGCGACATTAAGGTTCGCCCAGGAATTAGCCGATTGACAAGTGACGGAGCTGAGTTTGTGGATGGAAGTATTGAAGATTTTGATGCAATCATATTGGCAACTGGTTACAAAAGTAGTGTTCTATCTTGGTTAAAGGAAGGAGAGTTTTTCTCTGATAAGAGTGGATTGCCGAAGAGACCATTCCCTAACAGTTGGAAGGGCAAGCGAGGCCTCTACGCGGTTGGTTTCACGCAACGAGGTCTAACAGGGACTAAGATCGACTCGAGGAGGATTGCGCATGACATAGAGCAATGTTGGAAGGCTCAAGGAGATGTTATCCATGAACCTCTCTCTAGGGTTCTTCCATGTCAAGATTGA

>Wm_YUC-P2

ATGGCCAGAAAGAGATTCCGCGATCCTTCACTCCAACTCCACAGCCACCGCCAATATAATCACCCCGCCGCCGCCACTGCGGCCGCAGCTGTCGATAAAGCTGAGCAGTTCATCTGGTTTCCAGGGCCTATCATTGTAGGCGCGGGCCCTTCGGGGCTAGCCGTTGCCGCCTGCCTGAAGGAGAAGGGAATCCCGTCCATGGTCATCGAGCGCTCCGACTGCATCGCCTCCCTCTGGAAGCTGAAGACCTACGAGCGTCTCTGCCTCCATCTGCCGAAGCACTTCTGCGAACTTCCACTCATGCCATTCCCCTCCTCTTTCCCCAGATATCCGACGAAGCAGCAGTTCGTGGCCTACCTCGAGACCTATGCACGGCACTTCGACATACAACCGATGTTCAATCAGACCGTCGTCAATGCCGAGTACGATGATTGGATGCAGTCGTGGAGGGTGAAGACGACGACCGAGGCGACGGAGGAGAAGACGGCAACTAGGGAGTTCATCAGCCGATGGTTGGTGGTTGCTACCGGAGAGAATGCGGAGGAGGTGGTGCCGGAGATGGAAGGGATGACGGAGTTCAAAGGACGGATCATCCATACGAGCATGTTCAAGTCGGGCGAGGTGTTCGAGGGGAAGCGAGTGCTGGTGTTGGGCTGCGGCAATTCAGGGATGGAGATCAGCTTGGATCTTCACAACCACAATGCTCATCCTCACATCGTCGTAAGAGATACGGTGCACGTATTGCCCAGAGAAATCATGGGCCGATCGACTTTTGGGCTTTGTATGTGGCTGCTCAAATGGCTGCCTTTGTCCGCTGTGGATCGCTTCCTTCTGCTTGTCTCCAGGATCGTACTCGGCGACACGGAAAAATGCGGAATAAGCAGGCCTCAGTTAGGGCCACTCGAGCTGAAGTCTCTCTCTGGCAAAACTCCGATCCTCGACGTCGGAACCCTAGCCAAGATTCAGTCCGGCGACATTAAGGTTCGCCCAGGAATTAGCCGATTGACAAGTGACGGAGCTGAGTTTGTGGATGGAAGTATTGAAGATTTTGATGCAATCATATTGGCAACTGGTTACAAAAGTAGTGTTCTATCTTGGTTAAAGGAAGGAGAGTTTTTCTCTGATAAGAGTGGATTGCCGAAGAGACCATTCCCTAACAGTTGGAAGGGCAAGCGAGGCCTCTACGCGGTTGGTTTCACGCAACGAGGTCTAACAGGGACTAAGATCGACTCGAGGAGGATTGCGCATGACATAGAGCAATGCTGGAAGGCTCAAGGAGATGTTATCCATGAACCTCTCTCTAGGGTTCTTCCATGTCAAGATTGA

>Wp_YUC-P2

ATGGCCAGAAAGAGATTCCGTGATCCTTCACTCCAACTCCACAGCCACCGCCAATATAATCACCCCGCCGCCGCCACTGCGGCCGCAGCTGTCGATAAAGCTGAGCAGTTCATCTGGTTTCCAGGGCCTATCATTGTAGGCGCGGGCCCTTCGGGGCTAGCCGTTGCCGCCTGCCTGAAGGAGAAGGGAATCCCGTCCATGGTCATCGAGCGCTCCGACTGCATCGCCTCCCTCTGGAAGCTGAAGACCTACGAGCGTCTCTGCCTCCATCTGCCGAAGCACTTCTGCGAACTTCCACTCATGCCATTCCCCTCCTCTTTCCCCAGATATCCGACGAAGCAGCAGTTCGTGGCCTACCTCGAGACCTATGCACGGCACTTCGACATACAACCGATGTTCAATCAGACCGTCGTCAATGCCGAGTACGATGATTGGATGCAGTCGTGGAGGGTGAAGACGACGACCGAGGCGACGGAGGAGAAGACGGCAACTAGGGAGTTCATCAGCCGATGGTTGGTGGTTGCTACCGGAGAGAATGCGGAGGAGGTGGTGCCGGAGATGGAAGGGATGACGGAGTTCAAAGGACGGATCATCCATACGAGCATGTTCAAGTCGGGCGAGGTGTTCGAGGGGAAGCGAGTGCTGGTGTTGGGCTGCGGCAATTCAGGGATGGAGATCAGCTTGGATCTTCACAACCACAATGCTCATCCTCACATCGTCGTAAGAGATACGGTGCACGTATTGCCCAGAGAAATCATGGGCCGATCGACTTTTGGGCTTTGTATGTGGCTGCTCAAATGGCTGCCTTTGTCCGCTGTGGATCGCTTCCTTCTGCTTGTCTCCAGGATCGTACTAGGCGACACGGAAAAATGCGGAATAAGCAGGCCTCAGTTAGGGCCACTCGAGCTGAAGTCTCTCTCTGGCAAAACTCCGATCCTCGACGTCGGAACCCTAGCCAAGATTCAGTCCGGCGACATTAAGGTTCGCCCAGGAATTAGCCGATTGACAAGTGACGGAGCTGAGTTTGTGGATGGAAGTATTGAAGATTTTGATGCAATCATATTGGCAACTGGTTACAAAAGTAGTGTTCTATCTTGGTTAAAGGAAGGAGAGTTTTTCTCTGATAAGAGTGGATTGCCGAAGAGACCATTCCCTAACAGTTGGAAGGGCAAGCGAGGCCTCTACGCGGTTGGTTTCACGCAACGAGGTCTAACAGGGACTAAGATCGACTCGAGGAGGATTGCGCATGACATAGAGCAATGCTGGAAGGCTCAAGGAGATGTTATCCATGAACCTCTCTCTAGGGTTCTTCCATGTCAAGATTGA

>Wt_YUC-P2

ATGGCCAGAAAGAGATTCCGTGATCCTTCGCTCCAACTCCACAGCCACCGCCAATATAATCACCCCGCCGCCGCCACTGCGGCTGCAGCTGTCGATAAAGCTGAGCAGTTCATCTGGTTTCCAGGGCCTATCATTGTAGGCGCGGGTCCTTCGGGGCTAGCCGTCGCCGCCTGCCTGAAGGAGAAGGGAATCCCGTCCATGGTCATCGAGCGCTCCGACTGCATCGCCTCCCTCTGGAAGCTGAAGACCTACGACCGTCTCTGCCTCCATCTGCCGAAGCACTTCTGCGAACTTCCACTCATGCCATTCCCCTCCTCTTTCCCCAGATATCCGAGGAAGCAGCAGTTCGTGGCCTACCTCGAGACCTATGCACGGCACTTCGACATACAACCGATGTTCAATCAGACCGTCGTCAATGCCGAGTACGATGATTGGATGCAGTCGTGGAGGGTGAAGACGACGATCGAGGCGACGGAGGAGAAGGCGGCAACTAGGGAGTTCATCAGCCGATGGTTGGTGGTTGCTACCGGAGAGAATGCGGAGGAGGTGGTGCCGGAGATGGAAGGGATGACGGAGTTCAAAGGACGGATCATCCATACGAGCATGTTCAAGTCGGGCGAGGTGTTCGAGGGGAAGCGAGTGCTGGTGGTGGGCTGCGGCAATTCAGGGATGGAGATCAGCTTGGATCTACACAACCACAATGCTCATCCTCACATCGTCGTAAGAGATACGGTGCACGTATTGCCCAGAGAAATCATGGGCCGATCGACTTTTGGGCTTTGTATGTGGCTGCTCAAATGGCTGCCTTTGTCCGCTGTGGATCGCTTCCTTCTGCTTGTCTCCAGGATCGTACTCGGCGACACGGAACAATGCGGAATAAGCAGGCCTCAGTTAGGGCCACTCGAGCTGAAGTCTCTCTCTGGCAAAACTCCGGTCCTCGACGTCGGAACCCTAGCCAAGATTCAGTCCGGCGACATTAAGGTTCGCCCAGGAATTAGCCGATTGACAAGTGACGGAGCTGAGTTTGTGGATGGAAGTATTGAAGATTTTGATGCAATCATATTGGCAACTGGTTACAAAAGTAGTGTTCTATCTTGGTTAAAGGAAGGAGAGTTTTTCTCTGATAAGAGTGGATTGCCGAAGAGACCATTCCCTAACAGTTGGAAGGGCAAGCGAGGCCTCTACGCGGTTGGTTTCACGCAACGAGGTCTAACAGGGACTAAGATCGACTCGTGGAGGATTGCGCATGACATAGAGCAATGTTGGAAGGCTCAAGGAGATGCTATCCATGAACCTCTCTCTAGGGTTCTTCCATGTCAAGATTGA

>Ba_Karkloof_YUC-P1

ATGGCCGGAAAGAGATTCCATGATCCTTCGCTCCAACTCCACAGCCATCTCCAATATCAACATCCCGCCGCCGCCCCTGCGGCTGCGGCTGTCGATAAAACTGAACAGTTCATCTGGTTCCCTGGCCCTATCATCGTCGGCGCAGGCCCGTCAGGGCTAGCAGTCGCCGCCTGCCTGAAGGAGAAAGGGATCCCCTCGATGATCATCGAGCGATCCGACTGCATCGCCTCCCTCTGGCAGCTGAAAACCTACGACCGTCTCTGCCTCCATCTGCCGAAGCACTTCTGCGAACTTCCGCTCATGCCATTTCCTGCCTCCTTCCCCAGATATCCCACCAAGCAGCAGTTCGTCGCCTACCTCGAGACATATGCACGCCACTTCGACATACGGCCGGTGTTCGATCAGACCGTCGTCAACGCCGAGTACGACGATCGGATGCAATTCTGGAGGGTACAGACAACGACGGCACCGACGGAGGAGAAGACGGCGAGCAAGGAGTACATGAGTCGGTGGTTGGTGGTGGCTACCGGAGAGAATGCGGAGGAGGTGGTGCCGGAGATGGAAGGGATGACGGAGTTCGAAGGACCGATCATCCATACTAGTATGTTCAAGTCCGGCGAGGTGTTCGAGGGAAAGCGAGTGCTGGTGGTGGGAAGCGGCAATTCAGGGATGGAGATCAGCTTGGACCTTCACGACCACAATGCTCGTCCTCACCTCGTCGTAAGAGATACGGTGCACATATTGCCTAGAGAAATCATGGGCAGATCGACTTTTGGCCTTTTCATGTGGCTGCTCAAATGGCTGCCTTTGTCGATGGTGGATCGCTTCCTTCTGCTCGTCTCCAGGATCGTTCTCGGCGACACCGAGCAATGCGGCATAAGCAGGCCTCACTTAGGCCCACTCGAGCTCAAGTCTCTCTCCGGCAAGACTCCGGTCCTGGACGTCGGAACCCTTGCCAAAATTCAGTCCGGTCACATTAAGGTTCGGCCAGGAGTAAGTCGATTGACTAGAGATGGAGCTGAGTTTGTAGATGGAAGGATAGAAGATTTTGATGCAATCATATTGGCAACTGGCTACAAAAGCAATGTACTCTCTTGGTTAAAGGAACGGGAATTTTTCTCCGATAAGAGTGGATTGCCGTGGAGACAATTCCCCAATAGTTGGAAGGGCGAGCGAGGCCTCTACGTTGTCGGTTTCACGCAACGAGGTCTAATGGGGACTAAGATCGACTCGAGGAGGATTGCACATGACATAGAGCAATGTTGGAAGGCTCAAGATTGA

>Ba_Ngeli_YUC-P1

ATGGCCGGAAAGAGATTCCATGATCCTTCGCTCCAACTCCACAGCCATCTCCAATATCAACATCCCGCCGCCGCCCCTGCGGCTGCGGCTGTCGATAAAACTGAACAGTTCATCTGGTTCCCTGGCCCTATCATCGTCGGCGCAGGCCCGTCAGGGCTAGCCGTCGCCGCCTGCCTGAAGGAGAAAGGGATCCCCTCGATGATCATCGAGCGATCCGACTGCATCGCCTCCCTCTGGCAGCTGAAAACCTACGACCGTCTCTGCCTCCATCTGCCGAAGCACTTCTGCGAGCTTCCGCTCATGCCATTCCCTGCCTCCTTCCCCAGATATCCCACCAAGCAGCAGTTCGTCGCCTACCTCGAGACATATGCACGCCACTTCGACATACGGCCGGTGTTCAATCAGACCGTCGTCAACGCCGAGTACGACGATCGGATGCAATTCTGGAGGGTACAGACAACGACGACGGAGCCGACGGAGGAGAAGACGGCGAGCAAGGAGTACATGAGTCGGTGGTTGGTGGTGGCTACTGGAGAGAATGCGGAGGAGGTGGTGCCGGAGATGGAAGGGATGACGGAGTTCGAAGGACCGATCATCCATACTAGTATGTTCAAGTCCGGCGAGGTGTTCGAGGGAAAGCGAGTGCTGGTGGTGGGAAGTGGCAATTCAGGGATGGAGATCAGCTTGGACCTTCACAACCACAATGCTCGTCCTCACCTCGTCGTAAGAGATACGGTGCACATATTGCCTAGAGAAATCATGGGCAGATCGACTTTTGGCCTTTTCATGTGGCTGCTCAAATGGCTGCCTTTGTCAATGGTGGATCGCTTCCTTCTGCTCGTCTCCAGGATCGTTCTCGGCGACACCGAGCAATGCGGCATAAGCAGGCCTCACTTAGGCCCACTCCAGCTCAAGTCTCTCTCCGGCAAGACTCCGGTCCTGGACGTCGGAACCCTTGCCAAAATTCAGTCCGGTCACATTAAGGTTCGGCCAGGAGTAAGTCGATTGACTAGAGATGGAGCTGAGTTTGTAGATGGAAGGATAGAAGATTTTGATGCAATCATATTGGCAACTGGCTACAAAAGCAATGTACTCTCTTGGTTAAAGGAACGGGAATTTTTCTCCGATAAGAGTGGATTGCCGTGGAGACAATTCCCCAATAGTTGGAAGGGCGAGCGAGGCCTCTACGTTGTCGGTTTCACGCAACGAGGTCTAATGGGGACTAAGATCGACTCGAGGAGGATTGCGCATGACATAGAGCAATGCTGGAAGGCTCAAGATTGA

>Di_YUC

ATGGAGGGAAAAAGATCTCATGATCCCTTGTTTCGCCACCATCAACATCATCAATCCGGTGATACTTATTATAGAGTGGCGAAAGATACCGCCGGCGCCGTTGATAAAGCCGAGCAGTTAATCTGGTTCCCCGGCCCGATCATCGTCGGCGCAGGGCCGTCGGGGCTAGCCGTCGCCGCCTGCCTCAAGGACAAAGGGATACAGGCGCTGGTCATCGAACGGTCCGACTGCATCGCCTCCCTGTGGCAGCTCAAAACCTACGACCGTCTCTGCCTCCATCTGCCGAAGCATTTCTGCGAGCTTCCTCTCATGCCATTCCCCGCCGTCTTCCCCGGATACCCTACGAAGCAACAGTTCGTGACCTACCTGGAGGCCTATGCCCGGAAGTTCGACATACGGCCAGTGTTCAATCAGACAGTCGTGGGAGCCGAGTACGACGGACGGATGCAGTCGTGGAGAGTGAAGACTTCGGAGCAGAAGGACGGGAAGACGACGGAGTACGTCTGCCGTTGGTTGGTGGTGGCTACCGGAGAGAATGGGGAGGAGGCGGTGCCGAATATCGAGGGCATGGCGGACTTCGAGGGCCCGATCGTCCATACGAGCTTGTACAAGTCCGGTGAGGTGTTCCAGGGGAAGAGGGTGCTGGTGGTGGGCTGCGGGAATTCAGGAATGGAGATCAGCTTGGACCTCCACGACCACAATGCTCACCCTCACCTCGTCGTAAGAGATGCGGTACACATATTGCCCAGGGAAATCTTGGGCCGATCGACTTTTGGGCTCTGCATGTCGCTCCTCAAATGGCTACCCTTGTCCGCGGTAGACCGCTTCCTGTTGCTCGTCTCCAGGATCGTACTCGGCGACACGGAGCAATGCGGCATAACCAGGCCCCAGTTAGGGCCTCTCGAGCTCAAGTCCATCTCCGGCAAGACGCCGGTCCTCGATGTCGGAACCCTAGCCAAGATCCAGTCCGGTGACATCAAGGTTCGCCCAGGAATAAATCGATTGACAAGACATGGAGCTGAATTTGTGAATGGGAGGGTCGAGGATTTTGATGCGATAATCTTGGCAACTGGTTACAAAAGCAATGTGCTCTCTTGGTTGAAGGAGCGAGAGTTCTTCTCAGACAAGAATGGGTTGCCGAGGAGGCCATTTCCCCACAGTTGGAAGGGTAAGCGAGGCCTCTACGCCGTCGGTTTCACGCAGCGAGGCTTAATGGGGACTTCGGTCGATGCAAGGAGGATTGCGCACGACATCGAGCAATGCTGGAAGGCTGAAACAAATTAA

>At_YUC2

ATGGAGTTTGTTACAGAAACGTTAGGCAAGAGAATCCATGATCCGTACGTGGAGGAAACTAGGTGCTTAATGATTCCCGGACCAATCATTGTCGGTTCCGGGCCGTCGGGACTGGCCACAGCGGCATGTTTAAAGTCGAGAGACATCCCTAGTTTGATTCTAGAACGTTCCACTTGCATAGCGTCACTATGGCAGCACAAAACATATGATCGTCTTCGACTTCATCTCCCTAAAGATTTCTGTGAGCTTCCATTGATGCCTTTTCCTTCAAGCTACCCTACTTACCCTACAAAGCAACAGTTCGTCCAATACCTTGAGTCTTACGCCGAACATTTTGACCTAAAGCCCGTTTTTAACCAGACCGTGGAGGAAGCCAAGTTCGATAGGCGGTGTGGGTTATGGAGGGTGAGGACAACCGGAGGGAAGAAGGATGAGACAATGGAGTATGTATCACGGTGGCTTGTTGTGGCGACCGGGGAGAATGCCGAGGAGGTGATGCCGGAGATTGATGGAATCCCGGATTTTGGTGGACCTATCCTCCACACAAGCTCCTATAAGAGCGGTGAAATATTTAGTGAGAAGAAGATTTTGGTTGTAGGATGTGGAAACTCCGGGATGGAAGTTTGTTTAGACCTTTGCAACTTCAATGCTCTTCCTTCTCTTGTGGTTCGTGACTCGGTACACGTATTACCTCAAGAAATGCTAGGTATATCGACTTTCGGGATATCCACGAGCCTGCTCAAGTGGTTTCCAGTGCACGTGGTGGACCGGTTCTTGTTACGTATGTCTCGGTTGGTTCTTGGTGACACGGATCGGTTAGGGTTAGTTCGACCAAAACTTGGCCCTCTTGAACGCAAGATCAAATGCGGAAAGACTCCTGTTTTGGACGTTGGCACTCTTGCCAAAATCCGAAGTGGACACATCAAGGTGTATCCGGAGTTGAAACGGGTAATGCATTATTCGGCAGAGTTTGTTGATGGGAGAGTAGATAACTTCGACGCCATTATACTCGCCACGGGTTACAAAAGCAACGTACCCATGTGGCTAAAGGGAGTGAACATGTTTTCTGAGAAAGATGGATTTCCGCATAAACCATTTCCTAACGGTTGGAAAGGCGAAAGCGGATTGTATGCAGTCGGTTTCACAAAGCTTGGATTGCTTGGTGCAGCCATTGATGCCAAGAAGATCGCTGAGGATATTGAGGTTCAACGACATTTCTTACCATTGGCTCGTCCTCAACATTGTTAA

>At_YUC6

ATGGATTTCTGTTGGAAGAGAGAGATGGAAGGTAAACTAGCACATGACCACCGCGGCATGACGTCACCGCGTCGTATCTGCGTCGTCACCGGTCCGGTGATCGTAGGCGCCGGACCGTCGGGACTAGCCACGGCAGCATGTTTAAAAGAGAGAGGTATCACGTCCGTACTACTAGAGAGATCAAACTGTATAGCATCACTATGGCAGCTCAAGACTTATGACCGTCTTCATCTTCACCTTCCTAAACAATTCTGTGAACTTCCGATTATACCCTTCCCCGGAGATTTCCCTACCTACCCGACGAAGCAACAGTTCATCGAGTACCTTGAGGACTACGCTCGGAGGTTTGACATAAAGCCGGAGTTTAACCAAACGGTTGAGTCGGCTGCGTTTGATGAAAACCTTGGGATGTGGCGCGTGACTAGCGTGGGAGAAGAAGGCACGACGGAGTATGTTTGTCGGTGGTTAGTGGCGGCGACGGGGGAGAATGCGGAGCCGGTGGTACCTAGGTTTGAGGGGATGGATAAGTTTGCAGCCGCCGGGGTAGTTAAGCACACGTGTCATTATAAAACCGGTGGAGATTTCGCCGGAAAAAGGGTTCTTGTCGTCGGATGTGGAAACTCCGGTATGGAGGTTTGTTTGGATCTCTGCAACTTCGGTGCTCAGCCTTCTCTCGTTGTCAGAGACGCTGTGCACGTCCTACCACGAGAGATGTTGGGTACTTCAACTTTTGGGCTGTCCATGTTCTTACTGAAATGGCTGCCCATCCGGCTTGTTGACCGTTTCCTTTTGGTTGTTTCCCGGTTCATCCTCGGGGATACTACCCTTTTAGGTCTTAACAGGCCCCGGTTAGGTCCACTCGAGCTCAAAAATATCTCCGGTAAAACTCCGGTTCTCGACGTTGGCACGCTAGCCAAAATCAAAACCGGAGACATTAAGGTGTGTTCGGGGATAAGAAGGTTAAAACGACATGAAGTTGAGTTCGATAACGGAAAAACAGAGAGATTTGACGCCATTATATTAGCAACTGGCTACAAAAGCAACGTACCCTCTTGGCTAAAGGAGAATAAAATGTTTAGTAAGAAAGATGGATTTCCAATACAAGAGTTCCCTGAGGGATGGAGAGGGGAATGTGGGCTATACGCGGTCGGATTCACAAAACGTGGGATTAGTGGAGCATCAATGGATGCAAAGAGAATAGCTGAAGACATACACAAGTGTTGGAAACAAGACGAGCAACTGCAATGCAAATTGGGGAAAAGAATGAAAAGGAAATTTAGTGAGAGTGATTGTGGTGGGAATTGA

>Os_YUC2_XP_025876562.1

ATGGACCCTTGGAGTGAAATTGAGGGCAAGAGAGCCCATGATCCTATCTTCCAAAACTACTTCAGCCAAA

ACTGCCGCCAATCTGTTGATGGTTTCTGCAAGAAGAGGAGCGCAGATGCTGCCGTCGCTCGCGCCGAGCG

ATGCATCCGGGTTCTGGGGCCAATCATCGTGGGTGCTGGACCATCAGGGCTCGCTGTTGCTGCATGTCTC

AAGGAGAAGGGAGTTGACAGTCTTGTTCTCGAGCGCTCCAACTGCATAGCTTCCCTTTGGCAGCTGAAGA

CATACGATCGTCTCAGCCTTCATCTTCCTCGCCAATTCTGTGAGCTTCCCCTCATGCCTTTTCCTGCCTA

CTACCCTATTTATCCCTCAAAGCAGCAGTTTGTAGCCTACCTGGAGAGCTACGCTGCAAGGTTTGGGATC

TGCCCCACGTACAACCGGACGGTGGTGTGTGCAGAATATGATGAGCAGCTTCAGTTATGGCGGGTGAGGA

CACGGGCCACCGGCATAATGGGAGAGGAGGTCGAGTATGTGTCTCGGTGGTTGGTTGTGGCCACCGGTGA

GAACGCCGAGGTTGTGCTGCCAGAGATTGATGGCCTAGACGACTTCAAGGGAACTGTTATGCACACCAGT

TCATATAAAAGTGGCGGTGCATTTGCTGGGAAGCGTGTTCTCGTTGTTGGGAGTGGCAACTCCGGCATGG

AGGTGTGCCTAGACCTCTGCAACCACAATGCAAATCCCCATATTGTAGTAAGAGACGCTGTACACATCTT

GCCCAGGGAGATGCTGGGTCAGTCCACCTTTGGGCTGTCAATGTGGCTGCTCAAGTGGCTCCCAGTCCAC

GTGGTGGACCGAATTCTACTGCTCATAGCTCAGACCATGCTTGGGGATACTGCTCAGCTTGGGCTAAAGC

GTCCTACCATCGGTCCTCTCGAGCTCAAGTCACTCTCAGGGAAGACCCCAGTTCTTGACGTCGGCACATT

TGCAAAGATTAAGTCTGGTGACATCAAGGTACGGCCGGCCATAAAACAAATATCAGGGAGACAGGTAGAG

TTCATGGACACAAGGTTGGAGGAGTTTGATGTCATTGTGCTTGCCACCGGCTACAAGAGCAACGTTCCCT

TCTGGTTAAAGGACCGGGAGTTGTTTTCCGAAAAGGATGGGTTGCCAAGGAAGGCATTTCCAAACGGTTG

GAAGGGTGAGAACGGGCTCTACTCGGTCGGGTTCACCCGGCGCGGGCTGATGGGAACATCGGTGGATGCT

CGGAGAATTGCTCACGACATTGAGCAGCAATGGAAGGCCAGAGGGAAGCACCCGGGCGTGTTGCTCTAG

>Os_YUC2_XP_015638961.1

ATGTGTTGTTCCCACCAGCTTGTTTGGGTTCAAGGGCCAATAGTTGTTGGCGCAGGGCCATCTGGACTCG

CTGCTGCTGCATGCCTGAAGGAGAAGGGAATCGACAGCCTTGTTCTTGAGCGCTCGAGCTGCTTAGCTCC

TCTATGGCAGCTCAAGATGTATGACCGCCTCAGCCTTCATCTACCTCGTCAGTTCTGTGAACTTCCTCTC

TTTCCTTTCCCTGCCAGCTACCCTGATTACCCCACAAAGCAGCAGTTTGTGGCTTACCTGGAGAGCTATG

CTGCAAAGTTTGGTATCAATCCTATGTACAACCATACAGTGGTGTGCGCAGAATTTGACGAGCGACTGAT

GCTATGGCGGGTGAGGACTACACAGGCCACTGGCATGATGGAAGATGATGTTGAGTATGTGTCGCAGTGG

CTGGTTGTTGCAACCGGAGAGAATTCAGAGGCCGTGCTGCCAGTGATCGATGGCTTGGAAGAGTTTCGAG

GAAGTGTCATCCACACTAGCGCATACAAGAGCGGTTCTAAGTTCGCTGGGAAGACTGTCCTTGTTGTAGG

GTGTGGCAACTCTGGCATGGAGGTGTGCCTAGACCTGTGCAACCACAATGGTTACCCTCGTATTGTAGTA

AGAGATGCAGTGCACATCTTGCCCAGGGAGATGCTAGGTCAGCCGACCTTCCGGCTTGCAATGTGGCTGC

TCAAATGGTTGCCGATCCATATCGTAGACCGGATTTTACTGCTTGTTGCACGAGCGATTCTTGGTGATAC

ATCACAATTTGGGTTGAAAAGGCCTAGTCTTGGGCCACTTGAGCTGAAATCACTGTCAGGAAAGACACCC

ATCCTTGACATTGGCACTCTTGCAAAGATCAAGTCTGGAGATATAAAGGTCCGACCTGCCATAAGAAGAA

TTGCAGGGCAGCAAGTAAAATTTGTGGATGGTCGTTCAGAGCAGTTTGACGCCATTGTACTTGCCACCGG

CTACAAAAGCAACGTTCCCTGCTGGTTAAAGGACCAAGGGCTGTTTTCAGAAAAAGATGGGTTGCCAAGG

AAAGCGTTTCCAAACGGGTGGAAGGGCGAGAGGGGTCTATACTCCGTGGGATTCTCCCGTCGTGGCCTGA

TGGGAACGGCTGCCGATGCGAGAAGGATTGCTCATGACATTCATATGCAATGGAAGTCGTCCAAAGGGAG

GAGTCGTCCAGCCAAACCTTCTCCTTAG

>Ma_YUC2

ATGGAGCGTTGGAGGGAAGCAGAAGGTAAAACGCTCCACGATCCCTTGTTCCATCTCTGCCCCCCCCTTG

CCTTCCATCAACCGAGCGATGGCTTCCATGAGGTGGCGACCAGAGATGAAGCCGAGCGCAGCATCTGGGT

TCCCGGGTCGATCATCGTTGGCGCTGGTCCGTCGGGTCTCGCGGTGGCCGCGTGCCTCGAGGCGAAGGGC

GTCCCCAGCATGATCCTGGAGAGGTCCAACTGCATCGCCTCTCTCTGGCAGCTCAAGACGTACGACCGCC

TCCGTCTCCACCTGCCCAAGCGTTTCTGCCAGCTCCCCCTCGCCCCCTTCCCCGCCTGCTTCCCCACGTA

TCCCACCAAGCAGCAGTTCGTGGCCTACCTCGAGGCCTACGCCCGGCGGTTCCACATCCGCCCCTGCTTC

AACCAGACGGTGGTGAGCGCCGAGTACGACGGCCGCGTCAAGCTATGGCGAGTGCGGGCGGTCAGGGCCG

GGAACGACAAGGCGGCCGAGTACGTCTCCCGGTGGCTGGTGGTCGCCTCCGGGGAGAATGCTGAGGCGGC

CGTGCCCGACATCGATGGCATGTCGATATTTAAGGGCCCCATCATCCACACCAGCTCGTACAAGAGCGGC

GATGAGTTCCAAGGCAAGCGGGTCCTCGTGATCGGTTGCGGCAATTCCGGCATGGAGGTCTGCTTGGACC

TCTGCAATCACAGTGTCCGTCCTCGCATCGTCGTAAGAGAATCGGTGCACATCCTGCCGAGGGAGATGCT

GGGGCGGTCCACCTTTGGGCTCTGCATGTGGCTCCTAAAGTGGCTCCCCATGCGCACCGTGGACCGCATC

CTCTTGCTCGTCTCCAGGGTCATGCTGGGCGACACCCAGCGATACGGCCTCCGGCGGCCCCGGTTGGGTC

CCCTCGAGCTCAAGTCGCTCTCCGGGAAGACGCCGGTCCTCGATGTCGGGGCCCTGGCCAAGATCAAGTC

TGGCGACATCAAGGTTTGCCCCGCCGTAAAGCGACTGACAGGGCATGGAGCAGAGTTCGTGGATGGCAGG

TCCGAAGACTTCGATGCGATTATCCTAGCGACCGGCTACAAGAGCAACGTGGCGTCTTGGCTGAAGGAGA

GGGAGTTCTTCTCGGATAAGGATGGGTTTCCGAGGAAGGTGTTTCCCGATAGCTGGAAGGGAGAGCAGGG

CCTGTACGCGGTGGGGTTCACGCGACAGGGCTTGATGGGGACTTCCGTGGACGCCAAGAGGATAGCTCAT

GACATCAAACAGTGTTGGATGGCCGAACCAAAGCAACGCATGCTGCCATCACAAACTTGA
