## Supplementary material for "Supergene control of chiral development in mirror-image flowers": Data S4

Insertions are indicated with green highlight.

Breakpoints of deletion are indicated with yellow highlight.

>RH2 Wm right homostyle

TTTCAAACATGGAGCAGATGGATCTAAAAGATATAAGGCCATGTTAGTTGTCAAAGGATTTCAACAAAAGGCAGGTATAGACTTTACCGATATCTTTTCCCCTGTAATGAAAATGTCTACGGTAAGATATGTTCTAGGTTTAGTTGTTGCTGAAGATTTGCATTTAGAACAGATGGATGTTAAGACAGCTTTTCTTCATGGAGATTTGGAGGAAGATTTATGT|ATCTTTTATCAATTGCATCGCATGCAACTCTTGGATAATTAGATGAACAACTCTTGTTATTAACAACTCATCTCCTTGCATCTAG

>RH1 Wm right homostyle

CATGCTAATATGCCTTGGATGATCCACATCTTTGGGTCAGGCGCTGATGAGCACATTGAACTATCCATGTGGCTGAGAGCCCTCACAGCCCAAGCACAACGACGCAATTATCAGGGGCGAGCTCTACCACTGAGCTAATAGCCGTCTTGTGGGCCTCCCAGTGGGAGGCTTTCTATGCCAAAAGCGAGAAAAACCCACCCCTCTCTTTCCTTTTTGCGCCCCCATGTC|CCTCTTTGGCAATCCATTTTCATCTGAGAAGAACTCTGGATCCTGCATTTGATGAATTTATTATTAACTATGGCAAATTAGATTCACACTTTAACAAGAAAATGGCTTGTGTTACATGATTTCATTATATACCGCGAATATCAAAATCATTGTAATCACCTTTCCTCTTCGTACTAGGAATTGCCAATTAAATGATACTTTTTGGAAAAGGACTATCAAGAAGTACAGTTCATG

>F215 Wp right homostyle

GGCAATTTCCGAACCCGACCCGAAATTAGGTTCGGATCTGAACATGTTATCCGAACCCGACCCGAACCCGAAAGCTCGGGTACCCGACCCGAAATATTTGCTCGGGTTCGGGTCAGAAATTCGGGTATCGGGTATCCGTTAGCACCTTTAGATACAATCAGTCGGAAGTATGTCGTAGCG|ACTTTAACAAGAAAATGGCTTGTGTTACATGATTTCATTATATACCGCGAATATCAAAATCATTGTAATCACCTTTCCTCTTCGTACTAGGAATTGCCAATTAAATGATACTTTTTGGAAAAGAACTATCAAGAAGTACAGTTCAATGGAAAAGC

>GH Wp right homostyle

TCGAATATATCTCAGATATTTCGAGTCGATGTATAATAAGATGTAGCTAATCTCTACAACCCTAAGTAAAATCGGATAGGTACCCCTAGTTCAAATCCTGTCTCTCATCTCGGAATTTCCCAAAAGCCGAGGTGATCATTTCGTGTAAGTGGGTAGATCTTTATGCCTGATGACACTATGAAGTATGAAAGCATCTAAATTCAGAACAAATGTACAAAACATTTCAAAAATAATACACCATAACCTACAGAGTCCAAGACTCCTTACATAAACCTAAAAGATATTCCTAGGTAAAGACTTCATAAACGGATCAGCCACCATCATAGTCAGTATGTGTTGGAGAACCACTTCCCACTCCTCGATTGTCTCTCT|TCATGATGCTGCTGATGATCGC

>Ron. Wb left homostyle

ATGCTTAATCATGTTTTTTTAATCAGGACTATTCAAAACAATTTAAATCTCTAAGGAGTAACTGTAGAAGGACATTCATAATCATGGATTTTTCGCGTCAAAATGTTGATCAAGACTTATTAAGTAATACTTATCAAAATTAATTGACAAAATCCAAGTTTGGAAAAAATCCCTCAAAAGAAAAAAATGATGAAAAAATAAAAGAAGAAAACATAAATTGGCATTTTTTTTCTTTTTCACTTGTAGAATATATTTGCATATATTAGTCTAACTTACATAAATATGAGAATTTCTTTAAAGTACATTGAGATTTACTAGGATCCAAAGTATATATTAGTTTATGTATTCTAACTCTATTTTTTGGCTAACATTCTAAACATTTTATCTTTCAAATAAAGAGACAACTAAGTCCAATTTTAAAATTAAGCATATTTACAATTTAGTTAATGTGCTTTCTTTTCTTTCTTTTTGATTGATGCGTAATTTTGGGGGTTTTGAAAACATGATAAAGTGACTATTTTTCCAACTTTGATCTTATCCTGCATGATCCTCAAGTGTTTATTCATTT|AATGCTAAGGGTCCATTATGTA

>F215 Wb left homostyle

TCCTTTGGGGTATTTTTATCTGTCTGTTCGTTTTTGTTCTGGTGTTTTTCTACAGCCTTCTAAGTTTACCTGATTATGCAGTTCGCTCTCTAGCACTTACTTTTACTGTTTCACATACTTCAATAGATTTGTAGTTTATTGTCTTAGA|AGAAAATCACTAGAGGGGAATCCTAGTTTCTAGTGGACATAAGTATTAGTTTATTTTCTTGTAAAAAAAAAATTCTTATAATTCAAAAATCAAGTTTATAGTATATAAGAATATTTGAAAGTAGTAATAATATTTAAATTTATTTTTCACATTAATAAAATTAACATTTTTGATAGTTTCTAATCAAGATCATAAGATTGCATATTATCACTTTTATTGTGTTTAATTTGAATTAGATGTTATCTATTATAAATTAGTTGAGAGTTCTTGGATTTC
